# Cross-etiology transcriptomic conservation in hepatocellular carcinoma reveals opposing proliferation and hepatocyte-loss programs validated across cohorts

**DOI:** 10.64898/2026.03.10.710938

**Authors:** Ricardo Romero, Cinthia C. Toledo

**Affiliations:** Departamento de Ciencias Naturales, Universidad Autónoma Metropolitana, Unidad Cuajimalpa. Avenida Vasco de Quiroga 4871, Col. Santa Fe Cuajimalpa. Alcaldía Cuajimalpa de Morelos, C.P. 05348, Ciudad de México. México

**Keywords:** hepatocellular carcinoma, viral hepatitis, GSVA/ssGSEA, transcriptomic meta-analysis, gene module score

## Abstract

**Background:** Hepatocellular carcinoma (HCC) arises from diverse etiologies, but the extent to which viral etiologies converge on reproducible transcriptomic state axes remains incompletely resolved.

**Methods:** We analyzed HBV- and HCV-associated HCC discovery cohorts using Hallmark GSVA, limma-based differential modeling, and cross-cohort meta-analysis. Conserved tumor-upregulated and tumor-downregulated genes were distilled into ProlifHub and HepLoss modules, combined as HCCStateScore = ProlifHubScore - HepLossScore. Module performance was evaluated across multiple independent GEO cohorts, module-size robustness was tested across alternative top-N definitions, and TCGA-LIHC was used for continuous Cox survival modeling. An HBV-derived injury axis was constructed from an ordinal ALT/AST/HBV-DNA injury index in GSE83148 and tested in GSE121248 with adjustment for E2F/G2M activity and CIBERSORTx-inferred immune composition.

**Results:** HBV- and HCV-associated HCC showed conserved activation of proliferation/repair programs and suppression of hepatocyte functional programs. The HCCStateScore validated across independent HCC cohorts with consistently positive tumor-non-tumor deltas and high discrimination, and module-size sensitivity analysis showed that performance was not dependent on the top-20 cutoff. In TCGA-LIHC, higher ProlifHubScore and HCCStateScore were associated with poorer overall survival in continuous Cox models, including after age/sex/stage adjustment. A compact HBV injury program remained tumor-associated after simultaneous adjustment for E2F/G2M activity and CIBERSORTx-derived immune-composition covariates, with concordant results using an extended FDR-defined injury set.

**Conclusions:** HCC exhibits a robust cross-etiology transcriptomic state characterized by opposing proliferation and hepatocyte-loss programs. The module framework provides a portable bulk transcriptomic state score and supports a residual tumor-associated HBV injury component that is not fully explained by proliferation or inferred immune composition.

## Introduction

Viral hepatitis remains a major cause of liver-related morbidity and mortality worldwide, and it is a dominant upstream driver of hepatocellular carcinoma (HCC) [1]. HCC is the predominant primary liver malignancy and a major contributor to global cancer mortality, with a large fraction of cases arising in the setting of chronic hepatitis B virus (HBV) or hepatitis C virus (HCV) infection, often through progression to fibrosis and cirrhosis [2]. Despite major advances in prevention and therapy, such as HBV vaccination and nucleos(t)ide analog therapy, and curative direct-acting antivirals for HCV, viral hepatitis–associated liver cancer continues to impose a substantial clinical burden.

HBV and HCV differ in their virology and in how they promote malignant transformation. HBV is a DNA virus that persists through covalently closed circular DNA and can integrate into the host genome, enabling long-term antigenic stimulation and potentially contributing to genomic instability [3,4]. By contrast, HCV is an RNA virus that replicates in the cytoplasm; its oncogenic effects are thought to arise primarily from chronic inflammation, oxidative stress, and repeated cycles of hepatocyte death and compensatory regeneration [3]. Yet both etiologies converge on a final common pathway of chronic injury, dysregulated wound-healing, and selective pressures that favor proliferative, stress-tolerant cell states.

Transcriptomic profiling has repeatedly shown that HCC tumors share recurrent biological themes, including reactivation of cell-cycle programs (e.g., E2F targets and G2M checkpoint), replicative stress responses, and remodeling of DNA repair and stress pathways, alongside broad loss of hepatocyte differentiation and metabolic functions [5]. Yet published gene-expression signatures often vary across cohorts and platforms and can be sensitive to confounding from adjacent-tissue composition, inflammatory infiltrates, and technical batch effects. This motivates pathway- and module-level analyses that (i) compress gene-level variability into interpretable programs and (ii) emphasize signals that replicate across independent cohorts and etiologies.

Pathway-level scoring provides a practical route to cross-cohort comparability because it aggregates signal across sets of functionally related genes. Gene Set Variation Analysis (GSVA) enables sample-wise estimation of gene-set activity from bulk expression data without requiring a priori phenotype labels, allowing both discrete contrasts (tumor vs adjacent) and continuous axes to be modeled within a unified framework [6]. In parallel, the Hallmark collection in MSigDB offers a curated, low-redundancy set of gene programs that summarize major oncogenic and tissue-state processes, supporting interpretable comparisons across studies while reducing redundancy in pathway testing [7].

Beyond conserved proliferative programs, hepatitis-related injury and regeneration may represent a mechanistic bridge between chronic infection and malignant state, particularly in HBV where immune-mediated tissue damage can persist even under partial viral control. Prior liver transcriptome studies in HBV-associated acute liver failure have described regeneration- and inflammation-linked signatures consistent with an injury-to-repair continuum [8]. Motivated by this literature and by the availability of clinical biomarkers in hepatitis cohorts, we treat hepatitis injury as a quantitative axis and test whether it remains elevated in HBV-associated tumors after accounting for dominant proliferative programs.

Here, we use public HCC transcriptomic cohorts to separate a conserved bulk tumor-state axis from etiology-linked residual signals. We first quantify Hallmark pathway activity in HBV- and HCV-associated HCC discovery cohorts and use gene-level meta-analysis to construct compact proliferation and hepatocyte-loss modules. We then test whether the resulting HCCStateScore generalizes across independent GEO [9] cohorts and whether its prognostic coherence is supported in TCGA-LIHC using continuous Cox models [10]. Finally, we derive an HBV injury program from a non-tumor hepatitis cohort and evaluate whether its elevation in HBV-HCC persists after adjustment for dominant proliferative activity and inferred immune-cell composition. This design does not claim that proliferation-versus-hepatocyte-loss opposition is novel in HCC; rather, it tests whether this established biological axis can be converted into a reproducible cross-cohort state score and used to control for dominant tumor-state variation when evaluating HBV-linked residual injury. An overview of the multi-cohort design and analysis workflow is provided in Fig. 1.

**Figure 1.**
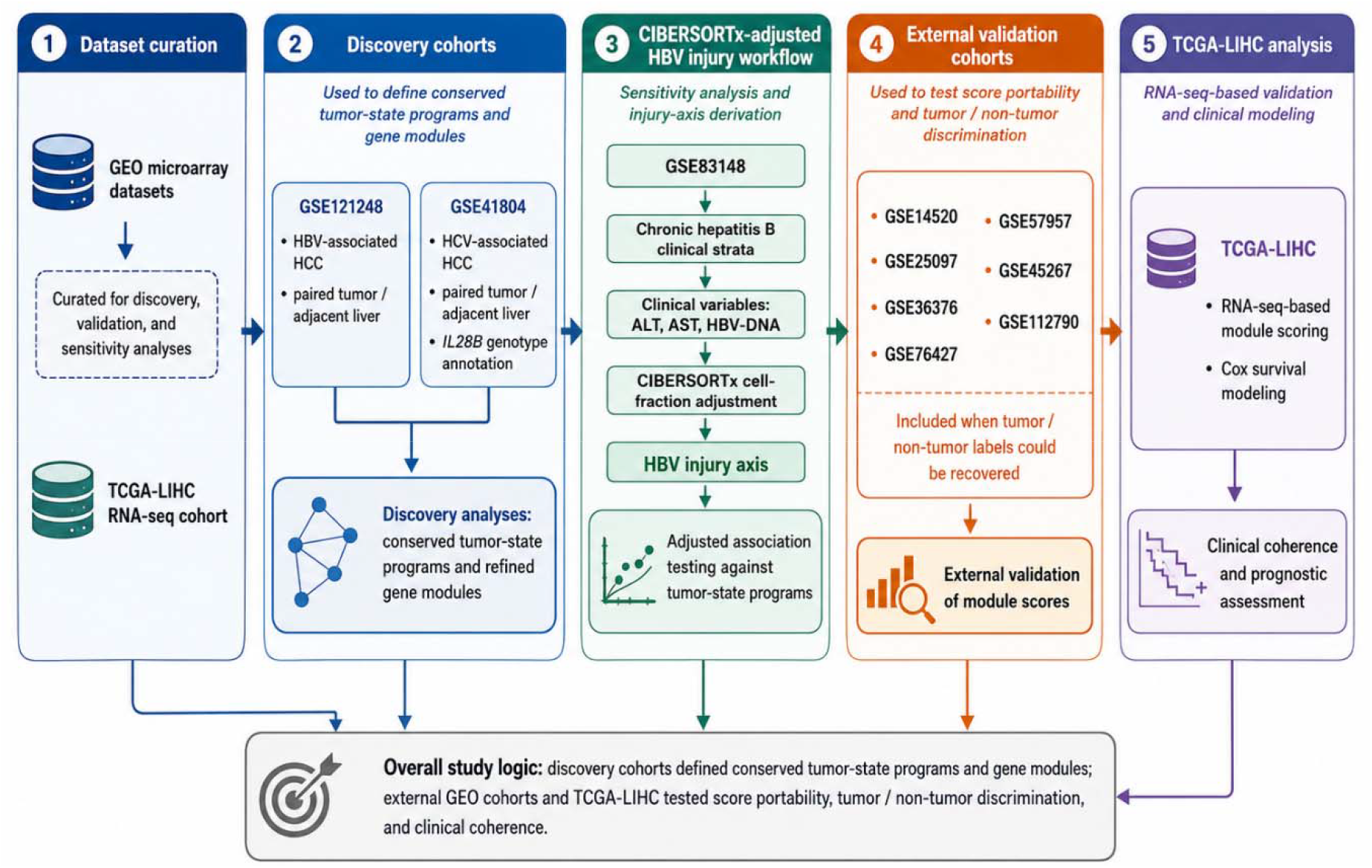
Study design and analysis workflow. We first curated GEO and TCGA-LIHC datasets to support discovery, validation, and sensitivity analyses. Two paired viral-HCC cohorts were retained as discovery datasets: GSE121248 for HBV-associated HCC and GSE41804 for HCV-associated HCC with IL28B genotype annotation. GSE83148 was used to derive an HBV injury axis from chronic hepatitis B clinical strata, including ALT, AST, and HBV-DNA. Additional GEO HCC tumor/non-tumor cohorts were curated for external validation, including GSE14520, GSE25097, GSE36376, GSE76427, GSE57957, GSE45267, and GSE112790 when tumor/non-tumor labels could be recovered. TCGA-LIHC was used for RNA-seq-based module scoring and Cox survival modeling. This expanded structure directly separated discovery from validation. The discovery cohorts were used to define conserved tumor-state programs and gene modules, whereas the external GEO cohorts and TCGA-LIHC were used to test score portability, tumor/non-tumor discrimination, and clinical coherence.

## Materials and Methods

### Study design, dataset selection, and cohort structure

We conducted a multi-cohort transcriptomic analysis of hepatocellular carcinoma (HCC) designed to identify conserved tumor-state programs across viral etiologies and to test whether an HBV-associated injury axis persists after adjustment for proliferation and inferred immune-cell composition. The analysis was organized into five components: discovery pathway analysis, gene-level meta-analysis, module construction, external validation, and adjusted modeling of the HBV injury axis.

Two paired HCC cohorts were used as discovery datasets: GSE121248, representing HBV-associated HCC, and GSE41804, representing HCV-associated HCC with IL28B rs8099917 stratification [11,12]. GSE83148 was used to derive an HBV injury axis based on chronic hepatitis B clinical strata, including ALT, AST, and HBV-DNA [13]. GSE38941, an HBV-associated acute liver failure cohort with repeated liver fragments per patient, was retained for contextual hepatitis-stage comparison and acute liver failure program construction [8].

To address external-validation concerns, additional HCC tumor/non-tumor cohorts were curated from the Gene Expression Omnibus. Validation datasets included GSE14520, GSE25097, GSE36376, GSE76427, GSE57957, GSE45267, and GSE112790 [14–20] when tumor/non-tumor labels could be recovered and sufficient sample numbers were available. Dataset inclusion required: human liver tissue transcriptomic data; genome-wide expression profiling; recoverable tumor and adjacent/non-tumor labels or, for TCGA-LIHC, clinical metadata; and adequate sample numbers for group-level inference. Exclusion criteria were: cell-line-only studies, treatment-response-only studies without baseline tumor tissue, datasets lacking recoverable phenotype labels, and datasets with insufficient sample numbers for tumor/non-tumor comparison.

All GEO datasets were downloaded from the NCBI GEO repository using R/Bioconductor workflows, including GEOquery where appropriate [9,21]. TCGA-LIHC RNA-seq and clinical metadata were obtained using TCGAbiolinks [22].

### Expression data acquisition, preprocessing, and gene mapping

For GEO microarray datasets, processed series matrix files were downloaded and imported into R/Bioconductor. Analyses were performed on log-scale expression values provided by the GEO submitters, typically RMA-normalized for Affymetrix platforms. No additional cross-study normalization was applied to the discovery cohorts before within-cohort modeling, to avoid mixing preprocessing assumptions across studies.

Probe identifiers were mapped to HGNC gene symbols using platform annotation tables. When multiple probes were mapped to the same gene symbol, one representative probe was retained. The representative probe was selected by highest variance or average expression across samples, depending on dataset-specific annotation quality, to reduce probe redundancy while preserving informative signal. Gene-symbol overlaps with MSigDB Hallmark gene sets and downstream module genes were audited for each curated dataset before scoring.

For TCGA-LIHC, RNA-seq expression values were transformed to log-scale expression prior to module scoring. TCGA sample barcodes were parsed to distinguish primary tumor and solid normal tissue, and clinical metadata were merged by patient barcode.

### Phenotype parsing, metadata curation, and quality control

Sample-level metadata were parsed from GEO series annotations and standardized into a common schema containing dataset accession, sample identifier, tissue class, etiology when available, and study-specific covariates. Tissue labels were harmonized as tumor or non-tumor/adjacent liver. Non-tumor labels were detected before tumor labels to avoid misclassification of strings such as “non-tumor” as tumor. For GSE41804, IL28B rs8099917 genotype was extracted and encoded as TT versus TG/GG when available.

Quality control included inspection of sample-label distributions, gene-symbol overlap, and multidimensional scaling (MDS) or principal-component summaries of expression matrices using limma-based exploratory plots [23]. Datasets lacking both tumor and non-tumor classes after metadata curation were excluded from tumor/non-tumor validation analyses.

### Differential expression modeling

Within each discovery HCC cohort, tumor-versus-adjacent/non-tumor differential expression was modeled using limma [23]. For GSE121248 and GSE41804, the primary design matrix included tissue status. For GSE41804, IL28B genotype and tissue-by-genotype interaction terms were additionally evaluated to test whether tumor-associated expression shifts differed by genotype. For GSE38941, where multiple liver fragments were sampled per acute liver failure patient, repeated measurements were modeled using duplicateCorrelation and patient-level blocking before empirical Bayes moderation. Multiple-testing correction used the Benjamini-Hochberg false-discovery rate (FDR) procedure [24].

### Hallmark pathway activity analysis

Hallmark pathway activities were quantified per sample using Gene Set Variation Analysis (GSVA) or ssGSEA-compatible GSVA scoring over the MSigDB Hallmark collection [6,7]. For microarray datasets, Gaussian-kernel GSVA was used when supported by the installed GSVA version. To ensure compatibility with recent Bioconductor releases, pathway scoring was implemented using the current parameter-object GSVA API when required, with fallback to older syntax for legacy versions.

Pathway-level tumor-versus-non-tumor effects were modeled using limma on sample-level Hallmark scores. Discovery analyses focused on conserved activation of proliferation and repair programs, including E2F targets, G2M checkpoint, MYC targets, mitotic spindle, and DNA repair, and suppression of hepatocyte functional programs, including xenobiotic metabolism, coagulation, bile acid metabolism, and fatty acid metabolism.

### HBV injury-axis derivation from GSE83148

An HBV injury axis was derived from GSE83148, a chronic HBV hepatitis cohort with ALT, AST, and HBV-DNA strata available in the GEO metadata [13]. Because these variables were encoded categorically, each component was recoded as an ordinal variable. “NON” values were encoded as 0, the lower abnormal or positive stratum as 1, and the higher abnormal or viral-load stratum as 2. Specifically, ALT, AST, and HBV-DNA were transformed into ordinal components and then independently z-standardized. A continuous HBV_INJURY_INDEX was defined as the mean of the standardized ALT, AST, and HBV-DNA components for each sample.

For each gene, expression was modeled as a function of the continuous HBV_INJURY_INDEX using limma [23]. Genes with positive coefficients were interpreted as positively injury-associated. To balance interpretability and robustness, ranked top-N injury programs were generated by ordering positive FDR-significant genes by FDR and moderated statistic. Based on top-N sensitivity analysis, the compact primary HBV injury representation was defined as the top 2,000 positive injury-associated genes, referred to as HBV_INJURY_TOP_2000. Broader sensitivity analyses used HBV_INJURY_TOP_5000 and the extended FDR-defined HBV_INJURY_EXTENDED_7792 program, comprising all positive genes with FDR < 0.10. Very compact top-200, top-500, and top-1000 definitions were retained as exploratory sensitivity checks but were not used as the primary injury representation because they showed lower stability.

HBV injury scores were computed in GSE121248 as the mean of gene-wise z-scored expression values across the selected injury-program genes present in the expression matrix.

### Gene-level meta-analysis and conserved module construction

To identify conserved HCC tumor-state genes across viral etiologies, gene-level meta-analysis was performed by combining the GSE121248 and GSE41804 tumor-versus-non-tumor contrasts. For each gene, cohort-specific log fold changes and standard errors were extracted from limma models and combined using inverse-variance fixed-effect meta-analysis. Between-cohort heterogeneity was quantified using tau-squared (τ^2^) and I-squared (I^2^), and random-effects estimates were retained as sensitivity analyses when heterogeneity was non-negligible [25].

Genes were considered conserved when they showed concordant direction across discovery cohorts and passed FDR control in the meta-analysis. Conserved upregulated and downregulated genes were ranked by combined evidence across cohorts. The top 20 conserved tumor-upregulated genes were used to define the ProlifHub module, enriched for proliferation and cell-cycle activity. The top 20 conserved tumor-downregulated genes were used to define the HepLoss module, enriched for hepatocyte differentiation and liver functional programs. To address the arbitrariness of the top-20 cutoff, module-size sensitivity analyses were performed using top-10, top-15, top-20, top-30, and top-50 definitions, and validation performance was compared across cohorts.

### Module scoring and HCCStateScore definition

For each dataset, module scores were computed from gene-wise z-scored expression values. For gene g in sample i, expression was standardized within dataset as:

Zgi = (Egi - mean(Eg)) / SD(Eg)

where Egi is the expression of gene g in sample i, and mean(Eg) and SD(Eg) are the mean and standard deviation of gene g across samples in that dataset.

The ProlifHubScore was defined as the mean of standardized expression values across ProlifHub genes. The HepLossScore was defined analogously as the mean of standardized expression values across HepLoss genes. The composite HCCStateScore was defined as:

HCCStateScore = ProlifHubScore - HepLossScore

Thus, high HCCStateScore values indicate coordinated activation of proliferation-associated genes and suppression of hepatocyte functional genes.

### External validation across GEO cohorts

External validation was performed by recomputing ProlifHubScore, HepLossScore, and HCCStateScore in independent HCC cohorts. GSE14520 was retained as the original validation cohort, and additional validation cohorts included GSE25097, GSE36376, GSE76427, GSE57957, GSE45267, and GSE112790 when tumor/non-tumor labels were recoverable. Dataset-level sample counts and inclusion decisions are provided in Supplementary Table S1.

For each validation cohort, tumor-versus-non-tumor differences were quantified using mean difference, Welch’s t-test, Benjamini-Hochberg FDR correction when multiple cohorts or scores were considered, and standardized effect size using Cohen’s d [26]. Tumor-versus-non-tumor discrimination by HCCStateScore was quantified using the area under the receiver-operating-characteristic curve (AUC) with pROC [27]. Validation was interpreted as transcriptomic tumor-state separability in tissue expression profiles, not as evidence of early diagnostic performance.

To evaluate whether module performance depended on the selected module size, module-size sensitivity analysis was performed across top-10, top-15, top-20, top-30, and top-50 gene definitions. HCCStateScore AUC and tumor-versus-non-tumor deltas were compared across datasets and module sizes.

### CIBERSORTx-adjusted modeling of the HBV injury axis

To assess whether the HBV injury signal in GSE121248 could be explained by proliferative state or immune-cell admixture, CIBERSORTx-based immune-composition adjustment was performed [28,29]. A GSE121248 expression mixture file was exported with genes as rows and samples as columns. CIBERSORTx was run externally using the LM22 signature matrix in relative mode. Because the input data were microarray-derived, quantile normalization was enabled. The resulting inferred immune-cell fractions were imported into R and matched to GSE121248 sample identifiers.

Because CIBERSORTx cell fractions are compositional and collinear, two adjustment strategies were used. First, centered log-ratio transformed fractions were summarized by principal components, and the leading CIBERSORTx principal components were included as immune-composition covariates. Second, a selected-fraction sensitivity model included the most variable inferred immune fractions while avoiding singular model fits.

Nested linear models were fit in GSE121248 with the HBV injury score as the outcome. For each HBV injury gene-set definition, the following models were evaluated:

HBV_INJURY ~ tissue

HBV_INJURY ~ tissue + E2F + G2M

HBV_INJURY ~ tissue + E2F + G2M + CIBERSORTx PCs

HBV_INJURY ~ tissue + E2F + G2M + selected CIBERSORTx fractions

where tissue was encoded as non-tumor versus tumor, and E2F and G2M corresponded to Hallmark E2F targets and G2M checkpoint GSVA scores. The tumor coefficient was compared across nested models to estimate attenuation after proliferation and immune-composition adjustment. Because all analyses were performed on bulk transcriptomes, persistence of the HBV injury signal after adjustment was interpreted as evidence of a residual tumor-associated injury component, not as definitive proof of tumor-cell-intrinsic expression.

### TCGA-LIHC validation and survival modeling

TCGA-LIHC RNA-seq and clinical data were obtained using TCGAbiolinks [22]. ProlifHubScore, HepLossScore, and HCCStateScore were recomputed in TCGA-LIHC using the same gene-wise z-score approach described above. Tumor and solid-normal tissue labels were inferred from TCGA barcode sample codes and cross-checked against available sample annotations.

Overall-survival analyses were performed using Cox proportional-hazards models with continuous module scores. Survival time was calculated from days to death when available or days to last follow-up otherwise. Vital status was encoded as event status. Cox models were fitted for each module score as a standardized continuous predictor. Three model classes were evaluated: score-only models, age/sex-adjusted models, and age/sex/pathologic-stage-adjusted models in the subset with complete staging information. Results were reported as hazard ratios per one-standard-deviation increase in score, with 95% confidence intervals and p-values. These analyses were interpreted as prognostic coherence checks rather than definitive clinical biomarker validation.

### Statistical analysis and reproducibility

All analyses were performed in R using Bioconductor and CRAN packages including limma, GSVA, msigdbr, metafor, pROC, TCGAbiolinks, survival, broom, tidyverse, and related dependencies [6,7,21–23,25,27,30]. Continuous variables were summarized using means, medians, standard deviations, and confidence intervals where appropriate. Group differences were evaluated using limma models or Welch’s t-tests depending on the analysis level. Multiple testing was controlled using the Benjamini-Hochberg FDR procedure [24]. ROC analyses used AUC estimates from pROC [27]. Meta-analytic heterogeneity was quantified using I^2^ and REML-estimated τ^2^ [25]. All intermediate files, curated metadata tables, module definitions, validation outputs, and figure-ready tables were generated through reproducible R scripts.

## Results

### Cross-etiology Hallmark program shifts define a conserved HCC tumor state

In HBV-associated HCC (GSE121248), from the evaluation of tumor-to-adjacent/non-tumor differences at the pathway level, tumors showed strong activation of proliferative and replication-stress programs, including E2F targets, G2M checkpoint, MYC targets, mitotic spindle, DNA repair, unfolded protein response, WNT/β-catenin signaling, protein secretion, PI3K/AKT/mTOR signaling, and MTORC1 signaling. In parallel, tumors showed suppression of hepatocyte functional and liver metabolic programs, with several hepatocyte metabolic and liver functional programs showing negative tumor-associated shifts.

The HCV-associated HCC cohort GSE41804 reproduced the same dominant tumor-state structure, with positive tumor-associated shifts in E2F, G2M, MYC, mitotic spindle, and DNA repair programs, and negative shifts in xenobiotic metabolism, bile acid metabolism, coagulation, and fatty acid metabolism. These pathway-level results support a conserved cross-etiology HCC state characterized by proliferation/repair activation and loss of hepatocyte functional programs rather than a cohort-specific artifact. These cross-etiology Hallmark pathway shifts are summarized in Fig. 2.

**Figure 2.**
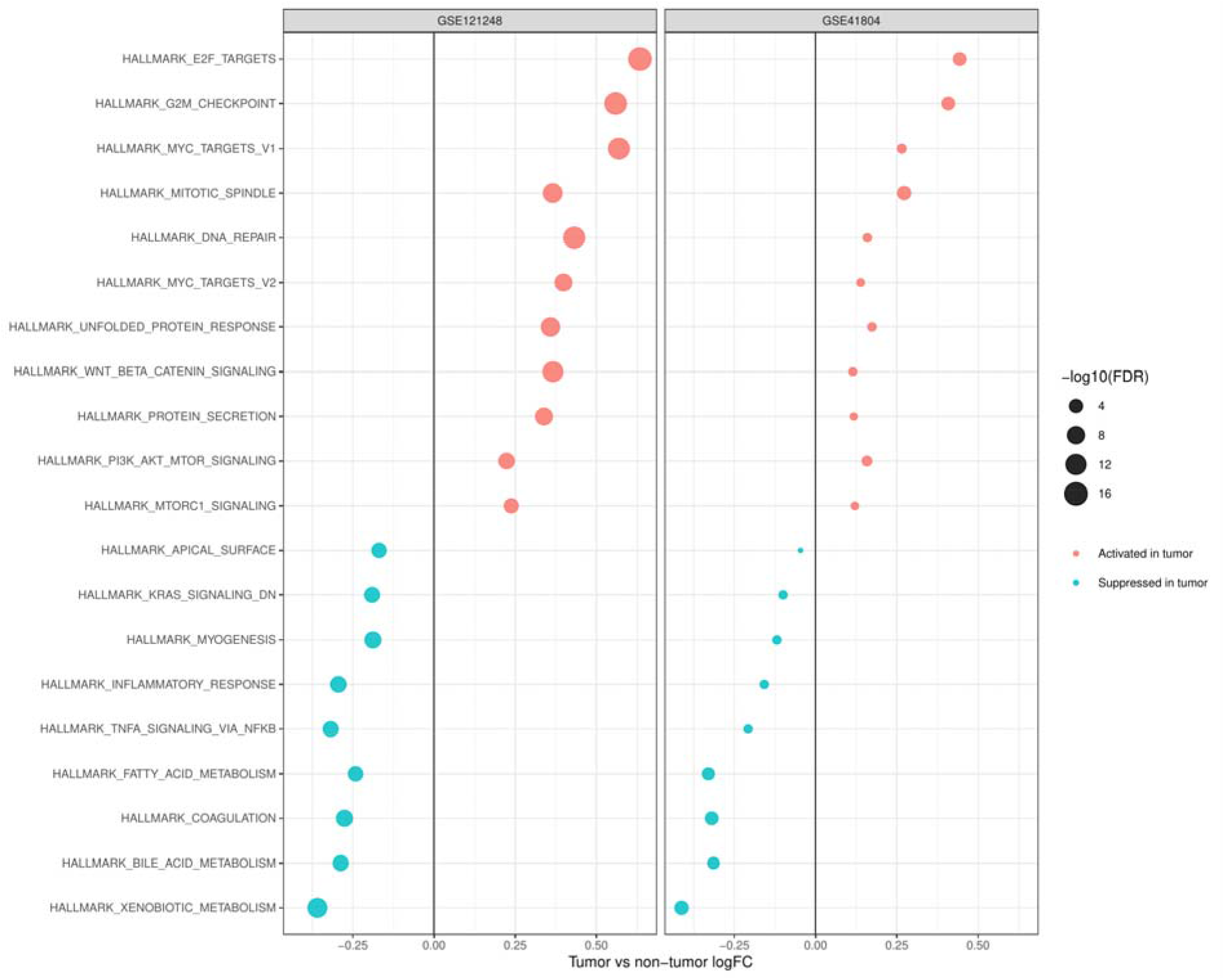
Cross-etiology Hallmark program shifts in HBV- and HCV-associated HCC. Dot plot summarizing Hallmark pathway differential activity, reported as tumor versus adjacent/non-tumor logFC with corresponding −log10(FDR), in GSE121248 and GSE41804. Red indicates activation in tumor and cyan indicates suppression in tumor. Across both cohorts, tumors show consistent enrichment of proliferation/repair programs and depletion of hepatocyte functional programs.

### Gene-level meta-analysis identifies conserved tumor-state signals and quantifies heterogeneity

We next performed gene-level meta-analysis across the two discovery cohorts to identify conserved tumor-associated expression changes. A total of 21,753 genes were estimable in the meta-analysis, with no non-estimable genes after preprocessing and gene-symbol harmonization. Of these, 12,444 genes reached fixed-effect meta-analysis FDR < 0.05. Heterogeneity analysis showed a median I^2^ of 0 and mean I^2^ of 25.71, although 6,249 genes showed I^2^ > 50%, indicating that a subset of genes displayed appreciable between-cohort heterogeneity. Therefore, conserved module construction emphasized concordant directionality and ranked cross-cohort evidence rather than significance alone. These meta-analysis and heterogeneity summaries are reported in Table 1.

**Table 1.** Gene-level meta-analysis summary and heterogeneity statistics across GSE121248 and GSE41804. I^2^ and τ^2^ were used to quantify between-cohort heterogeneity. I^2^ summarizes the proportion of variability attributable to between-cohort heterogeneity, whereas τ^2^ denotes the REML-estimated between-cohort variance from the random-effects sensitivity model.

| Metric | Value |
| --- | --- |
| Meta-analysis rows | 21,753 |
| Estimable genes | 21,753 |
| Non-estimable genes | 0 |
| Genes with fixed-effect meta-analysis FDR < 0.05 | 12,444 |
| Median $I^2$ | 0 |
| Mean $I^2$ | 25.71 |
| Genes with $I^2 > 50$ | 6,249 |
| Median $\tau^2$ (REML) | 0 |
| Mean $\tau^2$ (REML) | 0.07 |
| Genes with $\tau^2$ (REML) > 50 | 9,780 |

### Conserved genes define compact proliferation and hepatocyte-loss modules

Conserved tumor-upregulated and tumor-downregulated genes were distilled into compact module scores. The top conserved tumor-upregulated genes defined the ProlifHub module, capturing proliferation and cell-cycle-associated activity. The top conserved tumor-downregulated genes defined the HepLoss module, capturing loss of hepatocyte differentiation and liver functional programs. These modules were scored as mean gene-wise z-scored expression values, and the composite HCCStateScore was defined as ProlifHubScore minus HepLossScore. High HCCStateScore values therefore represent simultaneous activation of proliferative hubs and suppression of hepatocyte functional genes.

In both discovery cohorts, tumors showed higher ProlifHubScore and lower HepLossScore relative to adjacent/non-tumor tissue, while the composite HCCStateScore produced the clearest tumor/non-tumor separation (Fig. 3), consistent with the Hallmark-level observation that the dominant conserved HCC state contrasts proliferation/repair activation with hepatocyte functional loss.

**Figure 3.**
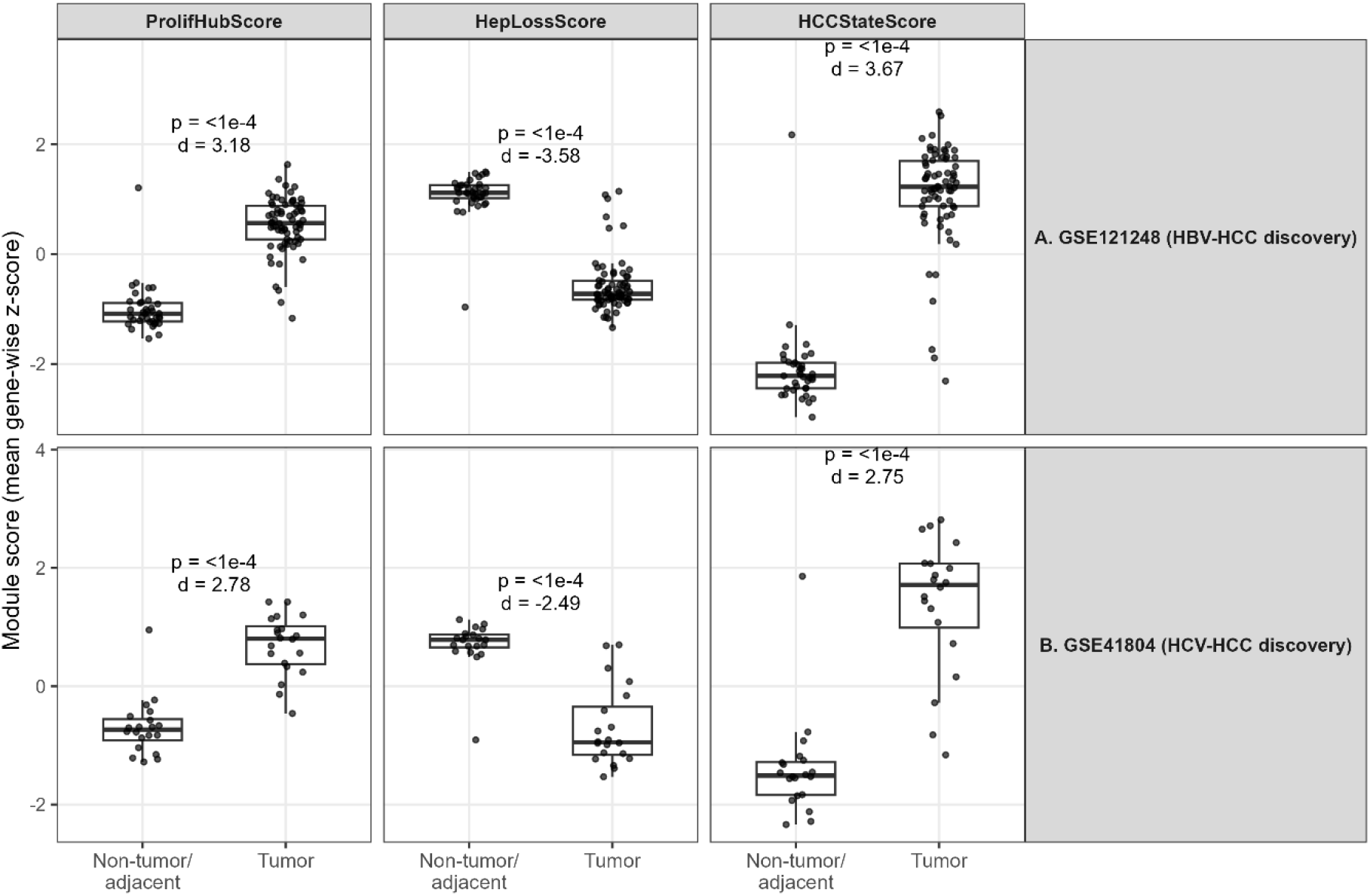
Discovery-cohort module score behavior in HBV- and HCV-associated HCC. Boxplots showing ProlifHubScore, HepLossScore, and HCCStateScore in tumor versus adjacent/non-tumor tissue from the two discovery cohorts, GSE121248 for HBV-associated HCC and GSE41804 for HCV-associated HCC. ProlifHubScore summarizes conserved tumor-upregulated proliferation/cell-cycle genes, HepLossScore summarizes conserved tumor-downregulated hepatocyte functional genes, and HCCStateScore is defined as ProlifHubScore minus HepLossScore. Across both discovery cohorts, tumors show increased ProlifHubScore, decreased HepLossScore, and increased HCCStateScore, supporting a conserved tumor-state axis characterized by proliferation/repair activation opposed to hepatocyte functional loss. Boxes indicate the interquartile range with median center lines; whiskers represent 1.5× the interquartile range. Reported p-values are from Welch’s t-tests, and effect sizes are shown as Cohen’s d.

### HCCStateScore validates across multiple independent HCC cohorts

We then tested whether the module scores generalized beyond the discovery datasets. ProlifHubScore, HepLossScore, and HCCStateScore were recomputed in independent GEO HCC cohorts with recoverable tumor/non-tumor labels. Across GSE14520, GSE25097, GSE36376, GSE76427, GSE57957, GSE45267, and GSE112790, HCCStateScore consistently increased in tumor tissue relative to non-tumor or adjacent liver. Tumor-versus-non-tumor discrimination was high across validation cohorts, with AUC values clustering around approximately 0.95–0.99. The multi-cohort validation pattern is summarized in Fig. 4.

**Figure 4.**
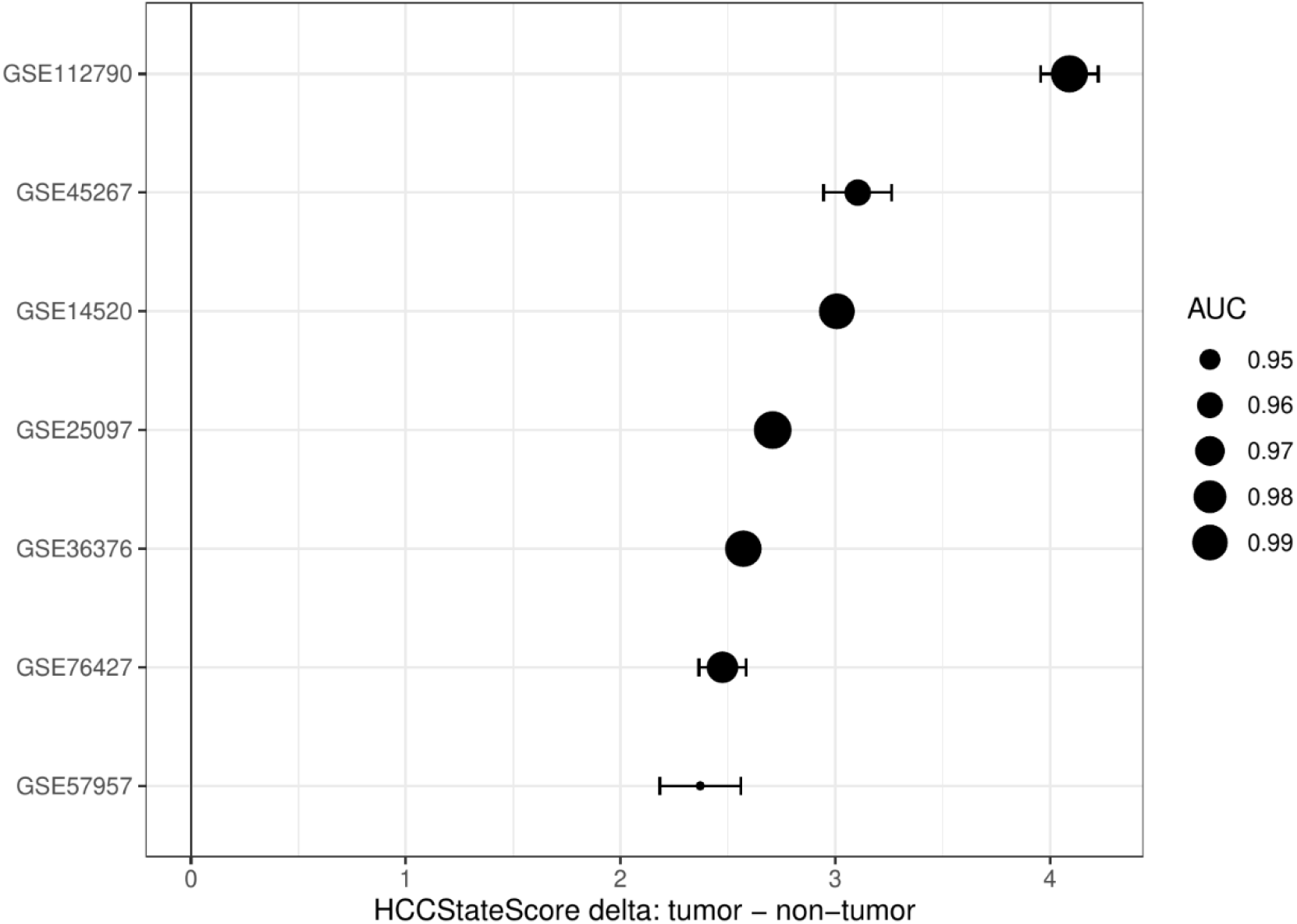
Multi-cohort validation of HCCStateScore. HCCStateScore tumor minus non-tumor deltas are shown across independent GEO validation cohorts. Point size/color denotes AUC, showing consistently high tumor/non-tumor discrimination across datasets.

These results support the portability of the module framework across independent HCC transcriptomic datasets. The validation should be interpreted as robust tissue-level tumor-state separability in bulk transcriptomes, not as evidence for early detection or non-invasive diagnostic performance.

### Module-size sensitivity analysis supports robustness of the HCCStateScore

To evaluate whether the module framework depended on the arbitrary top-20 cutoff, we repeated module construction using top-10, top-15, top-20, top-30, and top-50 gene definitions. HCCStateScore AUC remained high across module sizes in the discovery and validation cohorts, indicating that the score captured a stable proliferation-versus-hepatocyte-loss axis rather than a narrowly optimized gene list. Module-size robustness across discovery and validation cohorts is summarized in Supplementary Fig. S3.

### A compact HBV injury program remains tumor-associated after proliferation and immune-composition adjustment

We next evaluated whether hepatitis-related injury biology was detectable within HBV-associated tumors and whether this signal could be explained by proliferative state or immune-cell admixture. An HBV injury axis was derived from GSE83148 using an ordinal ALT/AST/HBV-DNA injury index. Ranked top-N HBV injury programs and an extended FDR-defined injury program were then projected into the HBV-HCC discovery cohort GSE121248.

Very compact top-200 and top-500 injury definitions were less stable, suggesting that the injury axis reflects a broad liver-injury-associated transcriptional gradient rather than a small discrete module. The top-2000 ranked injury program was therefore selected as the compact primary representation because it balanced interpretability and stability. In GSE121248, HBV_INJURY_TOP_2000 was elevated in tumor tissue in the unadjusted model (tumor coefficient = 0.237, 95% CI 0.0886–0.386, p = 0.00203). The coefficient was attenuated after E2F/G2M adjustment alone and was not statistically significant (coefficient = 0.130, 95% CI −0.0494–0.310, p = 0.153), indicating partial overlap between injury and proliferative/regenerative state. However, the tumor coefficient remained positive and significant after simultaneous adjustment for E2F/G2M activity and CIBERSORTx-derived immune-composition principal components (coefficient = 0.190, 95% CI 0.0281–0.352, p = 0.0220), and in the selected-fraction sensitivity model (coefficient = 0.188, 95% CI 0.0383–0.338, p = 0.0144) [28,29].

The extended 7,792-gene FDR-defined HBV_INJURY program confirmed the same qualitative conclusion and showed stronger statistical robustness across all adjustment models. The attenuation and persistence of the tumor-associated HBV injury coefficient across nested proliferation- and CIBERSORTx-adjusted models are summarized in Fig. 5. In the extended set, the tumor coefficient remained significant after E2F/G2M adjustment and after simultaneous E2F/G2M plus CIBERSORTx adjustment. Together, these results indicate that the HBV injury-associated signal is partly coupled to proliferation but is not fully explained by proliferative state or inferred immune-cell composition. Because the analysis is based on bulk transcriptomes, this should be interpreted as a residual tumor-associated HBV injury component, not as definitive proof of tumor-cell-intrinsic expression. The corresponding regression estimates for the compact and extended HBV injury programs are reported in Table 2.

**Table 2.** HBV injury-axis regression in GSE121248 before and after adjustment for proliferative state and inferred immune composition. The tumor coefficient represents the difference between tumor and adjacent/non-tumor tissue. Effect retained is calculated relative to the unadjusted model within each injury-set definition.

| HBV injury set | Role | Model | Tumor coefficient | Effect retained | p-value | 95% CI |
| --- | --- | --- | --- | --- | --- | --- |
| TOP 2000 | Primary compact | Unadjusted | 0.237 | 100% | 0.00203 | 0.0886–0.386 |
| TOP 2000 | Primary compact | E2F/G2M adjusted | 0.130 | 55.0% | 0.153 | –0.0494–0.310 |
| TOP 2000 | Primary compact | E2F/G2M + CIBERSORTx PC | 0.190 | 80.2% | 0.0220 | 0.0281–0.352 |
| TOP 2000 | Primary compact | E2F/G2M + selected fractions | 0.188 | 79.4% | 0.0144 | 0.0383–0.338 |
| EXTENDED 7792 | Sensitivity | Unadjusted | 0.233 | 100% | 5.92e–9 | 0.160–0.306 |
| EXTENDED 7792 | Sensitivity | E2F/G2M adjusted | 0.125 | 53.7% | 0.00408 | 0.0406–0.209 |
| EXTENDED 7792 | Sensitivity | E2F/G2M + CIBERSORTx PC | 0.148 | 63.4% | 0.000500 | 0.0662–0.229 |
| EXTENDED 7792 | Sensitivity | E2F/G2M + selected fractions | 0.133 | 57.3% | 0.000518 | 0.0597–0.207 |

**Figure 5.**
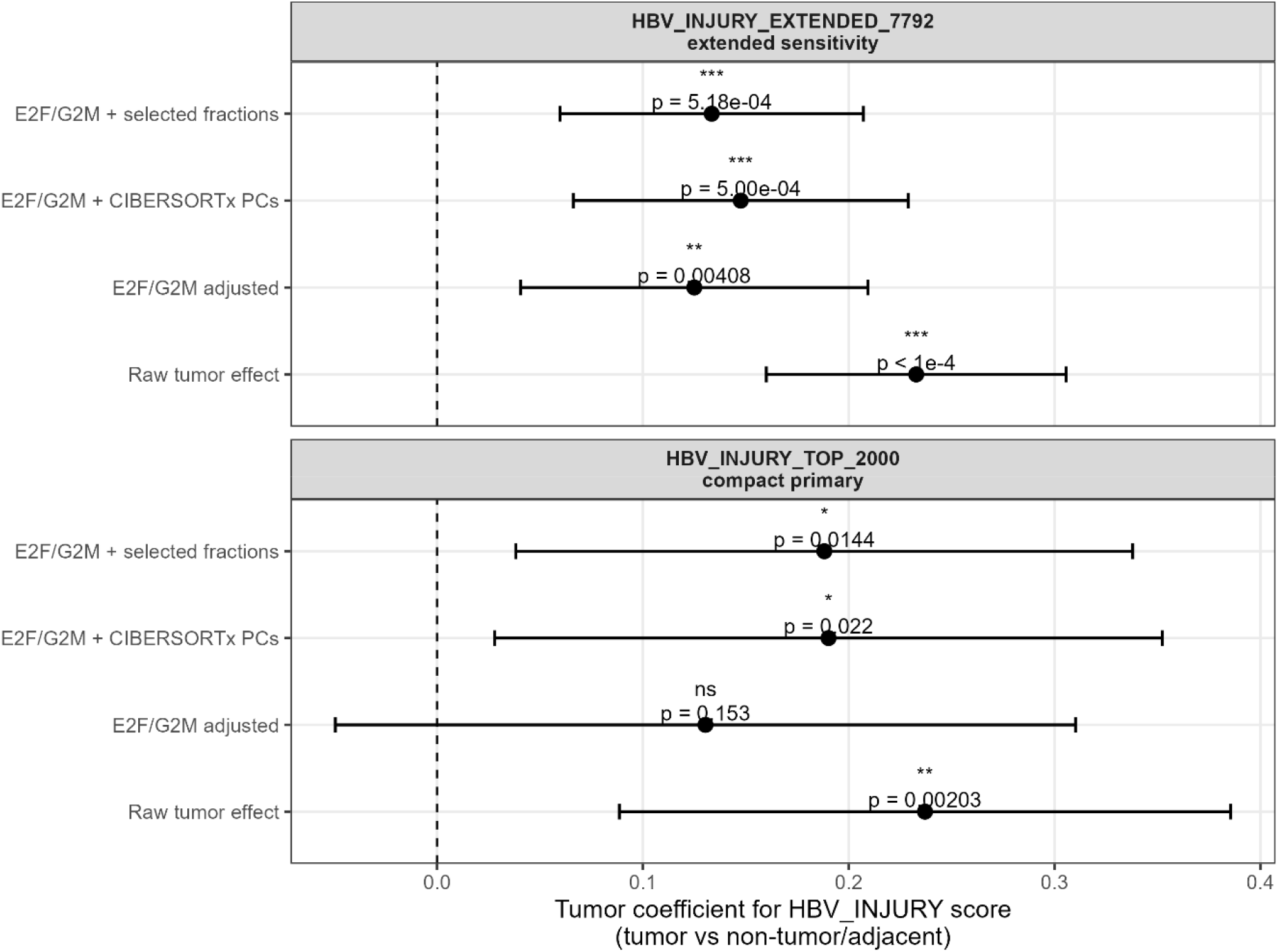
HBV injury-axis tumor coefficient before and after adjustment for proliferative state and inferred immune composition. Coefficient plot showing the tumor-associated effect on HBV_INJURY scores in GSE121248 across nested linear models. The compact ranked HBV_INJURY_TOP_2000 program was used as the primary injury representation, and the extended HBV_INJURY_EXTENDED_7792 program was retained as a sensitivity analysis. Points indicate the tumor coefficient relative to adjacent/non-tumor liver, and horizontal bars indicate 95% confidence intervals. Models include the unadjusted tumor effect, adjustment for E2F targets and G2M checkpoint Hallmark activity, simultaneous adjustment for E2F/G2M plus CIBERSORTx-derived immune-composition principal components, and adjustment for E2F/G2M plus selected CIBERSORTx-inferred immune-cell fractions. The compact top-2000 program remained positive and significant after combined proliferation and immune-composition adjustment, while the extended FDR-defined program showed the same direction with stronger statistical robustness. The results support a residual tumor-associated HBV injury component that is not fully explained by proliferative activity or inferred immune-cell composition.

### TCGA-LIHC Cox modeling supports prognostic coherence of the module framework

We performed module-level Cox proportional-hazards modeling in TCGA-LIHC. Module scores were modeled as standardized continuous predictors. Higher ProlifHubScore was associated with worse overall survival in the score-only model (HR = 1.44 per SD, 95% CI 1.20–1.73, p = 8.46 × 10−5), after age/sex adjustment (HR = 1.49, 95% CI 1.23–1.79, p = 3.51 × 10−5), and after additional pathologic-stage adjustment in the subset with complete staging information (HR = 1.44, 95% CI 1.13–1.83, p = 0.00324).

HCCStateScore showed similar prognostic coherence, including in the age/sex/stage-adjusted model (HR = 1.51 per SD, 95% CI 1.12–2.03, p = 0.00641). HepLossScore showed the expected protective direction but did not reach statistical significance. These results indicate that the composite state score captures survival-relevant tumor biology, primarily reflecting the adverse prognostic contribution of proliferative activation. Because the stage-adjusted analysis used a smaller complete-case subset, these results are interpreted as prognostic coherence rather than definitive clinical validation. The Cox model estimates are summarized in Table 3.

**Table 3.** TCGA-LIHC overall-survival Cox proportional-hazards models for module scores. Hazard ratios are reported per one-standard-deviation increase in score. Stage-adjusted models used the subset with complete pathologic-stage information.

| Model | Score | HR per SD | 95% CI | p-value | n | events |
| --- | --- | --- | --- | --- | --- | --- |
| Score only | ProlifHubScore | 1.44 | 1.20–1.73 | 8.46e–5 | 369 | 130 |
| Score only | HepLossScore | 0.863 | 0.713–1.04 | 0.131 | 369 | 130 |
| Score only | HCCStateScore | 1.43 | 1.17–1.75 | 5.07e–4 | 369 | 130 |
| Age/sex adjusted | ProlifHubScore | 1.49 | 1.23–1.79 | 3.51e–5 | 369 | 130 |
| Age/sex adjusted | HepLossScore | 0.866 | 0.714–1.05 | 0.140 | 369 | 130 |
| Age/sex adjusted | HCCStateScore | 1.45 | 1.18–1.78 | 3.92e–4 | 369 | 130 |
| Age/sex/stage adjusted | ProlifHubScore | 1.44 | 1.13–1.83 | 0.00324 | 173 | 74 |
| Age/sex/stage adjusted | HepLossScore | 0.882 | 0.661–1.18 | 0.394 | 173 | 74 |
| Age/sex/stage adjusted | HCCStateScore | 1.51 | 1.12–2.03 | 0.00641 | 173 | 74 |

## Discussion

This study was designed to distinguish transcriptomic programs that are conserved across viral etiologies of hepatocellular carcinoma from residual components that may retain etiology-linked structure. The revised analyses support three main conclusions. First, HBV- and HCV-associated HCC converge on a highly reproducible bulk tumor-state axis characterized by activation of proliferation and repair programs and suppression of hepatocyte functional programs. Second, this axis can be summarized by compact, interpretable module scores that generalize across multiple independent HCC cohorts. Third, an HBV-derived injury axis remains detectable in HBV-HCC tumors after adjustment for proliferative state and inferred immune-cell composition, although the bulk-transcriptomic nature of the analysis prevents definitive assignment of this signal to malignant hepatocytes.

The conserved proliferation-versus-hepatocyte-function axis observed here is not, by itself, a new biological principle in HCC. Prior molecular taxonomies and large-scale genomic studies have repeatedly shown that aggressive HCC states are associated with cell-cycle activation, MYC/E2F-related programs, dedifferentiation, and reduced hepatocyte metabolic identity [5,31]. The contribution of the present work is therefore not the discovery that HCC tumors are proliferative and hepatocyte functions are suppressed. Rather, the contribution is methodological and integrative: we convert this established biology into a compact, reproducible, cross-cohort scoring framework and use it as a quantitative tumor-state axis against which etiology-linked residual signals can be tested.

The Hallmark-level results provide the broad biological foundation for this framework. Across the HBV-HCC discovery cohort GSE121248 and the HCV-HCC discovery cohort GSE41804, tumors consistently showed activation of E2F targets, G2M checkpoint, MYC targets, mitotic spindle, DNA repair, and related proliferative or repair-associated programs, together with suppression of xenobiotic metabolism, bile acid metabolism, coagulation, fatty-acid metabolism, and related hepatocyte functional pathways. This pattern is consistent with TCGA-LIHC and other integrative HCC studies in which cell-cycle activation and dedifferentiation are associated with tumor progression and adverse clinicopathologic features [5,31]. By analyzing HBV- and HCV-associated tumors using the same pathway-scoring framework, the present study shows that this dominant tumor-state structure is reproducible across viral etiologies rather than being specific to one hepatitis background.

The module-based strategy was intended to improve interpretability and transferability. Many gene-level signatures are difficult to compare across cohorts because they may encode platform-specific effects, cohort composition, or unstable single-gene behavior. By selecting conserved tumor-upregulated and tumor-downregulated genes through cross-cohort meta-analysis and summarizing them as mean z-score modules, the ProlifHubScore and HepLossScore intentionally trade gene-level granularity for stability. The composite HCCStateScore further captures the opposition between proliferative activation and hepatocyte functional loss. Importantly, the expanded validation demonstrates that this score is not restricted to the original validation dataset. HCCStateScore showed consistently positive tumor-versus-non-tumor deltas and high AUC values across multiple independent GEO cohorts, and module-size sensitivity analysis showed that performance remained stable across top-10 to top-50 gene definitions. These findings directly reduce the likelihood that the result is an artifact of the top-20 cutoff or of one validation cohort.

The clinical interpretation of HCCStateScore should nevertheless remain cautious. The high AUC values observed in tumor-versus-non-tumor tissue comparisons demonstrate strong transcriptomic separation between resected tumor and non-tumor liver samples. They should not be interpreted as evidence for early detection performance, non-invasive diagnostic utility, or clinical screening applicability. The most appropriate use of HCCStateScore at this stage is as a quantitative bulk tumor-state covariate that captures proliferation and hepatocyte-loss biology. Such a covariate can be useful when testing etiology-linked effects, immune programs, microenvironmental signatures, or outcome associations while accounting for dominant tumor-state variation.

The TCGA-LIHC analysis supports the prognostic coherence of this tumor-state framework. Higher ProlifHubScore was associated with poorer overall survival in continuous Cox models and remained significant after adjustment for age, sex, and pathologic stage. HCCStateScore showed a similar association, including in the age/sex/stage-adjusted model. By contrast, HepLossScore showed the expected protective direction but did not reach statistical significance. These findings suggest that the survival association of the composite score is driven primarily by the proliferative component, which is consistent with the known adverse prognostic impact of cell-cycle activation in HCC [5,21]. These results should be viewed as supportive prognostic coherence rather than definitive clinical biomarker validation, particularly because the stage-adjusted model relied on the subset of TCGA-LIHC cases with complete staging information.

A second major focus of the manuscript is the HBV injury axis. In a first analysis, the HBV injury score remained elevated after E2F/G2M adjustment, but this did not address whether the signal reflected immune or stromal admixture. We therefore revised the injury-axis analysis in two ways. First, the HBV injury program was explicitly derived from an ordinal ALT/AST/HBV-DNA injury index in GSE83148. Second, the projected injury score in GSE121248 was evaluated in nested regression models that adjusted for proliferation and CIBERSORTx-inferred immune composition. This directly addresses the concern that the HBV injury signal could simply reflect cell-cycle activity or immune-cell content.

The top-N sensitivity analysis indicates that the HBV injury axis is broad rather than a small discrete transcriptional module. Very compact top-200 and top-500 injury programs were less stable, whereas the top-2000 program provided a more reliable compact representation. The HBV_INJURY_TOP_2000 score was elevated in HBV-HCC tumor tissue in the unadjusted model and remained significant after simultaneous adjustment for E2F/G2M activity and CIBERSORTx-derived immune-composition principal components. The extended 7,792-gene FDR-defined injury program showed the same direction and stronger statistical robustness across all models. These results suggest that the injury signal is partly coupled to proliferation but not fully explained by proliferative state or inferred immune-cell composition.

Mechanistically, several non-mutually exclusive explanations could account for this residual tumor-associated HBV injury signal. It may reflect persistent inflammatory or stress-response programs associated with chronic HBV infection, remodeling of the tumor microenvironment, injury-regeneration biology in adjacent diseased liver, or transcriptional consequences of HBV-associated carcinogenic processes such as long-term antigenic stimulation and viral integration [3,4]. The CIBERSORTx-adjusted analysis argues against the simplest explanation that the signal is entirely due to inferred immune-cell composition. However, bulk transcriptomes cannot determine whether the residual signal arises from malignant hepatocytes, non-malignant hepatocytes, stromal cells, infiltrating immune populations, or mixtures of these compartments. Therefore, the appropriate conclusion is that HBV-HCC retains a residual tumor-associated injury component, not that a tumor-cell-intrinsic HBV injury program has been proven.

Several limitations remain. First, the discovery analyses relied on processed public microarray matrices. Although this is standard for secondary transcriptomic studies and supports reproducibility, it limits direct control over preprocessing and batch-correction choices. Second, while the discovery cohorts were platform-compatible and the validation cohorts substantially expand generalizability, effect sizes may still vary across platforms, preprocessing pipelines, and clinical composition. Third, all primary analyses used bulk transcriptomes, which are sensitive to tumor purity, immune infiltration, stromal content, and adjacent-liver heterogeneity. CIBERSORTx adjustment mitigates this concern but does not replace single-cell or spatial localization. Fourth, the primary discovery design focused on viral hepatitis-associated HCC. Additional analyses in metabolic-, alcohol-, and autoimmune-associated HCC will be required to determine whether the same two-axis structure and residual injury components generalize across the broader etiologic spectrum of liver cancer [2]. Finally, the HBV injury axis was derived from one chronic HBV cohort using available ordinal clinical strata, and should be further validated using independent hepatitis cohorts with richer clinical, histologic, and molecular annotations.

Future work should proceed in three directions. First, the scoring framework should be applied to larger multi-etiology RNA-seq cohorts to determine whether HCCStateScore captures a universal tumor-state axis or whether additional etiology-specific modules are required. Second, single-cell and spatial transcriptomic datasets should be used to localize the HBV injury signal to malignant hepatocytes, non-malignant hepatocytes, immune cells, stromal cells, or spatially organized injury niches. Third, HCCStateScore and residualized injury scores should be tested as covariates or stratification variables in models of recurrence, therapy response, immune-checkpoint activity, and molecular subtype. Such analyses would clarify whether the present module framework is most useful for biological interpretation, patient stratification, or adjustment of downstream transcriptomic analyses.

## Conclusion

Across HBV- and HCV-associated HCC discovery cohorts and multiple independent validation datasets, this study identifies a reproducible bulk transcriptomic tumor-state axis defined by coordinated activation of proliferation/repair programs and suppression of hepatocyte functional programs. By compressing conserved gene-level signals into ProlifHubScore, HepLossScore, and the composite HCCStateScore, the analysis provides a compact framework for quantifying this established HCC biology across cohorts.

The validation demonstrates that HCCStateScore generalizes across multiple GEO cohorts and remains robust across module-size definitions, supporting its use as a portable tumor-state score rather than as a stand-alone diagnostic classifier. TCGA-LIHC Cox models further show that higher ProlifHubScore and HCCStateScore are associated with poorer overall survival, indicating prognostic coherence driven mainly by proliferative activation.

In HBV-associated HCC, a ranked HBV injury program derived from an ALT/AST/HBV-DNA injury index remains detectable in tumor tissue after adjustment for proliferative activity and inferred immune composition. The compact top-2000 HBV injury program and the extended FDR-defined injury set support the same qualitative conclusion: the HBV injury signal is partially coupled to proliferation but is not fully explained by cell-cycle activity or immune-cell admixture. Because these analyses are based on bulk tissue profiles, the signal should be interpreted as a residual tumor-associated HBV injury component rather than definitive evidence of tumor-cell-intrinsic injury biology.

Together, these findings support a revised two-level interpretation of viral HCC transcriptomes: a conserved tumor-state axis shared across HBV and HCV tumors, and a residual HBV-linked injury dimension that persists after accounting for major proliferative and immune-composition effects. This framework provides a reproducible basis for future multi-etiology, single-cell, spatial, and clinically annotated studies aimed at localizing injury-associated signals and testing their prognostic or therapeutic relevance.

## Declarations

The authors declare that they have no known competing financial interests or personal relationships that could have appeared to influence the work reported in this paper.

There was no funding for this study.

The study used publicly available, de-identified transcriptomic and clinical metadata. Formal institutional review board approval was not required because the analysis was restricted to secondary use of public, de-identified data.

## Data availability

All GEO transcriptomic datasets analyzed in this study are publicly available through the Gene Expression Omnibus under accession numbers GSE121248, GSE41804, GSE83148, GSE38941, GSE14520, GSE25097, GSE36376, GSE76427, GSE57957, GSE45267, and GSE112790. TCGA-LIHC RNA-seq and clinical data are available through the Genomic Data Commons and were accessed using TCGAbiolinks. Processed intermediate files generated in this study, including curated metadata tables, Hallmark scoring outputs, meta-analysis tables, module score tables, validation statistics, CIBERSORTx-adjusted regression outputs, TCGA-LIHC Cox-model outputs, and figure-ready summary files, are available at https://github.com/ricardo-romero-ochoa/hcc_cross_etiology_revised and archived at https://doi.org/10.5281/zenodo.20298127. TCGA-LIHC expression and clinical analyses were performed locally using the reproducible R pipeline described in the Methods.

## Authors’ contribution

Ricardo Romero

Conceptualization (Lead), Data curation (Equal), Formal analysis (Equal), Investigation (Equal), Methodology (Equal), Project administration (Lead), Supervision (Lead), Validation (Equal), Writing – original draft (Lead), Writing – review & editing (Lead).

Cinthia C. Toledo

Conceptualization (Supporting), Data curation (Equal), Formal analysis (Equal), Investigation (Equal), Methodology (Equal), Project administration (Supporting), Validation (Equal), Writing – original draft (Supporting), Writing – review & editing (Supporting).

