## Supplementary Figures for "Cross-etiology transcriptomic conservation in hepatocellular carcinoma reveals opposing proliferation and hepatocyte-loss programs validated across cohorts"

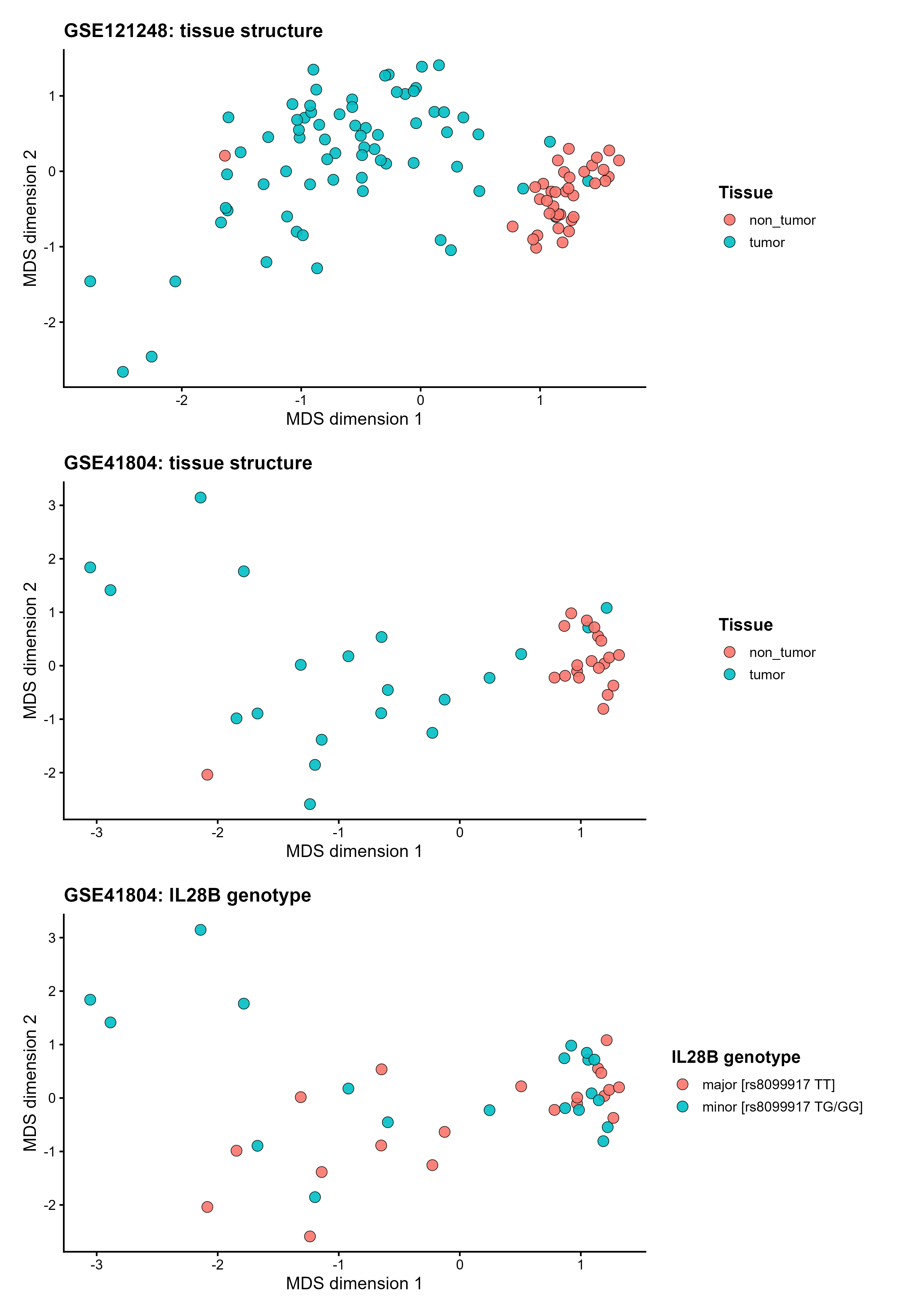
Supplementary Figures

**Supplementary Figure S1. Discovery-cohort quality control by multidimensional scaling.** MDS plots based on gene-level expression matrices for the discovery cohorts. Panel A shows GSE121248 samples colored by tissue type. Panel B shows GSE41804 samples colored by tissue type. Panel C shows GSE41804 samples colored by IL28B rs8099917 genotype. The dominant separation is tissue-associated, supporting the use of tumor versus adjacent/non-tumor contrasts as the primary discovery comparison. These plots were used for exploratory quality control and visual assessment of large-scale sample structure.

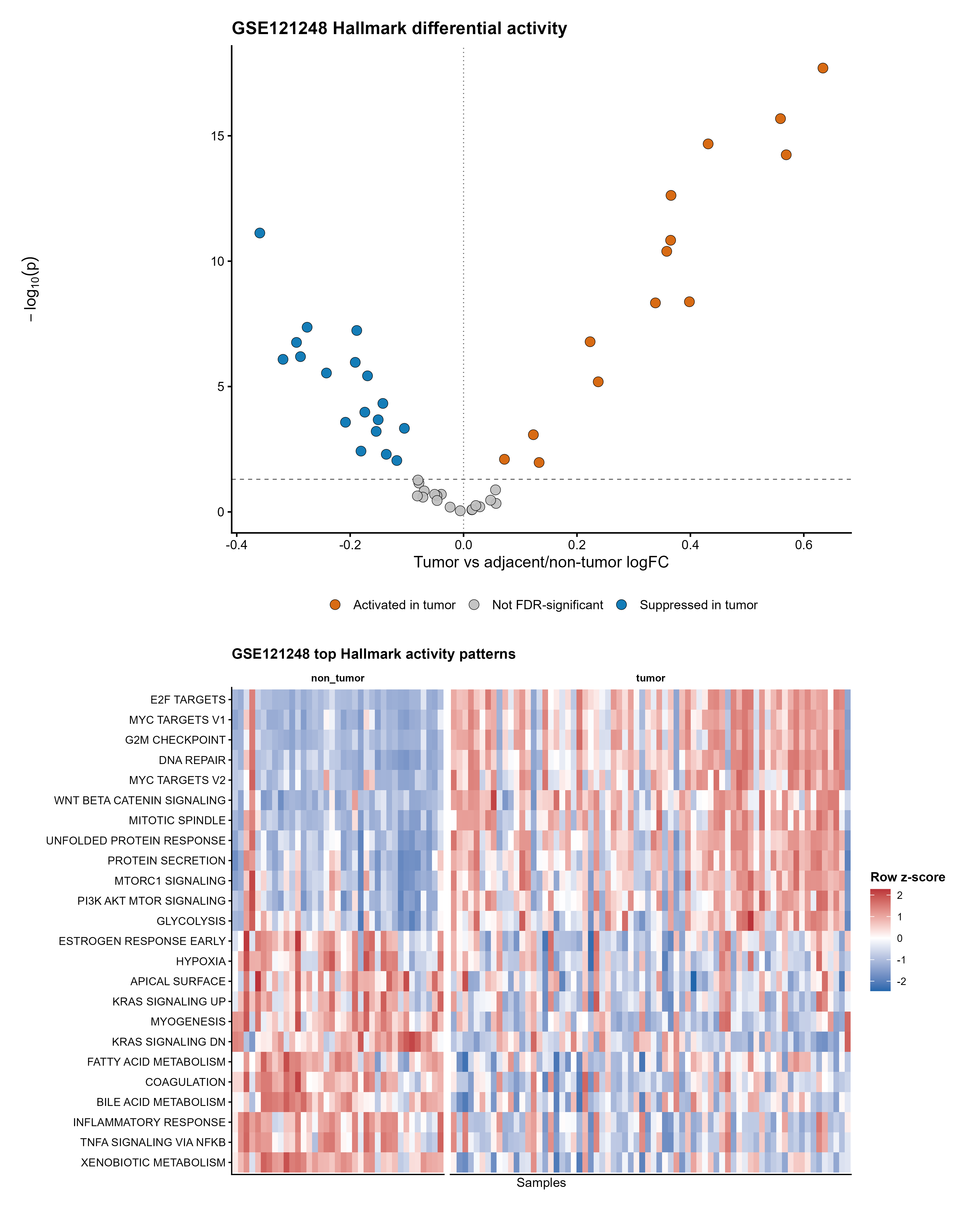

**Supplementary Figure S2. Hallmark pathway activity shifts in HBV-associated HCC.** Dataset-specific Hallmark GSVA results for GSE121248. Panel A shows a volcano plot of Hallmark differential activity for tumor versus adjacent/non-tumor tissue; each point represents one Hallmark gene set, the x-axis shows the tumor-associated effect size, and the y-axis shows -log10(p-value). Panel B shows a heatmap of the top-ranked Hallmark pathways across GSE121248 samples, with tissue annotation. The figure illustrates coordinated tumor-associated enrichment of proliferation/repair pathways and suppression of hepatocyte functional programs.

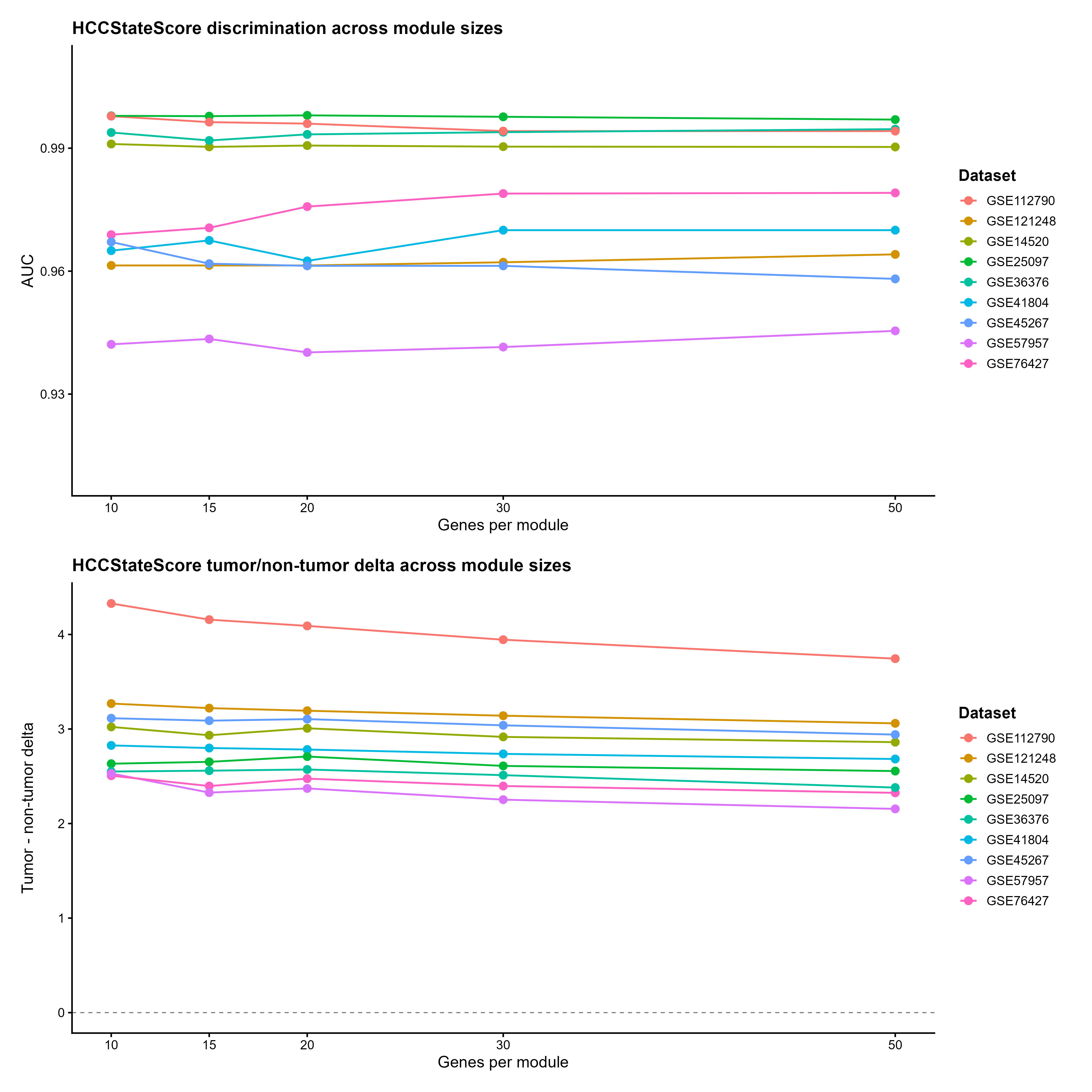

**Supplementary Figure S3. Module-size robustness of HCCStateScore across discovery and validation cohorts.** HCCStateScore tumor/non-tumor discrimination was evaluated across alternative module definitions using the top 10, 15, 20, 30, and 50 conserved genes per module. AUC values remained high across module sizes in discovery and validation cohorts, indicating that the score captures a stable proliferation-versus-hepatocyte-loss axis rather than a narrowly optimized top-20 gene list.

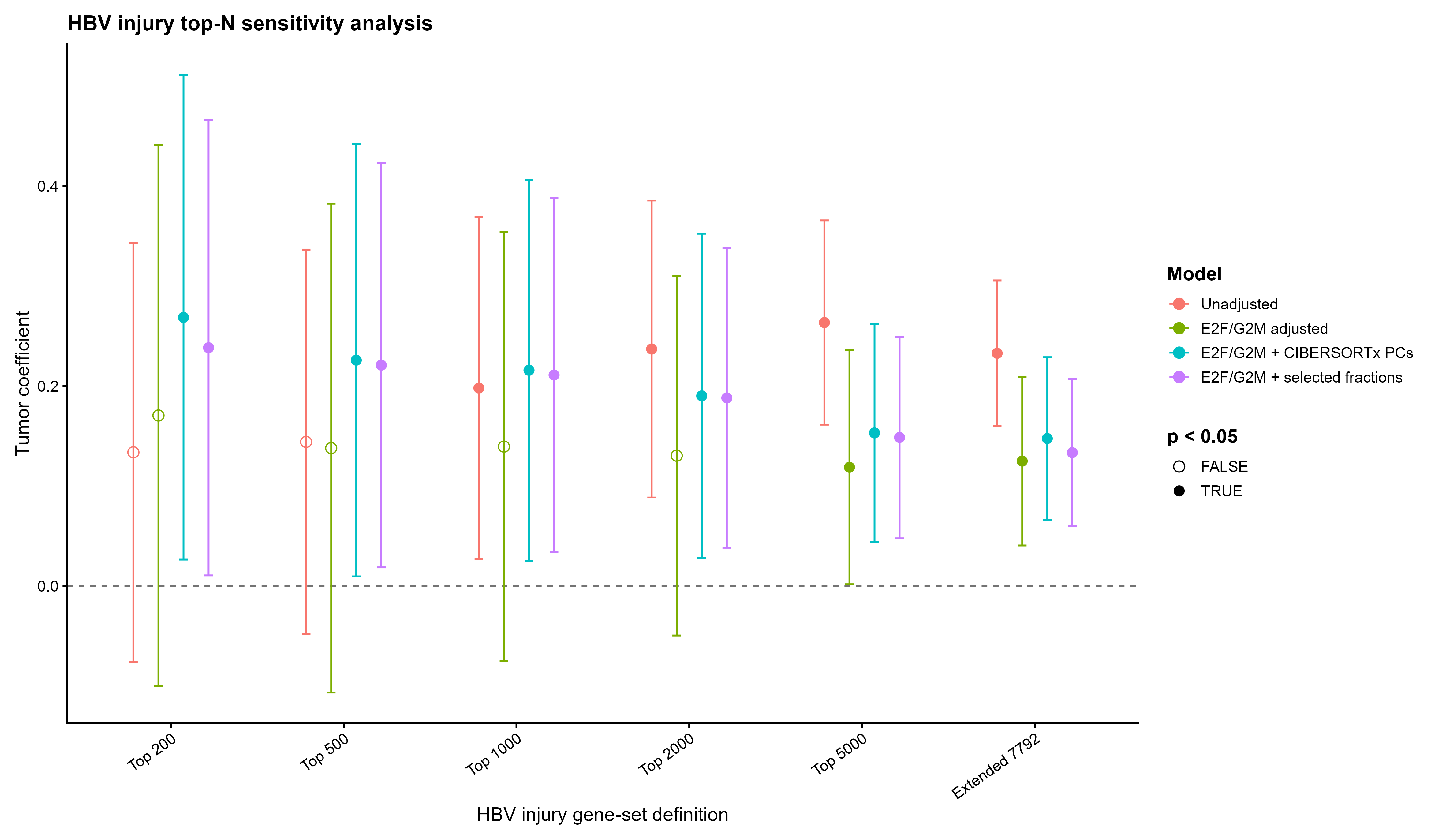

**Supplementary Figure S4. Top-N sensitivity analysis for the HBV injury program.** Tumor-associated HBV_INJURY coefficients in GSE121248 are shown across ranked injury-program sizes derived from the GSE83148 ALT/AST/HBV-DNA injury index. Very compact top-200 and top-500 definitions were less stable, whereas the top-2000 program provided a compact representation with positive tumor-associated effects and significant combined proliferation-plus-CIBERSORTx-adjusted associations. The top-5000 and extended FDR-defined 7,792-gene sets confirmed the same direction with stronger statistical robustness.

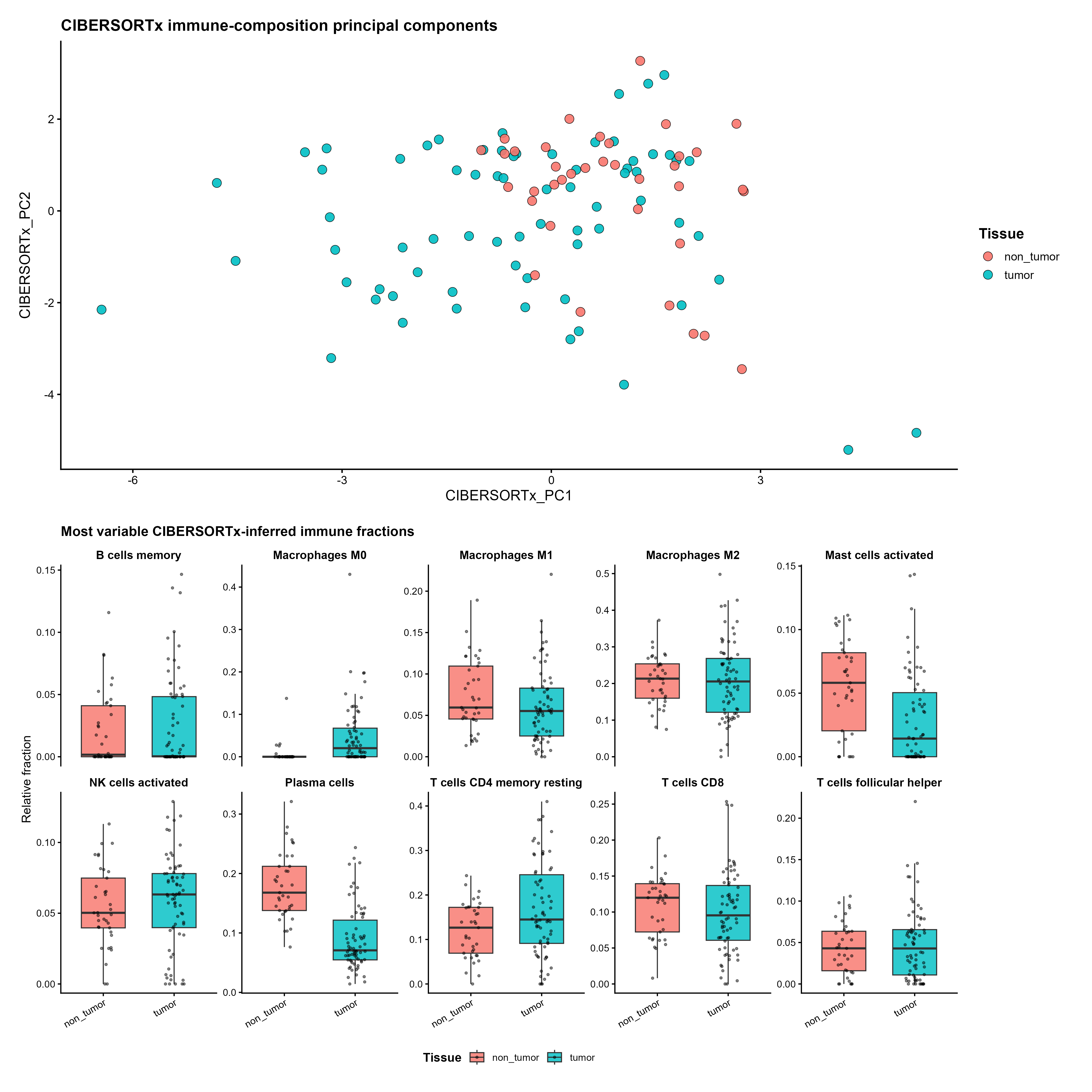
**Supplementary Figure S5. CIBERSORTx-inferred immune-composition structure in GSE121248.** CIBERSORTx-inferred immune-cell fractions were estimated from the GSE121248 expression mixture file using the LM22 signature matrix. Fractions were analyzed after compositional transformation and summarized by principal components for regression adjustment. The figure shows immune-composition structure by tissue group and supports the inclusion of CIBERSORTx-derived covariates in the HBV injury-axis models.

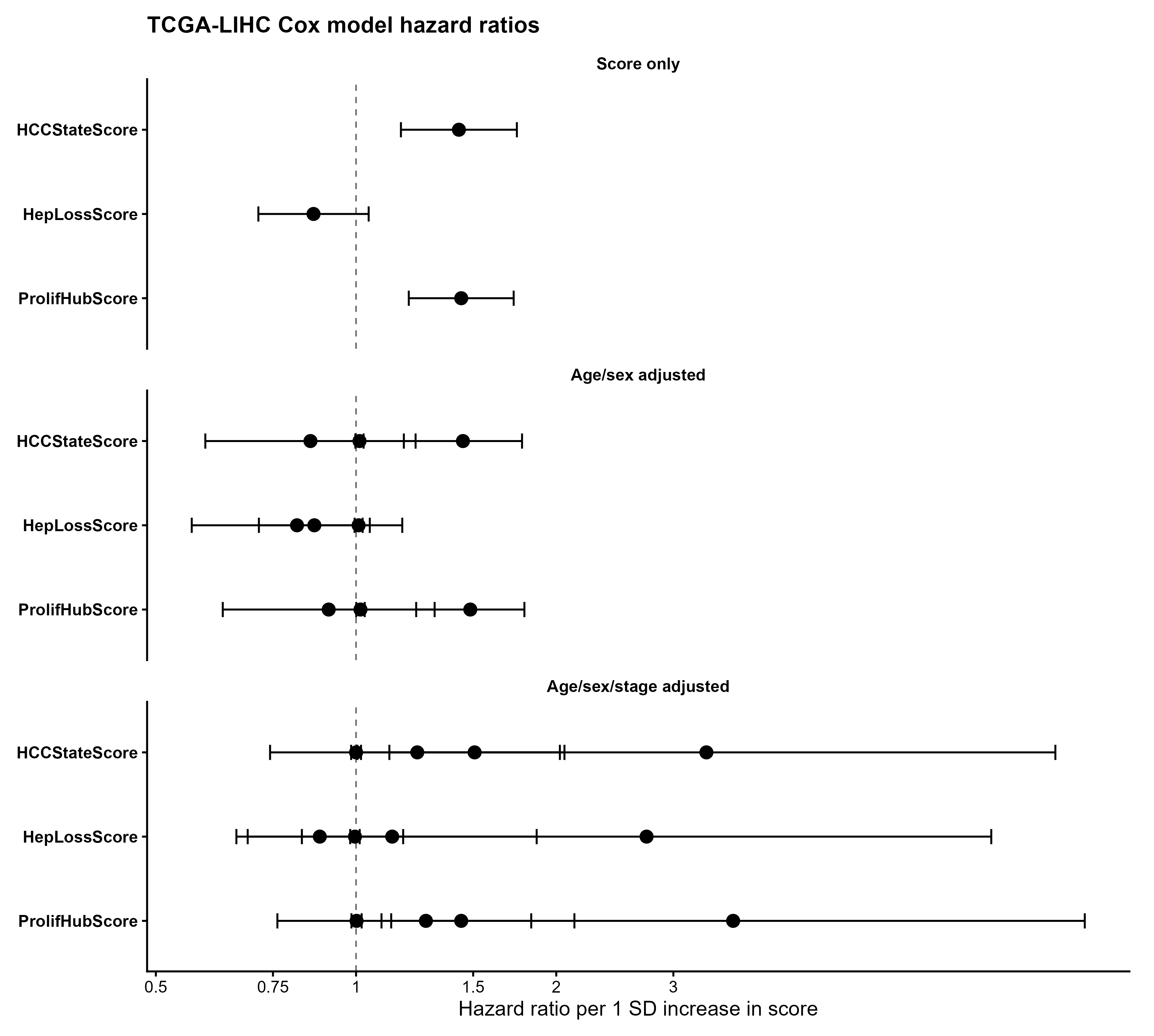

**Supplementary Figure S6. TCGA-LIHC Cox model forest plot for module scores.** Forest plot showing hazard ratios per one-standard-deviation increase in ProlifHubScore, HepLossScore, and HCCStateScore across score-only, age/sex-adjusted, and age/sex/stage-adjusted Cox proportional-hazards models. ProlifHubScore and HCCStateScore were associated with poorer overall survival, whereas HepLossScore showed a non-significant protective direction.
