## Supplementary Tables for "Cross-etiology transcriptomic conservation in hepatocellular carcinoma reveals opposing proliferation and hepatocyte-loss programs validated across cohorts"

**Supplementary Table S1. Dataset inventory, metadata curation, and inclusion decisions.** Cohort-level and sample-level curation summary for all public datasets considered in the study. The table reports GEO or TCGA accession, platform, associated publication, etiology when available, available tissue labels, numbers of tumor and adjacent/non-tumor samples, clinical or genotype variables parsed, use in the analysis, inclusion status, and reason for inclusion or exclusion. This table documents the dataset-selection process and supports the expanded multi-cohort validation design. Rows highlighted in yellow (GSE83148, GSE38941) do not contain conventional HCC tumor samples. GSE83148 = HBV chronic hepatitis (pre-neoplastic); GSE38941 = HBV-associated acute liver failure (ALF) vs. normal donor liver. Platform key: GPL570 = Affymetrix HG-U133 Plus 2.0 Array; GPL3921 = Affymetrix HT HG-U133A Array; GPL571 = Affymetrix HG-U133A 2.0 Array; GPL10558 = Illumina HumanHT-12 V4.0; GPL10687 = Rosetta/Merck Human RSTA Custom Affymetrix 1.0.

| **GEO Accession** | **Platform** | **n Tumor** | **n Non-Tumor** | **Etiology** | **Use in Study** | **Associated Publication / Citation** | **Reason for Inclusion** |
| --- | --- | --- | --- | --- | --- | --- | --- |
| GSE121248 | GPL570  (Affymetrix HG-U133 Plus 2.0) | 70 | 37 | HBV (chronic hepatitis B-induced HCC; adjacent normal tissue paired) | Discovery cohort; DEG identification across platforms | Wang SM et al. GEO submission 2018/2019. Gene expression profiling of chronic hepatitis B induced HCC and adjacent-normal tissues. (Singapore cohort) | Large HBV-specific HCC cohort; paired tumor/adjacent design; widely used as benchmark on Affymetrix GPL570; enables HBV-focused subgroup analyses |
| GSE41804 | GPL570  (Affymetrix HG-U133 Plus 2.0) | 20 | 20 | Not specified (general HCC) | Supplementary discovery cohort; DEG identification | Cited in: Spandidos Publ. (2017). Oncol Rep. 38(1):347–355. DOI: 10.3892/or.2017.5946 | Balanced paired design (20 tumor vs 20 adjacent non-tumor); same platform as GSE121248; adds independent replication for Affymetrix meta-analysis |
| GSE83148 | GPL570  (Affymetrix HG-U133 Plus 2.0) | 0 (HBV hepatitis samples, not HCC tumors) | 6 (healthy normal liver)  + 122 HBV-infected hepatitis | HBV (chronic hepatitis B; pre-neoplastic stage) | Pre-neoplastic HBV liver transcriptome reference; hepatitis-to-HCC progression context | Zhou et al. (2017); Li et al. (2018); Chen Z et al. (2019). 'Expression data of HBV infected liver tissue.' GEO submission. | Captures HBV-driven inflammatory milieu prior to overt HCC; provides hepatitis-stage reference for studying HBV disease continuum; same platform enables direct comparison with HCC datasets |
| GSE38941 | GPL570  (Affymetrix HG-U133 Plus 2.0) | 0 (not HCC)  17 HBV-ALF explanted livers | 10 (normal donor livers) | HBV-associated acute liver failure (ALF); 4 patients, massive or submassive hepatic necrosis | Reference for HBV-driven extreme liver injury; tumorigenesis gene signature in non-HCC context | Nissim O, Melis M, Diaz G, Kleiner DE, Tice A, Fantola G, Zamboni F, Mishra L, Farci P. (2012) PLoS One. 7(11):e49611. PMID: 23185381. PMC3504149. NIH/NIAID. | Not an HCC dataset per se — contains explanted HBV-ALF livers vs. normal donors. Notably, ALF shows strong tumorigenesis and cancer stem cell gene signatures (EpCAM, CK19, AKR1B10, TOP2A), making it relevant for studying pre-malignant transcriptional programs in HBV-damaged liver |
| GSE14520 | GPL3921 + GPL571  (Affymetrix HT HG-U133A / HG-U133A 2.0) | 247 total  (GPL3921: 225; GPL571: 22) | 241 total  (GPL3921: 220; GPL571: 21) | HBV (predominant; BCLC stages 0–C; Fudan University, Shanghai cohort) | Training/validation of prognostic signatures; benchmark for clinical annotation analyses | Roessler S et al. (2010). A Unique Metastasis Gene Signature Enables Prediction of Tumor Relapse in Early-Stage HCC Patients. Cancer Res. 70(24):10202–12. PMID: 21159642 | Largest HCC GEO microarray dataset; comprehensive clinical metadata (AFP, BCLC stage, survival); gold-standard benchmark; HBV-dominant cohort with full staging information |
| GSE25097 | GPL10687  (Rosetta/Merck Human RSTA Custom Affymetrix 1.0) | 268 | 289 | Mixed etiology (HBV, HCV, alcohol; US multicenter cohort) | Multi-etiology DEG discovery; training set for machine-learning biomarker models | Zhang C et al. GEO submission 2011. (Rosetta/Merck; NIH/NCI collaboration) | One of the largest HCC GEO expression datasets; multi-etiology cohort enabling pan-etiology signature derivation; unique Rosetta platform adds cross-platform diversity for robustness testing |
| GSE36376 | GPL10558  (Illumina HumanHT-12 V4.0) | 240 | 193 | HBV-predominant (Samsung Medical Center, Seoul, Korea; 2000–2006; Child-Pugh A) | DEG discovery; hub gene identification; cross-platform validation with Affymetrix datasets | Lim HY et al. (2012/2013). A Liver-Specific Long Noncoding RNA with a Role in Cell Viability is Conserved in Human, Mouse, and Rat Livers. Hepatology. 59(5):1871–84. PMID: 23913356 | Largest Illumina-based HCC microarray dataset; Korean HBV-endemic cohort; well-characterized clinical data; widely used as benchmark for Illumina-platform meta-analyses |
| GSE76427 | GPL10558  (Illumina HumanHT-12 V4.0) | 115 | 52 | Mixed (HBV/HCV/NASH; Singapore multicenter cohort) | DEG discovery; external validation of prognostic gene signatures | Grinchuk OV et al. (2018). Tumor-adjacent tissue co-expression profile analysis reveals pro-oncogenic ribosomal gene signature for prognosis of resectable HCC patients. Mol Oncol. 12(1):89–113. PMID: 29044954 | Paired design (52 matched tumor/adjacent non-tumor + 63 additional tumors); same Illumina platform as GSE36376 and GSE57957; Singapore multi-etiology cohort provides geographic diversity |
| GSE57957 | GPL10558  (Illumina HumanHT-12 V4.0) | 39 | 39 | Not specified (Singapore cohort) | DEG identification; multi-platform Illumina meta-analysis | Mah WC et al. (2014). Dosage effects of PITX2 on the transcriptome and epigenome of hepatocellular carcinoma cells. Epigenetics Chromatin. 7:13. PMID: 24995018 | Perfectly balanced paired design (1:1 tumor/adjacent); same Illumina platform as GSE36376/GSE76427; enables direct platform-controlled meta-analysis without batch correction artifacts |
| GSE45267 | GPL570  (Affymetrix HG-U133 Plus 2.0) | 48 | 39 | Not specified (general HCC) | DEG identification; validation of Affymetrix-derived gene signatures | Wang HM et al. GEO submission 2013. | Adds Affymetrix GPL570 cohort for cross-study overlap analysis; contributes to multi-cohort DEG consensus filtering alongside GSE121248; independently derived from different institution |
| GSE112790 | GPL570  (Affymetrix HG-U133 Plus 2.0) | 183 | 15 | Not specified (Japanese cohort, Shimada et al. 2019) | External validation of expression signatures; large tumor transcriptome reference set | Shimada S et al. (2019). YY1 complex promotes quaking expression via super-enhancer binding during EMT of hepatocellular carcinoma. Cancer Res. 79(7):1451–1464. PMID: 30679179 | Largest tumor sample count (n=183) among Affymetrix GPL570 datasets in this set; Japanese cohort adds geographic and genetic diversity; strong statistical power for validation despite low non-tumor n (15) |

**Supplementary Table S2. Complete Hallmark GSVA differential activity results.** Full Hallmark pathway differential activity results for the discovery cohorts GSE121248 and GSE41804. For each Hallmark gene set, the table reports cohort, tumor-versus-adjacent/non-tumor effect size, test statistic, nominal p-value, FDR-adjusted p-value, and direction of change. These data support the cross-etiology Hallmark pathway summary shown in the main text.

| dataset | hallmark gene set | logFC | t | p value | FDR |
| --- | --- | --- | --- | --- | --- |
| GSE121248 | HALLMARK_ADIPOGENESIS | -0.1177 | -2.6609 | 0.0089 | 0.0144 |
| GSE121248 | HALLMARK_ALLOGRAFT_REJECTION | -0.1809 | -2.9589 | 0.0038 | 0.0067 |
| GSE121248 | HALLMARK_ANDROGEN_RESPONSE | -0.0788 | -1.8274 | 0.0702 | 0.1033 |
| GSE121248 | HALLMARK_ANGIOGENESIS | 0.0147 | 0.2275 | 0.8205 | 0.8372 |
| GSE121248 | HALLMARK_APICAL_JUNCTION | 0.0564 | 1.5172 | 0.1320 | 0.1886 |
| GSE121248 | HALLMARK_APICAL_SURFACE | -0.1694 | -4.8624 | 3.738e-06 | 9.837e-06 |
| GSE121248 | HALLMARK_APOPTOSIS | -0.0694 | -1.4633 | 0.1461 | 0.2030 |
| GSE121248 | HALLMARK_BILE_ACID_METABOLISM | -0.2876 | -5.2739 | 6.413e-07 | 2.138e-06 |
| GSE121248 | HALLMARK_CHOLESTEROL_HOMEOSTASIS | -0.0237 | -0.4627 | 0.6445 | 0.6856 |
| GSE121248 | HALLMARK_COAGULATION | -0.2759 | -5.8723 | 4.315e-08 | 1.961e-07 |
| GSE121248 | HALLMARK_COMPLEMENT | -0.1363 | -2.8602 | 0.0050 | 0.0087 |
| GSE121248 | HALLMARK_DNA_REPAIR | 0.4313 | 9.1916 | 2.102e-15 | 3.504e-14 |
| GSE121248 | HALLMARK_E2F_TARGETS | 0.6337 | 10.4878 | 1.988e-18 | 9.940e-17 |
| GSE121248 | HALLMARK_EPITHELIAL_MESENCHYMAL_TRANSITION | -0.0717 | -1.1373 | 0.2578 | 0.3144 |
| GSE121248 | HALLMARK_ESTROGEN_RESPONSE_EARLY | -0.1423 | -4.2315 | 4.711e-05 | 1.122e-04 |
| GSE121248 | HALLMARK_ESTROGEN_RESPONSE_LATE | -0.1044 | -3.6054 | 4.635e-04 | 9.270e-04 |
| GSE121248 | HALLMARK_FATTY_ACID_METABOLISM | -0.2418 | -4.9240 | 2.886e-06 | 8.017e-06 |
| GSE121248 | HALLMARK_G2M_CHECKPOINT | 0.5589 | 9.6246 | 2.069e-16 | 5.172e-15 |
| GSE121248 | HALLMARK_GLYCOLYSIS | 0.1231 | 3.4330 | 8.324e-04 | 0.0015 |
| GSE121248 | HALLMARK_HEDGEHOG_SIGNALING | -0.0804 | -1.9494 | 0.0537 | 0.0814 |
| GSE121248 | HALLMARK_HEME_METABOLISM | -0.0393 | -1.2926 | 0.1988 | 0.2619 |
| GSE121248 | HALLMARK_HYPOXIA | -0.1506 | -3.8289 | 2.108e-04 | 4.582e-04 |
| GSE121248 | HALLMARK_IL2_STAT5_SIGNALING | -0.1541 | -3.5236 | 6.135e-04 | 0.0012 |
| GSE121248 | HALLMARK_IL6_JAK_STAT3_SIGNALING | -0.2082 | -3.7640 | 2.658e-04 | 5.538e-04 |
| GSE121248 | HALLMARK_INFLAMMATORY_RESPONSE | -0.2946 | -5.5677 | 1.737e-07 | 6.203e-07 |
| GSE121248 | HALLMARK_INTERFERON_ALPHA_RESPONSE | 0.0573 | 0.7441 | 0.4583 | 0.5208 |
| GSE121248 | HALLMARK_INTERFERON_GAMMA_RESPONSE | -0.0815 | -1.2056 | 0.2305 | 0.2881 |
| GSE121248 | HALLMARK_KRAS_SIGNALING_DN | -0.1910 | -5.1535 | 1.083e-06 | 3.186e-06 |
| GSE121248 | HALLMARK_KRAS_SIGNALING_UP | -0.1742 | -4.0188 | 1.053e-04 | 2.392e-04 |
| GSE121248 | HALLMARK_MITOTIC_SPINDLE | 0.3651 | 7.5042 | 1.466e-11 | 1.047e-10 |
| GSE121248 | HALLMARK_MTORC1_SIGNALING | 0.2375 | 4.7304 | 6.466e-06 | 1.616e-05 |
| GSE121248 | HALLMARK_MYC_TARGETS_V1 | 0.5690 | 9.0041 | 5.717e-15 | 7.147e-14 |
| GSE121248 | HALLMARK_MYC_TARGETS_V2 | 0.3983 | 6.3680 | 4.153e-09 | 2.304e-08 |
| GSE121248 | HALLMARK_MYOGENESIS | -0.1884 | -5.8079 | 5.809e-08 | 2.420e-07 |
| GSE121248 | HALLMARK_NOTCH_SIGNALING | 0.1332 | 2.5952 | 0.0107 | 0.0167 |
| GSE121248 | HALLMARK_OXIDATIVE_PHOSPHORYLATION | 0.0286 | 0.4896 | 0.6254 | 0.6798 |
| GSE121248 | HALLMARK_P53_PATHWAY | -0.0471 | -1.2163 | 0.2264 | 0.2881 |
| GSE121248 | HALLMARK_PANCREAS_BETA_CELLS | -0.0510 | -1.2917 | 0.1991 | 0.2619 |
| GSE121248 | HALLMARK_PEROXISOME | -0.0059 | -0.1265 | 0.8995 | 0.8995 |
| GSE121248 | HALLMARK_PI3K_AKT_MTOR_SIGNALING | 0.2232 | 5.5813 | 1.633e-07 | 6.203e-07 |
| GSE121248 | HALLMARK_PROTEIN_SECRETION | 0.3383 | 6.3464 | 4.608e-09 | 2.304e-08 |
| GSE121248 | HALLMARK_REACTIVE_OXYGEN_SPECIES_PATHWAY | 0.0477 | 0.9579 | 0.3402 | 0.4050 |
| GSE121248 | HALLMARK_SPERMATOGENESIS | 0.0721 | 2.7013 | 0.0080 | 0.0133 |
| GSE121248 | HALLMARK_TGF_BETA_SIGNALING | 0.0151 | 0.2470 | 0.8053 | 0.8372 |
| GSE121248 | HALLMARK_TNFA_SIGNALING_VIA_NFKB | -0.3184 | -5.2169 | 8.226e-07 | 2.571e-06 |
| GSE121248 | HALLMARK_UNFOLDED_PROTEIN_RESPONSE | 0.3583 | 7.3047 | 4.046e-11 | 2.529e-10 |
| GSE121248 | HALLMARK_UV_RESPONSE_DN | -0.0467 | -0.9358 | 0.3514 | 0.4086 |
| GSE121248 | HALLMARK_UV_RESPONSE_UP | 0.0217 | 0.5867 | 0.5586 | 0.6206 |
| GSE121248 | HALLMARK_WNT_BETA_CATENIN_SIGNALING | 0.3658 | 8.2991 | 2.382e-13 | 2.382e-12 |
| GSE121248 | HALLMARK_XENOBIOTIC_METABOLISM | -0.3593 | -7.6343 | 7.529e-12 | 6.274e-11 |
| GSE41804 | HALLMARK_ADIPOGENESIS | -0.2149 | -3.1065 | 0.0032 | 0.0201 |
| GSE41804 | HALLMARK_ALLOGRAFT_REJECTION | -0.0911 | -1.0430 | 0.3024 | 0.3779 |
| GSE41804 | HALLMARK_ANDROGEN_RESPONSE | -0.1251 | -1.8454 | 0.0713 | 0.1285 |
| GSE41804 | HALLMARK_ANGIOGENESIS | 0.0434 | 0.4653 | 0.6439 | 0.7154 |
| GSE41804 | HALLMARK_APICAL_JUNCTION | -0.0295 | -0.5301 | 0.5985 | 0.6960 |
| GSE41804 | HALLMARK_APICAL_SURFACE | -0.0467 | -0.7958 | 0.4302 | 0.5121 |
| GSE41804 | HALLMARK_APOPTOSIS | -0.0889 | -1.3937 | 0.1700 | 0.2298 |
| GSE41804 | HALLMARK_BILE_ACID_METABOLISM | -0.3144 | -3.9084 | 2.995e-04 | 0.0021 |
| GSE41804 | HALLMARK_CHOLESTEROL_HOMEOSTASIS | -0.1667 | -2.5017 | 0.0159 | 0.0613 |
| GSE41804 | HALLMARK_COAGULATION | -0.3200 | -4.6653 | 2.623e-05 | 2.623e-04 |
| GSE41804 | HALLMARK_COMPLEMENT | -0.1451 | -2.1611 | 0.0359 | 0.0951 |
| GSE41804 | HALLMARK_DNA_REPAIR | 0.1592 | 2.0426 | 0.0468 | 0.0963 |
| GSE41804 | HALLMARK_E2F_TARGETS | 0.4435 | 4.7254 | 2.149e-05 | 2.623e-04 |
| GSE41804 | HALLMARK_EPITHELIAL_MESENCHYMAL_TRANSITION | 0.0240 | 0.2573 | 0.7981 | 0.8654 |
| GSE41804 | HALLMARK_ESTROGEN_RESPONSE_EARLY | -0.1114 | -2.1254 | 0.0389 | 0.0951 |
| GSE41804 | HALLMARK_ESTROGEN_RESPONSE_LATE | -0.1323 | -2.8833 | 0.0059 | 0.0297 |
| GSE41804 | HALLMARK_FATTY_ACID_METABOLISM | -0.3302 | -4.2709 | 9.508e-05 | 7.924e-04 |
| GSE41804 | HALLMARK_G2M_CHECKPOINT | 0.4083 | 4.7780 | 1.804e-05 | 2.623e-04 |
| GSE41804 | HALLMARK_GLYCOLYSIS | -0.0102 | -0.1795 | 0.8583 | 0.8941 |
| GSE41804 | HALLMARK_HEDGEHOG_SIGNALING | -0.0544 | -0.8107 | 0.4217 | 0.5121 |
| GSE41804 | HALLMARK_HEME_METABOLISM | -0.0993 | -2.1200 | 0.0394 | 0.0951 |
| GSE41804 | HALLMARK_HYPOXIA | -0.1335 | -2.0296 | 0.0481 | 0.0963 |
| GSE41804 | HALLMARK_IL2_STAT5_SIGNALING | -0.1293 | -1.9815 | 0.0535 | 0.1028 |
| GSE41804 | HALLMARK_IL6_JAK_STAT3_SIGNALING | -0.1897 | -2.4345 | 0.0188 | 0.0671 |
| GSE41804 | HALLMARK_INFLAMMATORY_RESPONSE | -0.1582 | -2.1063 | 0.0406 | 0.0951 |
| GSE41804 | HALLMARK_INTERFERON_ALPHA_RESPONSE | -0.1836 | -1.6122 | 0.1137 | 0.1774 |
| GSE41804 | HALLMARK_INTERFERON_GAMMA_RESPONSE | -0.1950 | -2.0927 | 0.0419 | 0.0951 |
| GSE41804 | HALLMARK_KRAS_SIGNALING_DN | -0.1002 | -2.1167 | 0.0397 | 0.0951 |
| GSE41804 | HALLMARK_KRAS_SIGNALING_UP | -0.1033 | -1.5810 | 0.1206 | 0.1774 |
| GSE41804 | HALLMARK_MITOTIC_SPINDLE | 0.2725 | 5.2603 | 3.547e-06 | 8.868e-05 |
| GSE41804 | HALLMARK_MTORC1_SIGNALING | 0.1209 | 1.5870 | 0.1193 | 0.1774 |
| GSE41804 | HALLMARK_MYC_TARGETS_V1 | 0.2651 | 2.5779 | 0.0132 | 0.0549 |
| GSE41804 | HALLMARK_MYC_TARGETS_V2 | 0.1387 | 1.6675 | 0.1021 | 0.1647 |
| GSE41804 | HALLMARK_MYOGENESIS | -0.1193 | -2.0440 | 0.0466 | 0.0963 |
| GSE41804 | HALLMARK_NOTCH_SIGNALING | 0.0063 | 0.0919 | 0.9272 | 0.9272 |
| GSE41804 | HALLMARK_OXIDATIVE_PHOSPHORYLATION | -0.1882 | -1.7746 | 0.0825 | 0.1422 |
| GSE41804 | HALLMARK_P53_PATHWAY | -0.0927 | -1.7059 | 0.0947 | 0.1578 |
| GSE41804 | HALLMARK_PANCREAS_BETA_CELLS | -0.0257 | -0.4726 | 0.6387 | 0.7154 |
| GSE41804 | HALLMARK_PEROXISOME | -0.1954 | -2.7780 | 0.0079 | 0.0357 |
| GSE41804 | HALLMARK_PI3K_AKT_MTOR_SIGNALING | 0.1581 | 3.0130 | 0.0042 | 0.0232 |
| GSE41804 | HALLMARK_PROTEIN_SECRETION | 0.1175 | 1.4288 | 0.1597 | 0.2218 |
| GSE41804 | HALLMARK_REACTIVE_OXYGEN_SPECIES_PATHWAY | -0.1064 | -1.4885 | 0.1434 | 0.2048 |
| GSE41804 | HALLMARK_SPERMATOGENESIS | 0.0607 | 1.2988 | 0.2004 | 0.2637 |
| GSE41804 | HALLMARK_TGF_BETA_SIGNALING | -0.0167 | -0.2373 | 0.8135 | 0.8654 |
| GSE41804 | HALLMARK_TNFA_SIGNALING_VIA_NFKB | -0.2080 | -2.2386 | 0.0300 | 0.0938 |
| GSE41804 | HALLMARK_UNFOLDED_PROTEIN_RESPONSE | 0.1734 | 2.3167 | 0.0250 | 0.0832 |
| GSE41804 | HALLMARK_UV_RESPONSE_DN | -0.0089 | -0.1456 | 0.8849 | 0.9029 |
| GSE41804 | HALLMARK_UV_RESPONSE_UP | -0.0533 | -1.1026 | 0.2759 | 0.3537 |
| GSE41804 | HALLMARK_WNT_BETA_CATENIN_SIGNALING | 0.1146 | 1.8412 | 0.0720 | 0.1285 |
| GSE41804 | HALLMARK_XENOBIOTIC_METABOLISM | -0.4128 | -5.4848 | 1.646e-06 | 8.230e-05 |

**Supplementary Table S3. Full gene-level meta-analysis with heterogeneity statistics.** Complete gene-level meta-analysis output combining the GSE121248 and GSE41804 tumor-versus-non-tumor contrasts. The table reports cohort-specific log fold changes, standard errors and FDR values, fixed-effect meta-analysis estimates and p-values, FDR-adjusted meta-analysis results, direction concordance, I², REML-estimated τ², random-effects sensitivity estimates, and meta-analysis status. This table supports conserved-gene ranking and the heterogeneity summary reported in the main text. Preview shown: first 80 of 21753 rows. The full table is provided as a TSV file upon request.

| gene | meta status | meta logFC FE | meta se FE | meta z FE | meta p FE | meta FDR FE | Q | Q p | I2 | tau2 REML | tau2 FE | meta logFC REML | meta p REML | logFC 121248 | t 121248 | p 121248 | FDR 121248 | logFC 41804 | t 41804 | p 41804 | FDR 41804 | concordant direction | conserved | rank sum abs t |
| --- | --- | --- | --- | --- | --- | --- | --- | --- | --- | --- | --- | --- | --- | --- | --- | --- | --- | --- | --- | --- | --- | --- | --- | --- |
| A1BG | ok | -1.3617 | 0.2345 | -5.8068 | 6.369e-09 | 4.544e-08 | 3.2347 | 0.0721 | 69.0852 | 0.3406 | 0.0000 | -1.5426 | 0.0016 | -1.1271 | -4.2008 | 5.512e-05 | 2.170e-04 | -2.1201 | -4.3940 | 7.836e-05 | 0.0027 | TRUE | TRUE | 6145.0000 |
| A1BG-AS1 | ok | -0.1036 | 0.0351 | -2.9516 | 0.0032 | 0.0071 | 0.0283 | 0.8664 | 0.0000 | 0.0000 | 0.0000 | -0.1036 | 0.0032 | -0.1043 | -2.9509 | 0.0039 | 0.0092 | -0.0536 | -0.1792 | 0.8587 | 0.9346 | TRUE | FALSE | 2.921e+04 |
| A1CF | ok | -0.4085 | 0.1076 | -3.7969 | 1.465e-04 | 4.471e-04 | 0.0077 | 0.9301 | 0.0000 | 0.0000 | 0.0000 | -0.4085 | 1.465e-04 | -0.4198 | -2.4973 | 0.0140 | 0.0281 | -0.4006 | -2.8614 | 0.0066 | 0.0487 | TRUE | TRUE | 1.384e+04 |
| A2M | ok | -0.7927 | 0.1207 | -6.5661 | 5.164e-11 | 5.271e-10 | 0.1004 | 0.7514 | 0.0000 | 0.0000 | 0.0000 | -0.7927 | 5.164e-11 | -0.8177 | -5.6686 | 1.223e-07 | 9.600e-07 | -0.7342 | -3.3288 | 0.0019 | 0.0214 | TRUE | TRUE | 4673.0000 |
| A2M-AS1 | ok | -0.9896 | 0.1112 | -8.8955 | 5.817e-19 | 2.335e-17 | 3.9641 | 0.0465 | 74.7733 | 0.2383 | 0.0000 | -1.2379 | 0.0015 | -0.9225 | -7.9365 | 2.145e-12 | 6.743e-11 | -1.7209 | -4.4839 | 5.920e-05 | 0.0023 | TRUE | TRUE | 1255.0000 |
| A2ML1 | ok | -0.0657 | 0.0371 | -1.7717 | 0.0765 | 0.1178 | 5.5236 | 0.0188 | 81.8959 | 0.4292 | 0.0000 | -0.4788 | 0.3418 | -0.0582 | -1.5645 | 0.1206 | 0.1781 | -1.0820 | -2.4929 | 0.0169 | 0.0896 | TRUE | FALSE | 1.883e+04 |
| A2MP1 | ok | -0.1967 | 0.0516 | -3.8113 | 1.382e-04 | 4.240e-04 | 14.6214 | 1.314e-04 | 93.1607 | 0.9037 | 0.0000 | -0.8189 | 0.2386 | -0.1682 | -3.2247 | 0.0017 | 0.0044 | -1.5611 | -4.3300 | 9.555e-05 | 0.0031 | TRUE | TRUE | 8949.0000 |
| A4GALT | ok | -0.1602 | 0.0395 | -4.0549 | 5.016e-05 | 1.694e-04 | 0.1015 | 0.7501 | 0.0000 | 0.0000 | 0.0000 | -0.1602 | 5.016e-05 | -0.1586 | -3.9813 | 1.249e-04 | 4.462e-04 | -0.2583 | -0.8322 | 0.4102 | 0.6389 | TRUE | FALSE | 2.006e+04 |
| A4GNT | ok | -0.0208 | 0.0466 | -0.4470 | 0.6549 | 0.7193 | 7.4503 | 0.0063 | 86.5778 | 0.9475 | 0.0000 | -0.6517 | 0.3741 | -0.0098 | -0.2096 | 0.8344 | 0.8731 | -1.4892 | -2.7579 | 0.0087 | 0.0585 | TRUE | FALSE | 2.402e+04 |
| AA06 | ok | -0.0626 | 0.0375 | -1.6680 | 0.0953 | 0.1429 | 0.5822 | 0.4454 | 0.0000 | 0.0000 | 0.0000 | -0.0626 | 0.0953 | -0.0601 | -1.5971 | 0.1132 | 0.1686 | -0.3982 | -0.9021 | 0.3724 | 0.6052 | TRUE | FALSE | 2.798e+04 |
| AAAS | ok | 0.0949 | 0.0332 | 2.8604 | 0.0042 | 0.0092 | 1.1943 | 0.2745 | 16.2718 | 0.0020 | 0.0000 | 0.1064 | 0.0441 | 0.0861 | 2.5217 | 0.0131 | 0.0265 | 0.2443 | 1.7370 | 0.0900 | 0.2590 | TRUE | FALSE | 1.833e+04 |
| AACS | ok | 0.5015 | 0.1126 | 4.4554 | 8.375e-06 | 3.342e-05 | 3.3639 | 0.0666 | 70.2724 | 0.0801 | 0.0000 | 0.4173 | 0.0770 | 0.6204 | 4.7762 | 5.685e-06 | 2.906e-05 | 0.1430 | 0.6340 | 0.5297 | 0.7341 | TRUE | FALSE | 1.995e+04 |
| AACSP1 | ok | 0.0762 | 0.0786 | 0.9701 | 0.3320 | 0.4122 | 0.4583 | 0.4984 | 0.0000 | 0.0000 | 0.0000 | 0.0762 | 0.3320 | 0.0643 | 0.7978 | 0.4267 | 0.5121 | 0.3127 | 0.8734 | 0.3876 | 0.6182 | TRUE | FALSE | 3.177e+04 |
| AADAC | ok | -0.3886 | 0.1776 | -2.1881 | 0.0287 | 0.0501 | 1.5194 | 0.2177 | 34.1848 | 0.0345 | 0.0000 | -0.4060 | 0.0677 | -0.2149 | -0.9481 | 0.3452 | 0.4300 | -0.6645 | -2.3256 | 0.0251 | 0.1158 | TRUE | FALSE | 2.218e+04 |
| AADACL2 | ok | -0.0869 | 0.0336 | -2.5895 | 0.0096 | 0.0190 | 1.9977 | 0.1575 | 49.9434 | 0.1203 | 0.0000 | -0.2586 | 0.3908 | -0.0836 | -2.4866 | 0.0144 | 0.0288 | -0.7776 | -1.5875 | 0.1202 | 0.3109 | TRUE | FALSE | 1.932e+04 |
| AADACP1 | ok | -0.6299 | 0.1942 | -3.2426 | 0.0012 | 0.0029 | 1.4749 | 0.2246 | 32.1970 | 0.0688 | 0.0000 | -0.7027 | 0.0149 | -0.5292 | -2.5060 | 0.0137 | 0.0275 | -1.1828 | -2.3894 | 0.0216 | 0.1050 | TRUE | FALSE | 1.532e+04 |
| AADAT | ok | -1.6623 | 0.1530 | -10.8664 | 1.667e-27 | 2.484e-25 | 0.2294 | 0.6320 | 0.0000 | 0.0000 | 0.0000 | -1.6623 | 1.667e-27 | -1.7034 | -9.7100 | 2.238e-16 | 2.664e-14 | -1.5318 | -4.9014 | 1.582e-05 | 9.750e-04 | TRUE | TRUE | 535.0000 |
| AAED1 | ok | 0.3689 | 0.0902 | 4.0876 | 4.358e-05 | 1.492e-04 | 0.0245 | 0.8756 | 0.0000 | 0.0000 | 0.0000 | 0.3689 | 4.358e-05 | 0.3604 | 3.4328 | 8.496e-04 | 0.0024 | 0.3926 | 2.2248 | 0.0317 | 0.1345 | TRUE | FALSE | 1.270e+04 |
| AAGAB | ok | 0.3009 | 0.0606 | 4.9675 | 6.783e-07 | 3.359e-06 | 2.8411 | 0.0919 | 64.8024 | 0.0403 | 0.0000 | 0.3941 | 0.0197 | 0.2684 | 4.2208 | 5.109e-05 | 2.031e-04 | 0.6212 | 3.1148 | 0.0034 | 0.0313 | TRUE | TRUE | 7822.0000 |
| AAK1 | ok | 0.1640 | 0.0691 | 2.3738 | 0.0176 | 0.0325 | 6.3331 | 0.0119 | 84.2099 | 0.0762 | 0.0000 | 0.2672 | 0.2072 | 0.0738 | 0.9486 | 0.3450 | 0.4299 | 0.4993 | 3.3269 | 0.0019 | 0.0214 | TRUE | FALSE | 1.936e+04 |
| AAMDC | ok | 0.0593 | 0.0780 | 0.7604 | 0.4470 | 0.5268 | 6.8555 | 0.0088 | 85.4132 | 0.1139 | 0.0000 | -0.0755 | 0.7690 | 0.1597 | 1.8365 | 0.0690 | 0.1109 | -0.3569 | -2.0152 | 0.0506 | 0.1800 | FALSE | FALSE | 1.965e+04 |
| AAMP | ok | 0.0022 | 0.0434 | 0.0501 | 0.9600 | 0.9689 | 4.932e-04 | 0.9823 | 0.0000 | 0.0000 | 0.0000 | 0.0022 | 0.9600 | 0.0025 | 0.0545 | 0.9566 | 0.9680 | -7.947e-04 | -0.0057 | 0.9955 | 0.9982 | FALSE | FALSE | 4.319e+04 |
| AANAT | ok | -0.0892 | 0.0411 | -2.1726 | 0.0298 | 0.0518 | 0.0067 | 0.9347 | 0.0000 | 0.0000 | 0.0000 | -0.0892 | 0.0298 | -0.0889 | -2.1516 | 0.0337 | 0.0599 | -0.1207 | -0.3126 | 0.7562 | 0.8817 | TRUE | FALSE | 3.087e+04 |
| AAR2 | ok | 0.1718 | 0.0357 | 4.8104 | 1.506e-06 | 6.962e-06 | 1.2841 | 0.2571 | 22.1229 | 0.0032 | 0.0000 | 0.1551 | 0.0128 | 0.1820 | 4.9416 | 2.868e-06 | 1.576e-05 | 0.0107 | 0.0729 | 0.9422 | 0.9742 | TRUE | FALSE | 2.500e+04 |
| AARS | ok | 0.1219 | 0.0669 | 1.8213 | 0.0686 | 0.1073 | 0.2332 | 0.6291 | 0.0000 | 0.0000 | 0.0000 | 0.1219 | 0.0686 | 0.1355 | 1.8656 | 0.0648 | 0.1051 | 0.0454 | 0.2639 | 0.7932 | 0.9026 | TRUE | FALSE | 3.253e+04 |
| AARS2 | ok | 0.4122 | 0.0597 | 6.9055 | 5.003e-12 | 6.135e-11 | 3.9383 | 0.0472 | 74.6084 | 0.0393 | 0.0000 | 0.3295 | 0.0391 | 0.4636 | 7.1248 | 1.250e-10 | 2.295e-09 | 0.1392 | 0.9280 | 0.3589 | 0.5925 | TRUE | FALSE | 1.436e+04 |
| AASDH | ok | -0.0067 | 0.0539 | -0.1245 | 0.9009 | 0.9231 | 2.0736 | 0.1499 | 51.7753 | 0.0105 | 0.0000 | 0.0267 | 0.7809 | -0.0431 | -0.7242 | 0.4705 | 0.5543 | 0.1588 | 1.2509 | 0.2182 | 0.4434 | FALSE | FALSE | 2.917e+04 |
| AASDHPPT | ok | 0.0468 | 0.0580 | 0.8069 | 0.4197 | 0.5003 | 2.3898 | 0.1221 | 58.1556 | 0.0130 | 0.0000 | 0.0141 | 0.8910 | 0.0965 | 1.4547 | 0.1487 | 0.2126 | -0.1152 | -0.9616 | 0.3419 | 0.5765 | FALSE | FALSE | 2.811e+04 |
| AASS | ok | -0.7686 | 0.2312 | -3.3245 | 8.858e-04 | 0.0023 | 0.0826 | 0.7738 | 0.0000 | 0.0000 | 0.0000 | -0.7686 | 8.858e-04 | -0.7208 | -2.5298 | 0.0129 | 0.0260 | -0.8609 | -2.1759 | 0.0355 | 0.1446 | TRUE | FALSE | 1.608e+04 |
| AATBC | ok | -0.0307 | 0.0651 | -0.4707 | 0.6379 | 0.7038 | 0.1641 | 0.6854 | 0.0000 | 0.0000 | 0.0000 | -0.0307 | 0.6379 | -0.0225 | -0.3296 | 0.7424 | 0.7967 | -0.1157 | -0.5264 | 0.6015 | 0.7853 | TRUE | FALSE | 3.693e+04 |
| AATF | ok | 0.4326 | 0.0498 | 8.6890 | 3.656e-18 | 1.293e-16 | 1.0296 | 0.3103 | 2.8736 | 3.358e-04 | 0.0000 | 0.4309 | 1.792e-16 | 0.4517 | 8.4873 | 1.282e-13 | 6.143e-12 | 0.2988 | 2.1200 | 0.0402 | 0.1557 | TRUE | FALSE | 6069.0000 |
| AATK | ok | -0.2265 | 0.0523 | -4.3294 | 1.495e-05 | 5.641e-05 | 0.6282 | 0.4280 | 0.0000 | 0.0000 | 0.0000 | -0.2265 | 1.495e-05 | -0.2387 | -4.3769 | 2.803e-05 | 1.202e-04 | -0.0858 | -0.4640 | 0.6452 | 0.8149 | TRUE | FALSE | 2.229e+04 |
| ABAT | ok | -0.8041 | 0.1652 | -4.8676 | 1.129e-06 | 5.341e-06 | 0.3620 | 0.5474 | 0.0000 | 0.0000 | 0.0000 | -0.8041 | 1.129e-06 | -0.7342 | -3.6355 | 4.277e-04 | 0.0013 | -0.9454 | -3.2923 | 0.0021 | 0.0229 | TRUE | TRUE | 8991.0000 |
| ABCA1 | ok | -0.3127 | 0.0760 | -4.1162 | 3.852e-05 | 1.338e-04 | 1.4664 | 0.2259 | 31.8067 | 0.0068 | 0.0000 | -0.3276 | 8.474e-04 | -0.2563 | -2.8773 | 0.0048 | 0.0111 | -0.4628 | -3.1829 | 0.0028 | 0.0277 | TRUE | TRUE | 1.170e+04 |
| ABCA11P | ok | 0.0274 | 0.0856 | 0.3195 | 0.7493 | 0.8003 | 0.3024 | 0.5824 | 0.0000 | 0.0000 | 0.0000 | 0.0274 | 0.7493 | 0.0522 | 0.5392 | 0.5909 | 0.6664 | -0.0619 | -0.3372 | 0.7377 | 0.8713 | FALSE | FALSE | 3.770e+04 |
| ABCA12 | ok | 0.0221 | 0.0302 | 0.7321 | 0.4641 | 0.5437 | 0.4035 | 0.5253 | 0.0000 | 0.0000 | 0.0000 | 0.0221 | 0.4641 | 0.0209 | 0.6898 | 0.4918 | 0.5748 | 0.3204 | 0.6809 | 0.4998 | 0.7116 | TRUE | FALSE | 3.389e+04 |
| ABCA13 | ok | 0.0763 | 0.1144 | 0.6666 | 0.5050 | 0.5831 | 0.3191 | 0.5722 | 0.0000 | 0.0000 | 0.0000 | 0.0763 | 0.5050 | 0.0888 | 0.7619 | 0.4478 | 0.5327 | -0.2571 | -0.4276 | 0.6712 | 0.8319 | FALSE | FALSE | 3.584e+04 |
| ABCA17P | ok | 0.0600 | 0.0439 | 1.3674 | 0.1715 | 0.2365 | 3.7250 | 0.0536 | 73.1545 | 0.1987 | 0.0000 | -0.2024 | 0.5693 | 0.0698 | 1.5813 | 0.1167 | 0.1731 | -0.6673 | -1.7590 | 0.0861 | 0.2527 | FALSE | FALSE | 2.208e+04 |
| ABCA2 | ok | -0.2156 | 0.0498 | -4.3270 | 1.511e-05 | 5.694e-05 | 3.7433 | 0.0530 | 73.2854 | 0.1461 | 0.0000 | -0.4359 | 0.1534 | -0.2005 | -3.9761 | 1.273e-04 | 4.537e-04 | -0.8320 | -2.5802 | 0.0136 | 0.0780 | TRUE | FALSE | 9901.0000 |
| ABCA3 | ok | -0.0106 | 0.0514 | -0.2062 | 0.8367 | 0.8716 | 1.7159 | 0.1902 | 41.7224 | 0.0372 | 0.0000 | -0.0941 | 0.5929 | 4.210e-04 | 0.0081 | 0.9936 | 0.9955 | -0.4217 | -1.3260 | 0.1923 | 0.4120 | FALSE | FALSE | 3.186e+04 |
| ABCA4 | ok | 0.0304 | 0.0740 | 0.4109 | 0.6811 | 0.7426 | 0.5428 | 0.4613 | 0.0000 | 0.0000 | 0.0000 | 0.0304 | 0.6811 | 0.0218 | 0.2908 | 0.7717 | 0.8214 | 0.3750 | 0.7919 | 0.4331 | 0.6581 | TRUE | FALSE | 3.475e+04 |
| ABCA5 | ok | -0.0270 | 0.1119 | -0.2413 | 0.8093 | 0.8496 | 0.0130 | 0.9094 | 0.0000 | 0.0000 | 0.0000 | -0.0270 | 0.8093 | -0.0162 | -0.1104 | 0.9123 | 0.9344 | -0.0420 | -0.2429 | 0.8093 | 0.9105 | TRUE | FALSE | 4.057e+04 |
| ABCA6 | ok | -0.5066 | 0.2095 | -2.4180 | 0.0156 | 0.0292 | 0.0768 | 0.7817 | 0.0000 | 0.0000 | 0.0000 | -0.5066 | 0.0156 | -0.4696 | -1.8917 | 0.0612 | 0.1002 | -0.5979 | -1.5314 | 0.1335 | 0.3314 | TRUE | FALSE | 2.206e+04 |
| ABCA7 | ok | -0.0355 | 0.0541 | -0.6558 | 0.5120 | 0.5897 | 1.6651 | 0.1969 | 39.9448 | 0.0140 | 0.0000 | -0.0805 | 0.4801 | -0.0156 | -0.2777 | 0.7818 | 0.8300 | -0.2807 | -1.4206 | 0.1631 | 0.3742 | TRUE | FALSE | 2.997e+04 |
| ABCA8 | ok | -1.6415 | 0.2862 | -5.7352 | 9.742e-09 | 6.730e-08 | 0.1364 | 0.7119 | 0.0000 | 0.0000 | 0.0000 | -1.6415 | 9.742e-09 | -1.5957 | -5.1159 | 1.375e-06 | 8.166e-06 | -1.8856 | -2.6183 | 0.0124 | 0.0736 | TRUE | FALSE | 7319.0000 |
| ABCA9 | ok | -0.8608 | 0.1657 | -5.1936 | 2.063e-07 | 1.129e-06 | 0.0676 | 0.7948 | 0.0000 | 0.0000 | 0.0000 | -0.8608 | 2.063e-07 | -0.8440 | -4.7459 | 6.437e-06 | 3.246e-05 | -0.9715 | -2.1255 | 0.0397 | 0.1545 | TRUE | FALSE | 9902.0000 |
| ABCB1 | ok | -0.2440 | 0.1714 | -1.4233 | 0.1546 | 0.2163 | 0.0053 | 0.9421 | 0.0000 | 0.0000 | 0.0000 | -0.2440 | 0.1546 | -0.2589 | -0.9650 | 0.3367 | 0.4213 | -0.2336 | -1.0488 | 0.3005 | 0.5348 | TRUE | FALSE | 2.961e+04 |
| ABCB10 | ok | 0.3409 | 0.0785 | 4.3405 | 1.422e-05 | 5.396e-05 | 1.2376 | 0.2659 | 19.1957 | 0.0041 | 0.0000 | 0.3303 | 3.946e-04 | 0.3891 | 4.3381 | 3.258e-05 | 1.371e-04 | 0.1825 | 1.1218 | 0.2686 | 0.5003 | TRUE | FALSE | 1.684e+04 |
| ABCB11 | ok | -0.0591 | 0.2565 | -0.2303 | 0.8179 | 0.8568 | 1.5685 | 0.2104 | 36.2430 | 0.1418 | 0.0000 | -0.1693 | 0.6703 | 0.0792 | 0.2837 | 0.7772 | 0.8261 | -0.8055 | -1.2414 | 0.2216 | 0.4482 | FALSE | FALSE | 3.122e+04 |
| ABCB4 | ok | -0.8257 | 0.1823 | -4.5307 | 5.878e-06 | 2.416e-05 | 1.7376 | 0.1874 | 42.4487 | 0.0492 | 0.0000 | -0.8323 | 5.407e-04 | -0.6005 | -2.4038 | 0.0179 | 0.0347 | -1.0820 | -4.0604 | 2.186e-04 | 0.0052 | TRUE | TRUE | 1.215e+04 |
| ABCB5 | ok | 0.0201 | 0.0475 | 0.4242 | 0.6714 | 0.7343 | 0.3966 | 0.5288 | 0.0000 | 0.0000 | 0.0000 | 0.0201 | 0.6714 | 0.0230 | 0.4821 | 0.6307 | 0.7021 | -0.2931 | -0.5866 | 0.5607 | 0.7572 | FALSE | FALSE | 3.565e+04 |
| ABCB6 | ok | 0.3308 | 0.1051 | 3.1488 | 0.0016 | 0.0039 | 1.5491 | 0.2133 | 35.4451 | 0.0195 | 0.0000 | 0.3671 | 0.0160 | 0.2671 | 2.2850 | 0.0243 | 0.0452 | 0.5991 | 2.4985 | 0.0166 | 0.0887 | TRUE | FALSE | 1.576e+04 |
| ABCB7 | ok | -0.1309 | 0.0554 | -2.3636 | 0.0181 | 0.0334 | 0.1161 | 0.7333 | 0.0000 | 0.0000 | 0.0000 | -0.1309 | 0.0181 | -0.1421 | -2.2077 | 0.0294 | 0.0533 | -0.0990 | -0.9105 | 0.3680 | 0.6013 | TRUE | FALSE | 2.530e+04 |
| ABCB8 | ok | 0.2243 | 0.0450 | 4.9794 | 6.379e-07 | 3.174e-06 | 1.3595 | 0.2436 | 26.4423 | 0.0038 | 0.0000 | 0.2419 | 4.804e-04 | 0.2061 | 4.3236 | 3.446e-05 | 1.437e-04 | 0.3759 | 2.7314 | 0.0093 | 0.0612 | TRUE | FALSE | 8523.0000 |
| ABCB9 | ok | -0.0529 | 0.0403 | -1.3127 | 0.1893 | 0.2574 | 0.3647 | 0.5459 | 0.0000 | 0.0000 | 0.0000 | -0.0529 | 0.1893 | -0.0505 | -1.2479 | 0.2148 | 0.2898 | -0.3025 | -0.7284 | 0.4706 | 0.6890 | TRUE | FALSE | 3.098e+04 |
| ABCC1 | ok | -0.0054 | 0.1478 | -0.0362 | 0.9711 | 0.9778 | 0.8855 | 0.3467 | 0.0000 | 0.0000 | 0.0000 | -0.0054 | 0.9711 | 0.0717 | 0.4242 | 0.6722 | 0.7379 | -0.2564 | -0.8407 | 0.4054 | 0.6345 | FALSE | FALSE | 3.372e+04 |
| ABCC10 | ok | 0.6230 | 0.0853 | 7.3032 | 2.810e-13 | 4.213e-12 | 5.7359 | 0.0166 | 82.5661 | 0.1053 | 0.0000 | 0.5005 | 0.0463 | 0.7271 | 7.5943 | 1.208e-11 | 3.055e-10 | 0.2222 | 1.1830 | 0.2437 | 0.4739 | TRUE | FALSE | 1.205e+04 |
| ABCC11 | ok | -0.0149 | 0.0465 | -0.3203 | 0.7487 | 0.7998 | 0.2958 | 0.5865 | 0.0000 | 0.0000 | 0.0000 | -0.0149 | 0.7487 | -0.0119 | -0.2538 | 0.8001 | 0.8451 | -0.2272 | -0.5779 | 0.5665 | 0.7603 | TRUE | FALSE | 3.680e+04 |
| ABCC12 | ok | -0.0196 | 0.0496 | -0.3950 | 0.6928 | 0.7528 | 2.8819 | 0.0896 | 65.3007 | 0.2482 | 0.0000 | -0.2989 | 0.4659 | -0.0114 | -0.2283 | 0.8198 | 0.8617 | -0.8833 | -1.7279 | 0.0916 | 0.2619 | TRUE | FALSE | 2.830e+04 |
| ABCC13 | ok | -0.0985 | 0.0522 | -1.8885 | 0.0590 | 0.0940 | 1.2562 | 0.2624 | 20.3961 | 0.0216 | 0.0000 | -0.1439 | 0.3268 | -0.0910 | -1.7291 | 0.0867 | 0.1346 | -0.5508 | -1.3537 | 0.1833 | 0.4000 | TRUE | FALSE | 2.398e+04 |
| ABCC2 | ok | -0.3417 | 0.1360 | -2.5130 | 0.0120 | 0.0231 | 0.1407 | 0.7076 | 0.0000 | 0.0000 | 0.0000 | -0.3417 | 0.0120 | -0.3949 | -2.0098 | 0.0470 | 0.0800 | -0.2928 | -1.5544 | 0.1279 | 0.3226 | TRUE | FALSE | 2.140e+04 |
| ABCC3 | ok | -0.2422 | 0.1399 | -1.7310 | 0.0834 | 0.1272 | 1.5165 | 0.2181 | 34.0603 | 0.0272 | 0.0000 | -0.2766 | 0.1418 | -0.1435 | -0.8899 | 0.3755 | 0.4610 | -0.5429 | -1.9290 | 0.0608 | 0.2027 | TRUE | FALSE | 2.424e+04 |
| ABCC4 | ok | 0.8206 | 0.1674 | 4.9026 | 9.458e-07 | 4.542e-06 | 0.0554 | 0.8139 | 0.0000 | 0.0000 | 0.0000 | 0.8206 | 9.458e-07 | 0.8433 | 4.3626 | 2.962e-05 | 1.261e-04 | 0.7524 | 2.2491 | 0.0300 | 0.1297 | TRUE | FALSE | 1.014e+04 |
| ABCC5 | ok | 0.4966 | 0.0681 | 7.2917 | 3.060e-13 | 4.556e-12 | 3.8184 | 0.0507 | 73.8110 | 0.0593 | 0.0000 | 0.6072 | 0.0020 | 0.4461 | 6.1222 | 1.537e-08 | 1.527e-07 | 0.8469 | 4.4166 | 7.304e-05 | 0.0026 | TRUE | TRUE | 2793.0000 |
| ABCC6P1 | ok | -0.1978 | 0.1722 | -1.1484 | 0.2508 | 0.3261 | 0.0054 | 0.9417 | 0.0000 | 0.0000 | 0.0000 | -0.1978 | 0.2508 | -0.1855 | -0.7718 | 0.4419 | 0.5270 | -0.2108 | -0.8536 | 0.3984 | 0.6274 | TRUE | FALSE | 3.205e+04 |
| ABCC8 | ok | -0.1227 | 0.0393 | -3.1246 | 0.0018 | 0.0042 | 0.0123 | 0.9117 | 0.0000 | 0.0000 | 0.0000 | -0.1227 | 0.0018 | -0.1232 | -3.1153 | 0.0024 | 0.0059 | -0.0867 | -0.2652 | 0.7922 | 0.9021 | TRUE | FALSE | 2.775e+04 |
| ABCC9 | ok | 0.0292 | 0.1200 | 0.2432 | 0.8078 | 0.8486 | 3.6927 | 0.0546 | 72.9199 | 0.1866 | 0.0000 | -0.1703 | 0.6265 | 0.1133 | 0.8876 | 0.3767 | 0.4622 | -0.6021 | -1.7216 | 0.0928 | 0.2641 | FALSE | FALSE | 2.537e+04 |
| ABCD1 | ok | 0.0729 | 0.0497 | 1.4663 | 0.1426 | 0.2016 | 0.5297 | 0.4667 | 0.0000 | 0.0000 | 0.0000 | 0.0729 | 0.1426 | 0.0836 | 1.6125 | 0.1098 | 0.1644 | -0.0494 | -0.2819 | 0.7795 | 0.8955 | FALSE | FALSE | 3.346e+04 |
| ABCD2 | ok | -0.0714 | 0.0288 | -2.4816 | 0.0131 | 0.0250 | 0.0083 | 0.9274 | 0.0000 | 0.0000 | 0.0000 | -0.0714 | 0.0131 | -0.0713 | -2.4728 | 0.0150 | 0.0297 | -0.1190 | -0.2274 | 0.8212 | 0.9168 | TRUE | FALSE | 3.044e+04 |
| ABCD3 | ok | -0.3517 | 0.0722 | -4.8729 | 1.100e-06 | 5.208e-06 | 7.2536 | 0.0071 | 86.2137 | 0.0700 | 0.0000 | -0.3976 | 0.0484 | -0.2034 | -2.2402 | 0.0271 | 0.0498 | -0.6064 | -5.0971 | 8.458e-06 | 6.698e-04 | TRUE | TRUE | 1.213e+04 |
| ABCD4 | ok | -0.2817 | 0.0647 | -4.3508 | 1.356e-05 | 5.171e-05 | 2.5911 | 0.1075 | 61.4065 | 0.0239 | 0.0000 | -0.3387 | 0.0120 | -0.2349 | -3.3111 | 0.0013 | 0.0034 | -0.5140 | -3.2492 | 0.0023 | 0.0247 | TRUE | TRUE | 1.004e+04 |
| ABCE1 | ok | -0.0611 | 0.0571 | -1.0709 | 0.2842 | 0.3617 | 0.3810 | 0.5371 | 0.0000 | 0.0000 | 0.0000 | -0.0611 | 0.2842 | -0.0432 | -0.6748 | 0.5012 | 0.5834 | -0.1305 | -1.0356 | 0.3065 | 0.5411 | TRUE | FALSE | 3.101e+04 |
| ABCF1 | ok | 0.3564 | 0.0528 | 6.7473 | 1.507e-11 | 1.689e-10 | 0.2722 | 0.6018 | 0.0000 | 0.0000 | 0.0000 | 0.3564 | 1.507e-11 | 0.3685 | 6.3886 | 4.401e-09 | 5.139e-08 | 0.2936 | 2.2323 | 0.0312 | 0.1330 | TRUE | FALSE | 6966.0000 |
| ABCF2 | ok | 0.5945 | 0.0688 | 8.6382 | 5.710e-18 | 1.953e-16 | 0.0631 | 0.8016 | 0.0000 | 0.0000 | 0.0000 | 0.5945 | 5.710e-18 | 0.6025 | 7.9372 | 2.138e-12 | 6.739e-11 | 0.5573 | 3.4181 | 0.0015 | 0.0181 | TRUE | TRUE | 2429.0000 |
| ABCF3 | ok | -0.1066 | 0.0422 | -2.5235 | 0.0116 | 0.0225 | 2.0785 | 0.1494 | 51.8873 | 0.0130 | 0.0000 | -0.1553 | 0.1294 | -0.0886 | -2.0109 | 0.0468 | 0.0798 | -0.3124 | -2.0982 | 0.0422 | 0.1605 | TRUE | FALSE | 1.848e+04 |
| ABCG1 | ok | 0.2744 | 0.1537 | 1.7852 | 0.0742 | 0.1149 | 0.4272 | 0.5134 | 0.0000 | 0.0000 | 0.0000 | 0.2744 | 0.0742 | 0.3811 | 1.6998 | 0.0921 | 0.1415 | 0.1798 | 0.8511 | 0.3997 | 0.6286 | TRUE | FALSE | 2.798e+04 |
| ABCG2 | ok | -0.8501 | 0.1901 | -4.4728 | 7.721e-06 | 3.104e-05 | 0.0182 | 0.8927 | 0.0000 | 0.0000 | 0.0000 | -0.8501 | 7.721e-06 | -0.8328 | -3.6346 | 4.290e-04 | 0.0013 | -0.8881 | -2.6103 | 0.0126 | 0.0745 | TRUE | FALSE | 1.071e+04 |
| ABCG4 | ok | -0.1169 | 0.0404 | -2.8936 | 0.0038 | 0.0084 | 0.0552 | 0.8142 | 0.0000 | 0.0000 | 0.0000 | -0.1169 | 0.0038 | -0.1177 | -2.9031 | 0.0045 | 0.0104 | -0.0027 | -0.0055 | 0.9956 | 0.9983 | TRUE | FALSE | 3.110e+04 |
| ABCG5 | ok | -0.7117 | 0.1925 | -3.6978 | 2.175e-04 | 6.393e-04 | 0.0319 | 0.8583 | 0.0000 | 0.0000 | 0.0000 | -0.7117 | 2.175e-04 | -0.6775 | -2.4941 | 0.0142 | 0.0283 | -0.7462 | -2.7359 | 0.0092 | 0.0607 | TRUE | FALSE | 1.418e+04 |
| ABCG8 | ok | -0.3647 | 0.1358 | -2.6850 | 0.0073 | 0.0148 | 0.7005 | 0.4026 | 0.0000 | 0.0000 | 0.0000 | -0.3647 | 0.0073 | -0.3060 | -2.0017 | 0.0478 | 0.0812 | -0.5849 | -1.9755 | 0.0551 | 0.1896 | TRUE | FALSE | 1.913e+04 |

**Supplementary Table S4. ProlifHub and HepLoss module definitions.** Gene membership and derivation statistics for the ProlifHub and HepLoss modules. For each module gene, the table reports module assignment, rank, gene symbol, direction of change, discovery-cohort statistics, meta-analysis estimates, FDR values, and heterogeneity metrics. This table documents how the conserved tumor-upregulated and tumor-downregulated panels were generated.

| module | gene | rank | module size | meta logFC FE | meta FDR FE | I2 | tau2 REML |
| --- | --- | --- | --- | --- | --- | --- | --- |
| ProlifHub | CAP2 | 1.0000 | 20.0000 | 2.2466 | 1.092e-51 | 78.1434 | 0.1637 |
| ProlifHub | ASPM | 2.0000 | 20.0000 | 2.6756 | 7.128e-41 | 48.9355 | 0.0823 |
| ProlifHub | CENPW | 3.0000 | 20.0000 | 1.6770 | 1.760e-35 | 0.0000 | 0.0000 |
| ProlifHub | PLVAP | 4.0000 | 20.0000 | 1.3862 | 6.306e-48 | 74.6441 | 0.1928 |
| ProlifHub | TOP2A | 5.0000 | 20.0000 | 3.1002 | 6.327e-43 | 40.7965 | 0.0883 |
| ProlifHub | ECT2 | 6.0000 | 20.0000 | 2.0613 | 2.166e-33 | 0.0000 | 0.0000 |
| ProlifHub | NUSAP1 | 7.0000 | 20.0000 | 1.8479 | 2.627e-34 | 0.0000 | 0.0000 |
| ProlifHub | ITGA6 | 8.0000 | 20.0000 | 1.3238 | 1.573e-32 | 0.3990 | 1.089e-04 |
| ProlifHub | RACGAP1 | 9.0000 | 20.0000 | 1.8449 | 1.640e-38 | 70.1232 | 0.1085 |
| ProlifHub | TTC13 | 10.0000 | 20.0000 | 0.8860 | 2.181e-30 | 0.0000 | 0.0000 |
| ProlifHub | COL15A1 | 11.0000 | 20.0000 | 2.8919 | 8.663e-37 | 0.0000 | 0.0000 |
| ProlifHub | GJC1 | 12.0000 | 20.0000 | 1.5934 | 8.200e-30 | 59.6550 | 0.0883 |
| ProlifHub | PRC1 | 13.0000 | 20.0000 | 2.0476 | 2.742e-32 | 0.0000 | 0.0000 |
| ProlifHub | CDKN3 | 14.0000 | 20.0000 | 1.9604 | 4.441e-32 | 0.0000 | 0.0000 |
| ProlifHub | UBE2Q1 | 15.0000 | 20.0000 | 0.8670 | 1.438e-30 | 0.0000 | 0.0000 |
| ProlifHub | BUB1B | 16.0000 | 20.0000 | 1.9995 | 1.744e-29 | 0.0000 | 0.0000 |
| ProlifHub | ANLN | 17.0000 | 20.0000 | 2.4953 | 1.246e-31 | 0.0000 | 0.0000 |
| ProlifHub | PTTG1 | 18.0000 | 20.0000 | 1.6487 | 1.238e-28 | 49.2270 | 0.0641 |
| ProlifHub | CENPF | 19.0000 | 20.0000 | 1.8577 | 7.378e-28 | 80.5203 | 0.4026 |
| ProlifHub | IRAK1 | 20.0000 | 20.0000 | 1.0321 | 2.447e-29 | 76.5076 | 0.0543 |
| HepLoss | FCN2 | 1.0000 | 20.0000 | -3.7362 | 3.745e-53 | 0.0000 | 0.0000 |
| HepLoss | CLEC4M | 2.0000 | 20.0000 | -2.9746 | 2.094e-52 | 54.5597 | 0.2055 |
| HepLoss | LINC01093 | 3.0000 | 20.0000 | -4.1296 | 7.692e-48 | 0.0000 | 0.0000 |
| HepLoss | ADAMTS13 | 4.0000 | 20.0000 | -1.6848 | 1.424e-49 | 83.5140 | 0.4225 |
| HepLoss | FAM65C | 5.0000 | 20.0000 | -2.6018 | 1.885e-41 | 74.7681 | 0.4577 |
| HepLoss | ANGPTL6 | 6.0000 | 20.0000 | -1.6108 | 3.358e-63 | 0.0000 | 0.0000 |
| HepLoss | ADGRG7 | 7.0000 | 20.0000 | -2.2502 | 7.762e-41 | 0.0000 | 0.0000 |
| HepLoss | CDHR2 | 8.0000 | 20.0000 | -2.5184 | 1.212e-55 | 56.7001 | 0.2025 |
| HepLoss | CLEC1B | 9.0000 | 20.0000 | -3.2270 | 3.365e-60 | 66.5915 | 0.5529 |
| HepLoss | CXCL14 | 10.0000 | 20.0000 | -4.0699 | 6.344e-69 | 61.0008 | 0.6958 |
| HepLoss | VIPR1 | 11.0000 | 20.0000 | -1.3827 | 2.602e-59 | 93.1888 | 2.7543 |
| HepLoss | PLAC8 | 12.0000 | 20.0000 | -2.1198 | 9.314e-37 | 0.0000 | 0.0000 |
| HepLoss | CFP | 13.0000 | 20.0000 | -1.7341 | 6.087e-35 | 90.2669 | 1.4579 |
| HepLoss | CYP1A2 | 14.0000 | 20.0000 | -3.2742 | 8.287e-35 | 0.0000 | 0.0000 |
| HepLoss | CRHBP | 15.0000 | 20.0000 | -3.1194 | 7.455e-45 | 71.2869 | 0.7551 |
| HepLoss | HHIP | 16.0000 | 20.0000 | -2.8882 | 3.441e-55 | 69.7216 | 0.6128 |
| HepLoss | STAB2 | 17.0000 | 20.0000 | -1.7754 | 3.558e-45 | 82.4416 | 0.6176 |
| HepLoss | FAHD2A | 18.0000 | 20.0000 | -0.9968 | 6.492e-33 | 0.0000 | 0.0000 |
| HepLoss | ECM1 | 19.0000 | 20.0000 | -1.7042 | 5.237e-53 | 25.7087 | 0.0252 |
| HepLoss | FCN3 | 20.0000 | 20.0000 | -3.3961 | 1.547e-41 | 0.0000 | 0.0000 |

**Supplementary Table S5. Multi-cohort validation statistics for module scores.** Validation statistics for ProlifHubScore, HepLossScore, and HCCStateScore across independent GEO HCC cohorts with recoverable tumor/non-tumor labels. The table reports dataset, score, sample counts, tumor and non-tumor means, tumor-minus-non-tumor delta, Cohen's d, nominal p-value, FDR-adjusted p-value, and AUC where applicable. These results support the multi-cohort validation figure in the main text.

| dataset | score | n tumor | n non tumor | delta tumor minus nontumor | cohens d | p value | AUC | AUC low | AUC high | FDR |
| --- | --- | --- | --- | --- | --- | --- | --- | --- | --- | --- |
| GSE14520 | ProlifHubScore | 225.0000 | 220.0000 | 1.4426 | 3.9887 | 3.640e-138 | 0.9887 | 0.9805 | 0.9970 | 1.092e-137 |
| GSE14520 | HepLossScore | 225.0000 | 220.0000 | -1.5649 | -4.5027 | 2.928e-169 | 0.0126 | 0.0019 | 0.0234 | 2.050e-168 |
| GSE14520 | HCCStateScore | 225.0000 | 220.0000 | 3.0075 | 4.9309 | 1.056e-187 | 0.9906 | 0.9825 | 0.9988 | 1.109e-186 |
| GSE25097 | ProlifHubScore | 268.0000 | 289.0000 | 1.2314 | 3.0452 | 3.327e-110 | 0.9934 | 0.9886 | 0.9981 | 7.762e-110 |
| GSE25097 | HepLossScore | 268.0000 | 289.0000 | -1.4773 | -4.0254 | 3.982e-164 | 0.0037 | 2.188e-04 | 0.0071 | 1.672e-163 |
| GSE25097 | HCCStateScore | 268.0000 | 289.0000 | 2.7087 | 4.4412 | 4.436e-212 | 0.9980 | 0.9959 | 1.0000 | 9.316e-211 |
| GSE76427 | ProlifHubScore | 115.0000 | 52.0000 | 0.9280 | 1.9298 | 2.613e-33 | 0.9423 | 0.9008 | 0.9838 | 4.988e-33 |
| GSE76427 | HepLossScore | 115.0000 | 52.0000 | -1.5468 | -4.0950 | 5.194e-31 | 0.0097 | 0.0000 | 0.0195 | 9.089e-31 |
| GSE76427 | HCCStateScore | 115.0000 | 52.0000 | 2.4749 | 3.4063 | 9.317e-42 | 0.9758 | 0.9523 | 0.9992 | 1.957e-41 |
| GSE36376 | ProlifHubScore | 240.0000 | 193.0000 | 1.3061 | 3.8026 | 3.443e-132 | 0.9974 | 0.9948 | 1.0000 | 9.039e-132 |
| GSE36376 | HepLossScore | 240.0000 | 193.0000 | -1.2658 | -4.1056 | 2.083e-160 | 0.0171 | 0.0046 | 0.0296 | 7.291e-160 |
| GSE36376 | HCCStateScore | 240.0000 | 193.0000 | 2.5718 | 4.5942 | 2.431e-168 | 0.9934 | 0.9852 | 1.0000 | 1.276e-167 |
| GSE57957 | ProlifHubScore | 39.0000 | 39.0000 | 0.9911 | 1.9207 | 7.605e-12 | 0.9421 | 0.8764 | 1.0000 | 7.605e-12 |
| GSE57957 | HepLossScore | 39.0000 | 39.0000 | -1.3802 | -3.0014 | 3.543e-21 | 0.0592 | 0.0000 | 0.1241 | 4.133e-21 |
| GSE57957 | HCCStateScore | 39.0000 | 39.0000 | 2.3714 | 2.8006 | 1.442e-19 | 0.9402 | 0.8726 | 1.0000 | 1.594e-19 |
| GSE45267 | ProlifHubScore | 46.0000 | 41.0000 | 1.5341 | 3.8266 | 9.575e-28 | 0.9666 | 0.9256 | 1.0000 | 1.436e-27 |
| GSE45267 | HepLossScore | 46.0000 | 41.0000 | -1.5707 | -3.9329 | 4.217e-22 | 0.0456 | 0.0000 | 0.1013 | 5.209e-22 |
| GSE45267 | HCCStateScore | 46.0000 | 41.0000 | 3.1048 | 4.1175 | 2.116e-25 | 0.9613 | 0.9141 | 1.0000 | 2.963e-25 |
| GSE112790 | ProlifHubScore | 183.0000 | 15.0000 | 1.9642 | 3.4822 | 3.597e-28 | 0.9960 | 0.9891 | 1.0000 | 5.811e-28 |
| GSE112790 | HepLossScore | 183.0000 | 15.0000 | -2.1264 | -4.4086 | 3.144e-16 | 0.0080 | 0.0000 | 0.0184 | 3.301e-16 |
| GSE112790 | HCCStateScore | 183.0000 | 15.0000 | 4.0907 | 4.2572 | 1.387e-23 | 0.9960 | 0.9896 | 1.0000 | 1.821e-23 |

**Supplementary Table S6. Module-size robustness statistics.** Sensitivity analysis evaluating HCCStateScore performance across alternative module definitions using the top 10, 15, 20, 30, and 50 conserved genes per module. The table reports dataset, module size, score, tumor-versus-non-tumor delta, AUC, p-value, and effect size. These data support the module-size robustness supplementary figure and demonstrate that the score is not dependent on the top-20 cutoff.

| dataset | score | n tumor | n non tumor | delta tumor minus nontumor | cohens d | p value | AUC | AUC low | AUC high | module size |
| --- | --- | --- | --- | --- | --- | --- | --- | --- | --- | --- |
| GSE121248 | HCCStateScore | 70.0000 | 37.0000 | 3.2692 | 3.7849 | 3.484e-32 | 0.9614 | 0.9081 | 1.0000 | 10.0000 |
| GSE41804 | HCCStateScore | 20.0000 | 20.0000 | 2.8264 | 2.8006 | 1.540e-10 | 0.9650 | 0.9080 | 1.0000 | 10.0000 |
| GSE14520 | HCCStateScore | 225.0000 | 220.0000 | 3.0222 | 4.7477 | 1.259e-183 | 0.9910 | 0.9819 | 1.0000 | 10.0000 |
| GSE25097 | HCCStateScore | 268.0000 | 289.0000 | 2.6331 | 4.2757 | 6.858e-206 | 0.9979 | 0.9961 | 0.9997 | 10.0000 |
| GSE76427 | HCCStateScore | 115.0000 | 52.0000 | 2.5062 | 3.2657 | 8.536e-38 | 0.9689 | 0.9391 | 0.9987 | 10.0000 |
| GSE36376 | HCCStateScore | 240.0000 | 193.0000 | 2.5503 | 4.6419 | 7.168e-173 | 0.9938 | 0.9860 | 1.0000 | 10.0000 |
| GSE57957 | HCCStateScore | 39.0000 | 39.0000 | 2.5296 | 2.9342 | 1.770e-20 | 0.9421 | 0.8766 | 1.0000 | 10.0000 |
| GSE45267 | HCCStateScore | 46.0000 | 41.0000 | 3.1140 | 4.2292 | 7.191e-26 | 0.9671 | 0.9245 | 1.0000 | 10.0000 |
| GSE112790 | HCCStateScore | 183.0000 | 15.0000 | 4.3278 | 4.5703 | 9.025e-24 | 0.9978 | 0.9940 | 1.0000 | 10.0000 |
| GSE121248 | HCCStateScore | 70.0000 | 37.0000 | 3.2209 | 3.7985 | 1.456e-32 | 0.9614 | 0.9078 | 1.0000 | 15.0000 |
| GSE41804 | HCCStateScore | 20.0000 | 20.0000 | 2.7987 | 2.8408 | 1.144e-10 | 0.9675 | 0.9150 | 1.0000 | 15.0000 |
| GSE14520 | HCCStateScore | 225.0000 | 220.0000 | 2.9340 | 4.8925 | 1.015e-188 | 0.9903 | 0.9818 | 0.9989 | 15.0000 |
| GSE25097 | HCCStateScore | 268.0000 | 289.0000 | 2.6527 | 4.5014 | 3.215e-218 | 0.9978 | 0.9958 | 0.9999 | 15.0000 |
| GSE76427 | HCCStateScore | 115.0000 | 52.0000 | 2.3967 | 3.4141 | 5.588e-39 | 0.9706 | 0.9409 | 1.0000 | 15.0000 |
| GSE36376 | HCCStateScore | 240.0000 | 193.0000 | 2.5593 | 4.6561 | 4.384e-178 | 0.9919 | 0.9831 | 1.0000 | 15.0000 |
| GSE57957 | HCCStateScore | 39.0000 | 39.0000 | 2.3273 | 2.8557 | 3.790e-20 | 0.9435 | 0.8789 | 1.0000 | 15.0000 |
| GSE45267 | HCCStateScore | 46.0000 | 41.0000 | 3.0883 | 4.1962 | 4.654e-25 | 0.9618 | 0.9148 | 1.0000 | 15.0000 |
| GSE112790 | HCCStateScore | 183.0000 | 15.0000 | 4.1569 | 4.5136 | 2.111e-22 | 0.9964 | 0.9906 | 1.0000 | 15.0000 |
| GSE121248 | HCCStateScore | 70.0000 | 37.0000 | 3.1934 | 3.6688 | 1.612e-31 | 0.9614 | 0.9073 | 1.0000 | 20.0000 |
| GSE41804 | HCCStateScore | 20.0000 | 20.0000 | 2.7829 | 2.7492 | 2.572e-10 | 0.9625 | 0.9010 | 1.0000 | 20.0000 |
| GSE14520 | HCCStateScore | 225.0000 | 220.0000 | 3.0075 | 4.9309 | 1.056e-187 | 0.9906 | 0.9825 | 0.9988 | 20.0000 |
| GSE25097 | HCCStateScore | 268.0000 | 289.0000 | 2.7087 | 4.4412 | 4.436e-212 | 0.9980 | 0.9959 | 1.0000 | 20.0000 |
| GSE76427 | HCCStateScore | 115.0000 | 52.0000 | 2.4749 | 3.4063 | 9.317e-42 | 0.9758 | 0.9523 | 0.9992 | 20.0000 |
| GSE36376 | HCCStateScore | 240.0000 | 193.0000 | 2.5718 | 4.5942 | 2.431e-168 | 0.9934 | 0.9852 | 1.0000 | 20.0000 |
| GSE57957 | HCCStateScore | 39.0000 | 39.0000 | 2.3714 | 2.8006 | 1.442e-19 | 0.9402 | 0.8726 | 1.0000 | 20.0000 |
| GSE45267 | HCCStateScore | 46.0000 | 41.0000 | 3.1048 | 4.1175 | 2.116e-25 | 0.9613 | 0.9141 | 1.0000 | 20.0000 |
| GSE112790 | HCCStateScore | 183.0000 | 15.0000 | 4.0907 | 4.2572 | 1.387e-23 | 0.9960 | 0.9896 | 1.0000 | 20.0000 |
| GSE121248 | HCCStateScore | 70.0000 | 37.0000 | 3.1407 | 3.6024 | 1.132e-31 | 0.9622 | 0.9085 | 1.0000 | 30.0000 |
| GSE41804 | HCCStateScore | 20.0000 | 20.0000 | 2.7367 | 2.7735 | 2.560e-10 | 0.9700 | 0.9183 | 1.0000 | 30.0000 |
| GSE14520 | HCCStateScore | 225.0000 | 220.0000 | 2.9167 | 4.8920 | 1.317e-177 | 0.9904 | 0.9822 | 0.9986 | 30.0000 |
| GSE25097 | HCCStateScore | 268.0000 | 289.0000 | 2.6097 | 4.4116 | 1.162e-202 | 0.9977 | 0.9951 | 1.0000 | 30.0000 |
| GSE76427 | HCCStateScore | 115.0000 | 52.0000 | 2.3970 | 3.4445 | 2.312e-45 | 0.9789 | 0.9578 | 1.0000 | 30.0000 |
| GSE36376 | HCCStateScore | 240.0000 | 193.0000 | 2.5118 | 4.6788 | 4.773e-167 | 0.9939 | 0.9858 | 1.0000 | 30.0000 |
| GSE57957 | HCCStateScore | 39.0000 | 39.0000 | 2.2526 | 2.8757 | 8.346e-20 | 0.9415 | 0.8751 | 1.0000 | 30.0000 |
| GSE45267 | HCCStateScore | 46.0000 | 41.0000 | 3.0389 | 4.0051 | 3.172e-26 | 0.9613 | 0.9130 | 1.0000 | 30.0000 |
| GSE112790 | HCCStateScore | 183.0000 | 15.0000 | 3.9446 | 4.1088 | 5.161e-29 | 0.9942 | 0.9840 | 1.0000 | 30.0000 |
| GSE121248 | HCCStateScore | 70.0000 | 37.0000 | 3.0599 | 3.5770 | 2.943e-32 | 0.9641 | 0.9133 | 1.0000 | 50.0000 |
| GSE41804 | HCCStateScore | 20.0000 | 20.0000 | 2.6827 | 2.8307 | 1.772e-10 | 0.9700 | 0.9219 | 1.0000 | 50.0000 |
| GSE14520 | HCCStateScore | 225.0000 | 220.0000 | 2.8609 | 4.9268 | 6.803e-177 | 0.9903 | 0.9822 | 0.9984 | 50.0000 |
| GSE25097 | HCCStateScore | 268.0000 | 289.0000 | 2.5552 | 4.2816 | 3.456e-197 | 0.9970 | 0.9938 | 1.0000 | 50.0000 |
| GSE76427 | HCCStateScore | 115.0000 | 52.0000 | 2.3248 | 3.4684 | 9.141e-46 | 0.9791 | 0.9590 | 0.9992 | 50.0000 |
| GSE36376 | HCCStateScore | 240.0000 | 193.0000 | 2.3808 | 4.7233 | 3.493e-165 | 0.9946 | 0.9875 | 1.0000 | 50.0000 |
| GSE57957 | HCCStateScore | 39.0000 | 39.0000 | 2.1550 | 2.9999 | 1.328e-20 | 0.9454 | 0.8821 | 1.0000 | 50.0000 |
| GSE45267 | HCCStateScore | 46.0000 | 41.0000 | 2.9409 | 3.8907 | 2.782e-26 | 0.9581 | 0.9063 | 1.0000 | 50.0000 |
| GSE112790 | HCCStateScore | 183.0000 | 15.0000 | 3.7442 | 3.9632 | 1.391e-34 | 0.9942 | 0.9840 | 1.0000 | 50.0000 |

**Supplementary Table S7. HBV_INJURY derivation from GSE83148.** Gene-level derivation statistics for the HBV injury axis constructed from the GSE83148 ordinal ALT/AST/HBV-DNA injury index. For each gene, the table reports the coefficient for association with the continuous HBV_INJURY_INDEX, moderated statistic, p-value, FDR, rank, and membership in the top-200, top-500, top-1000, top-2000, top-5000, and extended FDR-defined injury programs.

| gene | beta injury index | AveExpr | t | p value | FDR | B | logFC | P Value | adj P Val | hbv injury rank | included TOP 200 | included TOP 500 | included TOP 1000 | included TOP 2000 | included TOP 5000 | included EXTENDED 7792 |
| --- | --- | --- | --- | --- | --- | --- | --- | --- | --- | --- | --- | --- | --- | --- | --- | --- |
| PTAFR | 0.3857 | 6.7152 | 9.6019 | 7.564e-17 | 1.419e-12 | 27.7098 | 0.3857 | 7.564e-17 | 1.419e-12 | 1.0000 | TRUE | TRUE | TRUE | TRUE | TRUE | TRUE |
| UBD | 1.4037 | 9.0074 | 9.5060 | 1.305e-16 | 1.419e-12 | 27.1831 | 1.4037 | 1.305e-16 | 1.419e-12 | 2.0000 | TRUE | TRUE | TRUE | TRUE | TRUE | TRUE |
| NUB1 | 0.2294 | 7.9835 | 9.3420 | 3.308e-16 | 2.399e-12 | 26.2840 | 0.2294 | 3.308e-16 | 2.399e-12 | 3.0000 | TRUE | TRUE | TRUE | TRUE | TRUE | TRUE |
| PNMA2 | 0.6180 | 4.8432 | 9.2662 | 5.081e-16 | 2.763e-12 | 25.8694 | 0.6180 | 5.081e-16 | 2.763e-12 | 4.0000 | TRUE | TRUE | TRUE | TRUE | TRUE | TRUE |
| LPXN | 0.3500 | 7.0703 | 9.1599 | 9.265e-16 | 4.031e-12 | 25.2891 | 0.3500 | 9.265e-16 | 4.031e-12 | 5.0000 | TRUE | TRUE | TRUE | TRUE | TRUE | TRUE |
| RTN1 | 0.4576 | 4.9656 | 8.8878 | 4.281e-15 | 1.552e-11 | 23.8100 | 0.4576 | 4.281e-15 | 1.552e-11 | 6.0000 | TRUE | TRUE | TRUE | TRUE | TRUE | TRUE |
| CDC25B | 0.3367 | 7.0787 | 8.7148 | 1.127e-14 | 3.503e-11 | 22.8746 | 0.3367 | 1.127e-14 | 3.503e-11 | 7.0000 | TRUE | TRUE | TRUE | TRUE | TRUE | TRUE |
| HCP5 | 0.5629 | 6.3983 | 8.6513 | 1.607e-14 | 3.929e-11 | 22.5320 | 0.5629 | 1.607e-14 | 3.929e-11 | 8.0000 | TRUE | TRUE | TRUE | TRUE | TRUE | TRUE |
| LOC101928589 | 0.3665 | 5.7738 | 8.6492 | 1.625e-14 | 3.929e-11 | 22.5209 | 0.3665 | 1.625e-14 | 3.929e-11 | 9.0000 | TRUE | TRUE | TRUE | TRUE | TRUE | TRUE |
| FGD2 | 0.3260 | 8.9556 | 8.5749 | 2.458e-14 | 4.556e-11 | 22.1213 | 0.3260 | 2.458e-14 | 4.556e-11 | 10.0000 | TRUE | TRUE | TRUE | TRUE | TRUE | TRUE |
| MAML2 | 0.4585 | 5.7168 | 8.5683 | 2.550e-14 | 4.556e-11 | 22.0858 | 0.4585 | 2.550e-14 | 4.556e-11 | 11.0000 | TRUE | TRUE | TRUE | TRUE | TRUE | TRUE |
| CXCL10 | 1.1889 | 8.5253 | 8.5597 | 2.675e-14 | 4.556e-11 | 22.0396 | 1.1889 | 2.675e-14 | 4.556e-11 | 12.0000 | TRUE | TRUE | TRUE | TRUE | TRUE | TRUE |
| KLHL6 | 0.5413 | 4.9232 | 8.5565 | 2.723e-14 | 4.556e-11 | 22.0225 | 0.5413 | 2.723e-14 | 4.556e-11 | 13.0000 | TRUE | TRUE | TRUE | TRUE | TRUE | TRUE |
| EGR2 | 0.5715 | 3.8836 | 8.5387 | 3.005e-14 | 4.670e-11 | 21.9272 | 0.5715 | 3.005e-14 | 4.670e-11 | 14.0000 | TRUE | TRUE | TRUE | TRUE | TRUE | TRUE |
| BARD1 | 0.4697 | 4.6483 | 8.5003 | 3.720e-14 | 5.123e-11 | 21.7211 | 0.4697 | 3.720e-14 | 5.123e-11 | 15.0000 | TRUE | TRUE | TRUE | TRUE | TRUE | TRUE |
| BTG2 | 0.4709 | 6.4262 | 8.4980 | 3.768e-14 | 5.123e-11 | 21.7086 | 0.4709 | 3.768e-14 | 5.123e-11 | 16.0000 | TRUE | TRUE | TRUE | TRUE | TRUE | TRUE |
| MTHFD2 | 0.7030 | 5.6313 | 8.4856 | 4.036e-14 | 5.165e-11 | 21.6422 | 0.7030 | 4.036e-14 | 5.165e-11 | 17.0000 | TRUE | TRUE | TRUE | TRUE | TRUE | TRUE |
| SAMD9L | 0.5428 | 6.5597 | 8.4349 | 5.346e-14 | 6.460e-11 | 21.3708 | 0.5428 | 5.346e-14 | 6.460e-11 | 18.0000 | TRUE | TRUE | TRUE | TRUE | TRUE | TRUE |
| STAT1 | 0.5231 | 10.2177 | 8.4114 | 6.086e-14 | 6.968e-11 | 21.2455 | 0.5231 | 6.086e-14 | 6.968e-11 | 19.0000 | TRUE | TRUE | TRUE | TRUE | TRUE | TRUE |
| CDCP1 | 0.3016 | 4.8835 | 8.3480 | 8.641e-14 | 8.789e-11 | 20.9069 | 0.3016 | 8.641e-14 | 8.789e-11 | 20.0000 | TRUE | TRUE | TRUE | TRUE | TRUE | TRUE |
| HPSE | 0.6149 | 5.7839 | 8.3440 | 8.835e-14 | 8.789e-11 | 20.8854 | 0.6149 | 8.835e-14 | 8.789e-11 | 21.0000 | TRUE | TRUE | TRUE | TRUE | TRUE | TRUE |
| MAB21L2 | 0.6191 | 5.0647 | 8.3358 | 9.246e-14 | 8.789e-11 | 20.8415 | 0.6191 | 9.246e-14 | 8.789e-11 | 22.0000 | TRUE | TRUE | TRUE | TRUE | TRUE | TRUE |
| DOCK11 | 0.4955 | 6.1949 | 8.3348 | 9.293e-14 | 8.789e-11 | 20.8366 | 0.4955 | 9.293e-14 | 8.789e-11 | 23.0000 | TRUE | TRUE | TRUE | TRUE | TRUE | TRUE |
| CDCA7 | 0.4591 | 4.2595 | 8.3220 | 9.975e-14 | 8.879e-11 | 20.7682 | 0.4591 | 9.975e-14 | 8.879e-11 | 24.0000 | TRUE | TRUE | TRUE | TRUE | TRUE | TRUE |
| JUN | 0.5181 | 7.8988 | 8.3140 | 1.042e-13 | 8.879e-11 | 20.7257 | 0.5181 | 1.042e-13 | 8.879e-11 | 25.0000 | TRUE | TRUE | TRUE | TRUE | TRUE | TRUE |
| TLR7 | 0.3934 | 4.1078 | 8.3086 | 1.074e-13 | 8.879e-11 | 20.6968 | 0.3934 | 1.074e-13 | 8.879e-11 | 26.0000 | TRUE | TRUE | TRUE | TRUE | TRUE | TRUE |
| C5AR1 | 0.4537 | 5.3485 | 8.3039 | 1.102e-13 | 8.879e-11 | 20.6718 | 0.4537 | 1.102e-13 | 8.879e-11 | 27.0000 | TRUE | TRUE | TRUE | TRUE | TRUE | TRUE |
| MX2 | 0.4970 | 6.3262 | 8.2855 | 1.220e-13 | 9.477e-11 | 20.5738 | 0.4970 | 1.220e-13 | 9.477e-11 | 28.0000 | TRUE | TRUE | TRUE | TRUE | TRUE | TRUE |
| TCIRG1 | 0.2992 | 7.8381 | 8.2541 | 1.450e-13 | 1.087e-10 | 20.4070 | 0.2992 | 1.450e-13 | 1.087e-10 | 29.0000 | TRUE | TRUE | TRUE | TRUE | TRUE | TRUE |
| FPR3 | 0.5291 | 6.9881 | 8.2444 | 1.530e-13 | 1.109e-10 | 20.3551 | 0.5291 | 1.530e-13 | 1.109e-10 | 30.0000 | TRUE | TRUE | TRUE | TRUE | TRUE | TRUE |
| STX7 | 0.3103 | 7.0058 | 8.2378 | 1.586e-13 | 1.113e-10 | 20.3200 | 0.3103 | 1.586e-13 | 1.113e-10 | 31.0000 | TRUE | TRUE | TRUE | TRUE | TRUE | TRUE |
| PTPRE | 0.4440 | 6.0336 | 8.2298 | 1.657e-13 | 1.126e-10 | 20.2779 | 0.4440 | 1.657e-13 | 1.126e-10 | 32.0000 | TRUE | TRUE | TRUE | TRUE | TRUE | TRUE |
| EMILIN2 | 0.4358 | 4.8761 | 8.2241 | 1.710e-13 | 1.127e-10 | 20.2476 | 0.4358 | 1.710e-13 | 1.127e-10 | 33.0000 | TRUE | TRUE | TRUE | TRUE | TRUE | TRUE |
| CD300LF | 0.5054 | 5.3811 | 8.2033 | 1.917e-13 | 1.180e-10 | 20.1372 | 0.5054 | 1.917e-13 | 1.180e-10 | 34.0000 | TRUE | TRUE | TRUE | TRUE | TRUE | TRUE |
| IL32 | 0.7695 | 8.1167 | 8.2003 | 1.949e-13 | 1.180e-10 | 20.1211 | 0.7695 | 1.949e-13 | 1.180e-10 | 35.0000 | TRUE | TRUE | TRUE | TRUE | TRUE | TRUE |
| RGS1 | 0.6268 | 5.4629 | 8.2000 | 1.952e-13 | 1.180e-10 | 20.1195 | 0.6268 | 1.952e-13 | 1.180e-10 | 36.0000 | TRUE | TRUE | TRUE | TRUE | TRUE | TRUE |
| MICB | 0.5437 | 5.8001 | 8.1875 | 2.091e-13 | 1.229e-10 | 20.0531 | 0.5437 | 2.091e-13 | 1.229e-10 | 37.0000 | TRUE | TRUE | TRUE | TRUE | TRUE | TRUE |
| LAIR1 | 0.4475 | 5.8871 | 8.1766 | 2.220e-13 | 1.254e-10 | 19.9956 | 0.4475 | 2.220e-13 | 1.254e-10 | 38.0000 | TRUE | TRUE | TRUE | TRUE | TRUE | TRUE |
| SLC7A6 | 0.4430 | 5.9169 | 8.1742 | 2.249e-13 | 1.254e-10 | 19.9829 | 0.4430 | 2.249e-13 | 1.254e-10 | 39.0000 | TRUE | TRUE | TRUE | TRUE | TRUE | TRUE |
| LAMP3 | 0.6896 | 4.7213 | 8.1524 | 2.534e-13 | 1.378e-10 | 19.8675 | 0.6896 | 2.534e-13 | 1.378e-10 | 40.0000 | TRUE | TRUE | TRUE | TRUE | TRUE | TRUE |
| CD83 | 0.3637 | 5.9102 | 8.1196 | 3.034e-13 | 1.610e-10 | 19.6938 | 0.3637 | 3.034e-13 | 1.610e-10 | 41.0000 | TRUE | TRUE | TRUE | TRUE | TRUE | TRUE |
| FGR | 0.4172 | 6.3404 | 8.0813 | 3.741e-13 | 1.938e-10 | 19.4914 | 0.4172 | 3.741e-13 | 1.938e-10 | 42.0000 | TRUE | TRUE | TRUE | TRUE | TRUE | TRUE |
| STAT4 | 0.3116 | 5.9524 | 8.0683 | 4.016e-13 | 2.032e-10 | 19.4228 | 0.3116 | 4.016e-13 | 2.032e-10 | 43.0000 | TRUE | TRUE | TRUE | TRUE | TRUE | TRUE |
| CFL1 | 0.2502 | 11.6824 | 8.0601 | 4.202e-13 | 2.057e-10 | 19.3792 | 0.2502 | 4.202e-13 | 2.057e-10 | 44.0000 | TRUE | TRUE | TRUE | TRUE | TRUE | TRUE |
| HOXB2 | 0.2527 | 6.8311 | 8.0559 | 4.299e-13 | 2.057e-10 | 19.3571 | 0.2527 | 4.299e-13 | 2.057e-10 | 45.0000 | TRUE | TRUE | TRUE | TRUE | TRUE | TRUE |
| IFIT5 | 0.3215 | 7.3785 | 8.0538 | 4.349e-13 | 2.057e-10 | 19.3460 | 0.3215 | 4.349e-13 | 2.057e-10 | 46.0000 | TRUE | TRUE | TRUE | TRUE | TRUE | TRUE |
| FABP5 | 0.8749 | 6.5896 | 8.0470 | 4.512e-13 | 2.088e-10 | 19.3104 | 0.8749 | 4.512e-13 | 2.088e-10 | 47.0000 | TRUE | TRUE | TRUE | TRUE | TRUE | TRUE |
| CCND2 | 0.5895 | 6.0813 | 8.0392 | 4.709e-13 | 2.134e-10 | 19.2691 | 0.5895 | 4.709e-13 | 2.134e-10 | 48.0000 | TRUE | TRUE | TRUE | TRUE | TRUE | TRUE |
| NUSAP1 | 0.7015 | 6.2467 | 8.0342 | 4.841e-13 | 2.149e-10 | 19.2425 | 0.7015 | 4.841e-13 | 2.149e-10 | 49.0000 | TRUE | TRUE | TRUE | TRUE | TRUE | TRUE |
| SLC6A6 | 0.5206 | 5.9696 | 8.0004 | 5.820e-13 | 2.532e-10 | 19.0646 | 0.5206 | 5.820e-13 | 2.532e-10 | 50.0000 | TRUE | TRUE | TRUE | TRUE | TRUE | TRUE |
| GRK3 | 0.4810 | 5.9689 | 7.9938 | 6.034e-13 | 2.573e-10 | 19.0298 | 0.4810 | 6.034e-13 | 2.573e-10 | 51.0000 | TRUE | TRUE | TRUE | TRUE | TRUE | TRUE |
| CXCL9 | 1.1416 | 8.5611 | 7.9891 | 6.190e-13 | 2.589e-10 | 19.0052 | 1.1416 | 6.190e-13 | 2.589e-10 | 52.0000 | TRUE | TRUE | TRUE | TRUE | TRUE | TRUE |
| TC2N | 0.4004 | 3.1738 | 7.9743 | 6.708e-13 | 2.753e-10 | 18.9276 | 0.4004 | 6.708e-13 | 2.753e-10 | 53.0000 | TRUE | TRUE | TRUE | TRUE | TRUE | TRUE |
| SLC7A1 | 0.5914 | 5.6576 | 7.9562 | 7.402e-13 | 2.938e-10 | 18.8324 | 0.5914 | 7.402e-13 | 2.938e-10 | 54.0000 | TRUE | TRUE | TRUE | TRUE | TRUE | TRUE |
| SH2D2A | 0.2329 | 5.4061 | 7.9556 | 7.430e-13 | 2.938e-10 | 18.8288 | 0.2329 | 7.430e-13 | 2.938e-10 | 55.0000 | TRUE | TRUE | TRUE | TRUE | TRUE | TRUE |
| CCNA2 | 0.5547 | 4.8342 | 7.9161 | 9.206e-13 | 3.521e-10 | 18.6219 | 0.5547 | 9.206e-13 | 3.521e-10 | 56.0000 | TRUE | TRUE | TRUE | TRUE | TRUE | TRUE |
| ZNF738 | 0.4880 | 4.6982 | 7.9149 | 9.269e-13 | 3.521e-10 | 18.6152 | 0.4880 | 9.269e-13 | 3.521e-10 | 57.0000 | TRUE | TRUE | TRUE | TRUE | TRUE | TRUE |
| PCED1B | 0.3642 | 5.6047 | 7.9125 | 9.389e-13 | 3.521e-10 | 18.6028 | 0.3642 | 9.389e-13 | 3.521e-10 | 58.0000 | TRUE | TRUE | TRUE | TRUE | TRUE | TRUE |
| LOC101928269 | 0.3371 | 6.0468 | 7.8887 | 1.068e-12 | 3.939e-10 | 18.4782 | 0.3371 | 1.068e-12 | 3.939e-10 | 59.0000 | TRUE | TRUE | TRUE | TRUE | TRUE | TRUE |
| PARVG | 0.3529 | 6.4271 | 7.8630 | 1.228e-12 | 4.454e-10 | 18.3433 | 0.3529 | 1.228e-12 | 4.454e-10 | 60.0000 | TRUE | TRUE | TRUE | TRUE | TRUE | TRUE |
| SLAMF7 | 0.7459 | 5.8001 | 7.8587 | 1.257e-12 | 4.483e-10 | 18.3209 | 0.7459 | 1.257e-12 | 4.483e-10 | 61.0000 | TRUE | TRUE | TRUE | TRUE | TRUE | TRUE |
| GAS7 | 0.2858 | 6.0658 | 7.8492 | 1.324e-12 | 4.645e-10 | 18.2711 | 0.2858 | 1.324e-12 | 4.645e-10 | 62.0000 | TRUE | TRUE | TRUE | TRUE | TRUE | TRUE |
| OSBPL3 | 0.5127 | 4.5982 | 7.8413 | 1.382e-12 | 4.771e-10 | 18.2298 | 0.5127 | 1.382e-12 | 4.771e-10 | 63.0000 | TRUE | TRUE | TRUE | TRUE | TRUE | TRUE |
| ARHGAP18 | 0.3798 | 6.2659 | 7.8372 | 1.413e-12 | 4.801e-10 | 18.2085 | 0.3798 | 1.413e-12 | 4.801e-10 | 64.0000 | TRUE | TRUE | TRUE | TRUE | TRUE | TRUE |
| PLP2 | 0.4593 | 6.4778 | 7.8304 | 1.466e-12 | 4.905e-10 | 18.1729 | 0.4593 | 1.466e-12 | 4.905e-10 | 65.0000 | TRUE | TRUE | TRUE | TRUE | TRUE | TRUE |
| RAP2B | 0.3899 | 8.1141 | 7.8258 | 1.502e-12 | 4.905e-10 | 18.1491 | 0.3899 | 1.502e-12 | 4.905e-10 | 66.0000 | TRUE | TRUE | TRUE | TRUE | TRUE | TRUE |
| ENTPD1 | 0.3886 | 6.5052 | 7.8248 | 1.511e-12 | 4.905e-10 | 18.1435 | 0.3886 | 1.511e-12 | 4.905e-10 | 67.0000 | TRUE | TRUE | TRUE | TRUE | TRUE | TRUE |
| PLXNC1 | 0.7047 | 5.1847 | 7.8134 | 1.606e-12 | 5.139e-10 | 18.0843 | 0.7047 | 1.606e-12 | 5.139e-10 | 68.0000 | TRUE | TRUE | TRUE | TRUE | TRUE | TRUE |
| HLA-B | 0.2842 | 12.6414 | 7.8035 | 1.695e-12 | 5.343e-10 | 18.0327 | 0.2842 | 1.695e-12 | 5.343e-10 | 69.0000 | TRUE | TRUE | TRUE | TRUE | TRUE | TRUE |
| MYO5A | 0.4751 | 5.9812 | 7.7726 | 2.003e-12 | 6.216e-10 | 17.8712 | 0.4751 | 2.003e-12 | 6.216e-10 | 70.0000 | TRUE | TRUE | TRUE | TRUE | TRUE | TRUE |
| CRLF3 | 0.2994 | 7.7927 | 7.7702 | 2.029e-12 | 6.216e-10 | 17.8589 | 0.2994 | 2.029e-12 | 6.216e-10 | 71.0000 | TRUE | TRUE | TRUE | TRUE | TRUE | TRUE |
| STK17B | 0.5795 | 6.5563 | 7.7637 | 2.102e-12 | 6.351e-10 | 17.8247 | 0.5795 | 2.102e-12 | 6.351e-10 | 72.0000 | TRUE | TRUE | TRUE | TRUE | TRUE | TRUE |
| CD27 | 0.5203 | 6.0183 | 7.7553 | 2.199e-12 | 6.450e-10 | 17.7811 | 0.5203 | 2.199e-12 | 6.450e-10 | 73.0000 | TRUE | TRUE | TRUE | TRUE | TRUE | TRUE |
| PAG1 | 0.4445 | 7.4135 | 7.7547 | 2.207e-12 | 6.450e-10 | 17.7779 | 0.4445 | 2.207e-12 | 6.450e-10 | 74.0000 | TRUE | TRUE | TRUE | TRUE | TRUE | TRUE |
| BCAT1 | 0.6529 | 4.4456 | 7.7533 | 2.224e-12 | 6.450e-10 | 17.7704 | 0.6529 | 2.224e-12 | 6.450e-10 | 75.0000 | TRUE | TRUE | TRUE | TRUE | TRUE | TRUE |
| SLC25A24 | 0.4507 | 5.9911 | 7.7471 | 2.299e-12 | 6.515e-10 | 17.7382 | 0.4507 | 2.299e-12 | 6.515e-10 | 76.0000 | TRUE | TRUE | TRUE | TRUE | TRUE | TRUE |
| RAD51AP1 | 0.5718 | 4.5873 | 7.7465 | 2.306e-12 | 6.515e-10 | 17.7353 | 0.5718 | 2.306e-12 | 6.515e-10 | 77.0000 | TRUE | TRUE | TRUE | TRUE | TRUE | TRUE |
| MCUB | 0.4066 | 5.6080 | 7.7432 | 2.348e-12 | 6.547e-10 | 17.7181 | 0.4066 | 2.348e-12 | 6.547e-10 | 78.0000 | TRUE | TRUE | TRUE | TRUE | TRUE | TRUE |
| C15ORF48 | 0.8272 | 4.2946 | 7.7372 | 2.425e-12 | 6.677e-10 | 17.6869 | 0.8272 | 2.425e-12 | 6.677e-10 | 79.0000 | TRUE | TRUE | TRUE | TRUE | TRUE | TRUE |
| BTN2A2 | 0.2619 | 6.2204 | 7.7227 | 2.621e-12 | 7.060e-10 | 17.6116 | 0.2619 | 2.621e-12 | 7.060e-10 | 80.0000 | TRUE | TRUE | TRUE | TRUE | TRUE | TRUE |

**Supplementary Table S8. HBV_INJURY top-N and CIBERSORTx-adjusted regression sensitivity.** Regression results for ranked HBV injury programs projected into GSE121248. The table reports injury-set definition, number of input genes, number of genes overlapping the expression matrix, regression model, tumor coefficient, 95% confidence interval, p-value, and percentage of effect retained relative to the unadjusted model. Models include unadjusted, E2F/G2M-adjusted, E2F/G2M plus CIBERSORTx principal-component-adjusted, and E2F/G2M plus selected-fraction-adjusted analyses.

| analysis role | injury set | n input genes | n overlap genes | model | tumor coefficient | percent retained vs unadjusted | p value | ci low | ci high |
| --- | --- | --- | --- | --- | --- | --- | --- | --- | --- |
| topN_sensitivity | HBV_INJURY_TOP_200 | 200.0000 | 200.0000 | unadjusted | 0.1337 | 100.0000 | 0.2081 | -0.0756 | 0.3431 |
| topN_sensitivity | HBV_INJURY_TOP_200 | 200.0000 | 200.0000 | proliferation_adjusted | 0.1706 | 127.5511 | 0.2142 | -0.1001 | 0.4413 |
| topN_sensitivity | HBV_INJURY_TOP_200 | 200.0000 | 200.0000 | proliferation_cibersortx_pc_adjusted | 0.2687 | 200.8945 | 0.0300 | 0.0265 | 0.5108 |
| topN_sensitivity | HBV_INJURY_TOP_200 | 200.0000 | 200.0000 | proliferation_selected_fraction_adjusted | 0.2383 | 178.1999 | 0.0404 | 0.0107 | 0.4659 |
| topN_sensitivity | HBV_INJURY_TOP_500 | 500.0000 | 500.0000 | unadjusted | 0.1442 | 100.0000 | 0.1400 | -0.0481 | 0.3364 |
| topN_sensitivity | HBV_INJURY_TOP_500 | 500.0000 | 500.0000 | proliferation_adjusted | 0.1379 | 95.6884 | 0.2656 | -0.1064 | 0.3823 |
| topN_sensitivity | HBV_INJURY_TOP_500 | 500.0000 | 500.0000 | proliferation_cibersortx_pc_adjusted | 0.2259 | 156.6777 | 0.0408 | 0.0097 | 0.4420 |
| topN_sensitivity | HBV_INJURY_TOP_500 | 500.0000 | 500.0000 | proliferation_selected_fraction_adjusted | 0.2209 | 153.2466 | 0.0326 | 0.0187 | 0.4231 |
| topN_sensitivity | HBV_INJURY_TOP_1000 | 1000.0000 | 1000.0000 | unadjusted | 0.1980 | 100.0000 | 0.0236 | 0.0271 | 0.3689 |
| topN_sensitivity | HBV_INJURY_TOP_1000 | 1000.0000 | 1000.0000 | proliferation_adjusted | 0.1395 | 70.4364 | 0.2003 | -0.0751 | 0.3541 |
| topN_sensitivity | HBV_INJURY_TOP_1000 | 1000.0000 | 1000.0000 | proliferation_cibersortx_pc_adjusted | 0.2158 | 108.9615 | 0.0267 | 0.0254 | 0.4061 |
| topN_sensitivity | HBV_INJURY_TOP_1000 | 1000.0000 | 1000.0000 | proliferation_selected_fraction_adjusted | 0.2110 | 106.5652 | 0.0200 | 0.0339 | 0.3881 |
| compact_primary_candidate | HBV_INJURY_TOP_2000 | 2000.0000 | 2000.0000 | unadjusted | 0.2370 | 100.0000 | 0.0020 | 0.0886 | 0.3855 |
| compact_primary_candidate | HBV_INJURY_TOP_2000 | 2000.0000 | 2000.0000 | proliferation_adjusted | 0.1304 | 55.0108 | 0.1534 | -0.0494 | 0.3102 |
| compact_primary_candidate | HBV_INJURY_TOP_2000 | 2000.0000 | 2000.0000 | proliferation_cibersortx_pc_adjusted | 0.1902 | 80.2346 | 0.0220 | 0.0281 | 0.3523 |
| compact_primary_candidate | HBV_INJURY_TOP_2000 | 2000.0000 | 2000.0000 | proliferation_selected_fraction_adjusted | 0.1881 | 79.3723 | 0.0144 | 0.0383 | 0.3380 |
| topN_sensitivity | HBV_INJURY_TOP_5000 | 5000.0000 | 5000.0000 | unadjusted | 0.2635 | 100.0000 | 1.420e-06 | 0.1614 | 0.3656 |
| topN_sensitivity | HBV_INJURY_TOP_5000 | 5000.0000 | 5000.0000 | proliferation_adjusted | 0.1188 | 45.0810 | 0.0465 | 0.0019 | 0.2357 |
| topN_sensitivity | HBV_INJURY_TOP_5000 | 5000.0000 | 5000.0000 | proliferation_cibersortx_pc_adjusted | 0.1531 | 58.1147 | 0.0063 | 0.0442 | 0.2621 |
| topN_sensitivity | HBV_INJURY_TOP_5000 | 5000.0000 | 5000.0000 | proliferation_selected_fraction_adjusted | 0.1486 | 56.3876 | 0.0043 | 0.0478 | 0.2494 |
| extended_sensitivity | HBV_INJURY_EXTENDED_7792 | 7792.0000 | 7792.0000 | unadjusted | 0.2328 | 100.0000 | 5.923e-09 | 0.1600 | 0.3056 |
| extended_sensitivity | HBV_INJURY_EXTENDED_7792 | 7792.0000 | 7792.0000 | proliferation_adjusted | 0.1250 | 53.6879 | 0.0041 | 0.0406 | 0.2094 |
| extended_sensitivity | HBV_INJURY_EXTENDED_7792 | 7792.0000 | 7792.0000 | proliferation_cibersortx_pc_adjusted | 0.1475 | 63.3795 | 4.995e-04 | 0.0662 | 0.2289 |
| extended_sensitivity | HBV_INJURY_EXTENDED_7792 | 7792.0000 | 7792.0000 | proliferation_selected_fraction_adjusted | 0.1334 | 57.3170 | 5.182e-04 | 0.0597 | 0.2071 |

**Supplementary Table S9. CIBERSORTx immune-fraction summaries.** Summary of CIBERSORTx-inferred immune-cell fractions in GSE121248. The table reports sample-level or group-level inferred immune fractions, tissue label, and CIBERSORTx-derived principal components used as immune-composition covariates in the HBV injury-axis regression models. These data support the CIBERSORTx adjustment and immune-composition supplementary figure.

| tissue | cell fraction | n | mean | median | sd |
| --- | --- | --- | --- | --- | --- |
| non_tumor | B cells memory | 37.0000 | 0.0212 | 0.0016 | 0.0296 |
| non_tumor | B cells naive | 37.0000 | 0.0166 | 0.0099 | 0.0223 |
| non_tumor | Dendritic cells activated | 37.0000 | 0.0033 | 0.0000 | 0.0072 |
| non_tumor | Dendritic cells resting | 37.0000 | 0.0121 | 0.0117 | 0.0114 |
| non_tumor | Eosinophils | 37.0000 | 1.951e-04 | 0.0000 | 8.825e-04 |
| non_tumor | Macrophages M0 | 37.0000 | 0.0062 | 0.0000 | 0.0235 |
| non_tumor | Macrophages M1 | 37.0000 | 0.0739 | 0.0595 | 0.0428 |
| non_tumor | Macrophages M2 | 37.0000 | 0.2100 | 0.2133 | 0.0668 |
| non_tumor | Mast cells activated | 37.0000 | 0.0560 | 0.0581 | 0.0364 |
| non_tumor | Mast cells resting | 37.0000 | 0.0040 | 0.0000 | 0.0106 |
| non_tumor | Monocytes | 37.0000 | 0.0353 | 0.0253 | 0.0317 |
| non_tumor | NK cells activated | 37.0000 | 0.0545 | 0.0503 | 0.0276 |
| non_tumor | NK cells resting | 37.0000 | 0.0078 | 0.0000 | 0.0176 |
| non_tumor | Neutrophils | 37.0000 | 0.0268 | 0.0271 | 0.0174 |
| non_tumor | Plasma cells | 37.0000 | 0.1794 | 0.1678 | 0.0563 |
| non_tumor | T cells CD4 memory activated | 37.0000 | 0.0000 | 0.0000 | 0.0000 |
| non_tumor | T cells CD4 memory resting | 37.0000 | 0.1203 | 0.1265 | 0.0616 |
| non_tumor | T cells CD4 naive | 37.0000 | 0.0000 | 0.0000 | 0.0000 |
| non_tumor | T cells CD8 | 37.0000 | 0.1094 | 0.1199 | 0.0420 |
| non_tumor | T cells follicular helper | 37.0000 | 0.0428 | 0.0430 | 0.0316 |
| non_tumor | T cells gamma delta | 37.0000 | 0.0117 | 0.0000 | 0.0200 |
| non_tumor | T cells regulatory (Tregs) | 37.0000 | 0.0085 | 0.0000 | 0.0117 |
| tumor | B cells memory | 70.0000 | 0.0253 | 3.486e-04 | 0.0373 |
| tumor | B cells naive | 70.0000 | 0.0212 | 0.0089 | 0.0267 |
| tumor | Dendritic cells activated | 70.0000 | 0.0108 | 0.0000 | 0.0182 |
| tumor | Dendritic cells resting | 70.0000 | 0.0133 | 0.0073 | 0.0187 |
| tumor | Eosinophils | 70.0000 | 4.197e-04 | 0.0000 | 0.0023 |
| tumor | Macrophages M0 | 70.0000 | 0.0481 | 0.0203 | 0.0719 |
| tumor | Macrophages M1 | 70.0000 | 0.0619 | 0.0553 | 0.0447 |
| tumor | Macrophages M2 | 70.0000 | 0.2089 | 0.2056 | 0.1027 |
| tumor | Mast cells activated | 70.0000 | 0.0286 | 0.0143 | 0.0359 |
| tumor | Mast cells resting | 70.0000 | 0.0170 | 0.0000 | 0.0318 |
| tumor | Monocytes | 70.0000 | 0.0366 | 0.0320 | 0.0302 |
| tumor | NK cells activated | 70.0000 | 0.0584 | 0.0633 | 0.0328 |
| tumor | NK cells resting | 70.0000 | 0.0114 | 0.0000 | 0.0256 |
| tumor | Neutrophils | 70.0000 | 0.0235 | 0.0137 | 0.0290 |
| tumor | Plasma cells | 70.0000 | 0.0897 | 0.0708 | 0.0524 |
| tumor | T cells CD4 memory activated | 70.0000 | 2.872e-05 | 0.0000 | 2.403e-04 |
| tumor | T cells CD4 memory resting | 70.0000 | 0.1673 | 0.1449 | 0.1051 |
| tumor | T cells CD4 naive | 70.0000 | 4.040e-04 | 0.0000 | 0.0034 |
| tumor | T cells CD8 | 70.0000 | 0.0989 | 0.0954 | 0.0568 |
| tumor | T cells follicular helper | 70.0000 | 0.0482 | 0.0428 | 0.0439 |
| tumor | T cells gamma delta | 70.0000 | 0.0016 | 0.0000 | 0.0046 |
| tumor | T cells regulatory (Tregs) | 70.0000 | 0.0287 | 0.0209 | 0.0302 |

**Supplementary Table S10. TCGA-LIHC Cox model full output.** Complete Cox proportional-hazards model output for TCGA-LIHC module-score survival analyses. The table reports model type, score, model term, hazard ratio per one-standard-deviation increase, 95% confidence interval, p-value, sample size, number of events, and model formula. Models include score-only, age/sex-adjusted, and age/sex/pathologic-stage-adjusted specifications.

| score | model | formula | n | events | term | estimate | std error | statistic | p value | conf low | conf high |
| --- | --- | --- | --- | --- | --- | --- | --- | --- | --- | --- | --- |
| ProlifHubScore | score_only | Surv(time_months, event) ~ scale(ProlifHubScore) | 369.0000 | 130.0000 | scale(ProlifHubScore) | 1.4393 | 0.0926 | 3.9309 | 8.462e-05 | 1.2003 | 1.7259 |
| HepLossScore | score_only | Surv(time_months, event) ~ scale(HepLossScore) | 369.0000 | 130.0000 | scale(HepLossScore) | 0.8630 | 0.0974 | -1.5117 | 0.1306 | 0.7130 | 1.0446 |
| HCCStateScore | score_only | Surv(time_months, event) ~ scale(HCCStateScore) | 369.0000 | 130.0000 | scale(HCCStateScore) | 1.4277 | 0.1024 | 3.4771 | 5.068e-04 | 1.1681 | 1.7451 |
| ProlifHubScore | age_sex_adjusted | Surv(time_months, event) ~ scale(ProlifHubScore) + age_tmp + gender_tmp | 369.0000 | 130.0000 | scale(ProlifHubScore) | 1.4852 | 0.0956 | 4.1379 | 3.506e-05 | 1.2314 | 1.7912 |
| ProlifHubScore | age_sex_adjusted | Surv(time_months, event) ~ scale(ProlifHubScore) + age_tmp + gender_tmp | 369.0000 | 130.0000 | age_tmp | 1.0154 | 0.0075 | 2.0304 | 0.0423 | 1.0005 | 1.0305 |
| ProlifHubScore | age_sex_adjusted | Surv(time_months, event) ~ scale(ProlifHubScore) + age_tmp + gender_tmp | 369.0000 | 130.0000 | gender_tmpmale | 0.9096 | 0.1871 | -0.5063 | 0.6126 | 0.6304 | 1.3126 |
| HepLossScore | age_sex_adjusted | Surv(time_months, event) ~ scale(HepLossScore) + age_tmp + gender_tmp | 369.0000 | 130.0000 | scale(HepLossScore) | 0.8655 | 0.0979 | -1.4743 | 0.1404 | 0.7144 | 1.0487 |
| HepLossScore | age_sex_adjusted | Surv(time_months, event) ~ scale(HepLossScore) + age_tmp + gender_tmp | 369.0000 | 130.0000 | age_tmp | 1.0089 | 0.0071 | 1.2421 | 0.2142 | 0.9949 | 1.0231 |
| HepLossScore | age_sex_adjusted | Surv(time_months, event) ~ scale(HepLossScore) + age_tmp + gender_tmp | 369.0000 | 130.0000 | gender_tmpmale | 0.8151 | 0.1859 | -1.0996 | 0.2715 | 0.5662 | 1.1735 |
| HCCStateScore | age_sex_adjusted | Surv(time_months, event) ~ scale(HCCStateScore) + age_tmp + gender_tmp | 369.0000 | 130.0000 | scale(HCCStateScore) | 1.4479 | 0.1044 | 3.5452 | 3.923e-04 | 1.1800 | 1.7766 |
| HCCStateScore | age_sex_adjusted | Surv(time_months, event) ~ scale(HCCStateScore) + age_tmp + gender_tmp | 369.0000 | 130.0000 | age_tmp | 1.0119 | 0.0073 | 1.6246 | 0.1042 | 0.9976 | 1.0265 |
| HCCStateScore | age_sex_adjusted | Surv(time_months, event) ~ scale(HCCStateScore) + age_tmp + gender_tmp | 369.0000 | 130.0000 | gender_tmpmale | 0.8539 | 0.1857 | -0.8501 | 0.3953 | 0.5934 | 1.2289 |
| ProlifHubScore | age_sex_stage_adjusted | Surv(time_months, event) ~ scale(ProlifHubScore) + age_tmp + gender_tmp + stage_collapsed | 173.0000 | 74.0000 | scale(ProlifHubScore) | 1.4392 | 0.1237 | 2.9439 | 0.0032 | 1.1294 | 1.8339 |
| ProlifHubScore | age_sex_stage_adjusted | Surv(time_months, event) ~ scale(ProlifHubScore) + age_tmp + gender_tmp + stage_collapsed | 173.0000 | 74.0000 | age_tmp | 1.0017 | 0.0090 | 0.1905 | 0.8489 | 0.9841 | 1.0196 |
| ProlifHubScore | age_sex_stage_adjusted | Surv(time_months, event) ~ scale(ProlifHubScore) + age_tmp + gender_tmp + stage_collapsed | 173.0000 | 74.0000 | gender_tmpmale | 1.2735 | 0.2623 | 0.9216 | 0.3567 | 0.7615 | 2.1297 |
| ProlifHubScore | age_sex_stage_adjusted | Surv(time_months, event) ~ scale(ProlifHubScore) + age_tmp + gender_tmp + stage_collapsed | 173.0000 | 74.0000 | stage_collapsedII |  | 0.0000 |  |  |  |  |
| ProlifHubScore | age_sex_stage_adjusted | Surv(time_months, event) ~ scale(ProlifHubScore) + age_tmp + gender_tmp + stage_collapsed | 173.0000 | 74.0000 | stage_collapsedIII |  | 0.0000 |  |  |  |  |
| ProlifHubScore | age_sex_stage_adjusted | Surv(time_months, event) ~ scale(ProlifHubScore) + age_tmp + gender_tmp + stage_collapsed | 173.0000 | 74.0000 | stage_collapsedIV | 3.6897 | 0.6212 | 2.1017 | 0.0356 | 1.0921 | 12.4660 |
| HepLossScore | age_sex_stage_adjusted | Surv(time_months, event) ~ scale(HepLossScore) + age_tmp + gender_tmp + stage_collapsed | 173.0000 | 74.0000 | scale(HepLossScore) | 0.8820 | 0.1473 | -0.8526 | 0.3939 | 0.6608 | 1.1772 |
| HepLossScore | age_sex_stage_adjusted | Surv(time_months, event) ~ scale(HepLossScore) + age_tmp + gender_tmp + stage_collapsed | 173.0000 | 74.0000 | age_tmp | 0.9962 | 0.0084 | -0.4548 | 0.6493 | 0.9799 | 1.0127 |
| HepLossScore | age_sex_stage_adjusted | Surv(time_months, event) ~ scale(HepLossScore) + age_tmp + gender_tmp + stage_collapsed | 173.0000 | 74.0000 | gender_tmpmale | 1.1333 | 0.2552 | 0.4901 | 0.6240 | 0.6872 | 1.8689 |
| HepLossScore | age_sex_stage_adjusted | Surv(time_months, event) ~ scale(HepLossScore) + age_tmp + gender_tmp + stage_collapsed | 173.0000 | 74.0000 | stage_collapsedII |  | 0.0000 |  |  |  |  |
| HepLossScore | age_sex_stage_adjusted | Surv(time_months, event) ~ scale(HepLossScore) + age_tmp + gender_tmp + stage_collapsed | 173.0000 | 74.0000 | stage_collapsedIII |  | 0.0000 |  |  |  |  |
| HepLossScore | age_sex_stage_adjusted | Surv(time_months, event) ~ scale(HepLossScore) + age_tmp + gender_tmp + stage_collapsed | 173.0000 | 74.0000 | stage_collapsedIV | 2.7342 | 0.6089 | 1.6519 | 0.0986 | 0.8290 | 9.0185 |
| HCCStateScore | age_sex_stage_adjusted | Surv(time_months, event) ~ scale(HCCStateScore) + age_tmp + gender_tmp + stage_collapsed | 173.0000 | 74.0000 | scale(HCCStateScore) | 1.5075 | 0.1506 | 2.7258 | 0.0064 | 1.1222 | 2.0251 |
| HCCStateScore | age_sex_stage_adjusted | Surv(time_months, event) ~ scale(HCCStateScore) + age_tmp + gender_tmp + stage_collapsed | 173.0000 | 74.0000 | age_tmp | 1.0001 | 0.0088 | 0.0058 | 0.9953 | 0.9830 | 1.0174 |
| HCCStateScore | age_sex_stage_adjusted | Surv(time_months, event) ~ scale(HCCStateScore) + age_tmp + gender_tmp + stage_collapsed | 173.0000 | 74.0000 | gender_tmpmale | 1.2360 | 0.2600 | 0.8149 | 0.4151 | 0.7425 | 2.0576 |
| HCCStateScore | age_sex_stage_adjusted | Surv(time_months, event) ~ scale(HCCStateScore) + age_tmp + gender_tmp + stage_collapsed | 173.0000 | 74.0000 | stage_collapsedII |  | 0.0000 |  |  |  |  |
| HCCStateScore | age_sex_stage_adjusted | Surv(time_months, event) ~ scale(HCCStateScore) + age_tmp + gender_tmp + stage_collapsed | 173.0000 | 74.0000 | stage_collapsedIII |  | 0.0000 |  |  |  |  |
| HCCStateScore | age_sex_stage_adjusted | Surv(time_months, event) ~ scale(HCCStateScore) + age_tmp + gender_tmp + stage_collapsed | 173.0000 | 74.0000 | stage_collapsedIV | 3.3633 | 0.6165 | 1.9675 | 0.0491 | 1.0047 | 11.2594 |
